# Emergent frequency-dependent selection predicts mutation outcomes in complex ecological communities

**DOI:** 10.64898/2026.04.13.718251

**Authors:** Shing Yan Li, Zhijie Feng, Akshit Goyal, Pankaj Mehta

## Abstract

Ecological interactions can dramatically alter evolutionary outcomes in complex communities. Yet, most population genetics models neglect how interactions within complex communities shape evolutionary dynamics. Here, we bridge this gap by using dynamical mean-field theory to integrate community ecology into classical population genetics models. We show that near an ecological steady state, interactions generate emergent negative frequency-dependent selection between parents and phenotypically similar mutants. Strikingly, even in species-rich communities, this frequency-dependent selection can be characterized by a single parameter measuring the strength of ecological feedback. We find that these interactions strongly suppress the fixation of moderately beneficial mutations. This suppression arises because frequency-dependent selection promotes prolonged co-existence between parent and mutant lineages, creating a barrier to fixation. These effects strengthen with larger effective population sizes and more open niches in the community. Applying our theory to human gut microbiome data, we estimate the average strength of ecological feedback and find it consistent with the negative frequency dependence our theory predicts. Our study establishes a framework for integrating ecological feedback from complex communities into population genetics.

## INTRODUCTION

A major goal of biology is to develop a quantitative theory of evolution. The mathematical framework of population genetics was a foundational step toward this goal. A cornerstone of population genetics is Kimura’s diffusion model (also called Wright-Fisher diffusion) [1– 5], which conceptualizes the evolutionary dynamics of a mutant’s frequency as a stochastic process, subject to both diffusion (random genetic drift) and deterministic selection. Using this theory, one can predict two fundamental aspects of a mutant’s fate: its fixation probability and the time it takes to either fix in the population or go extinct.

Since then, it has been shown that a large class of evolutionary dynamics map onto Kimura’s diffusion model. Examples range from Wright-Fisher and Moran processes [4, 6] to single-species serial dilution experiments like Lenski’s long-term evolution [7, 8]. These developments have led to the widespread view that the results of classical population genetics apply broadly to evolving populations [4, 9, 10]. Indeed, subsequent studies have used Kimura’s basic framework as a starting point while extending evolution to other biological contexts, e.g., in fluctuating environments [11], and with spatial structure [12, 13].

While successful, these models make a simplifying assumption: that the evolutionary dynamics of mutants are mainly determined by their interaction with their parents. In doing so, they incorporate ecological context implicitly rather than explicitly. This makes it hard to understand how ecological interactions will alter evolutionary trajectories [14–16] which is important since almost all evolving populations are embedded in complex ecological communities with which they interact [17–19]. Any comprehensive theory of evolution must therefore explain how ecological interactions with a complex community will affect mutation outcomes. Without such an eco-evolutionary perspective, our understanding of evolution in natural contexts will remain incomplete [20–22].

Numerous works have sought to bridge this gap. These include population genetics models with frequency-dependent selection that capture more realistic evolutionary scenarios [23–28]. However, since these models assume communities with one to two species [29–31], it is unclear whether these apply to more complex ecological settings [32, 33]. Other evolutionary theories seek to integrate ecological context [34, 35] but many of these efforts remain qualitative in nature and give rise to very different predictions [36–41]. For example, experimental studies have shown that communities may promote [36, 37] or constrain [40, 41] adaptation depending on the specific ecological setting.

Over the past decade, quantitative approaches have made significant progress understanding the role of ecological interactions in shaping mutant evolutionary trajectories. However, several of these works often make specific assumptions that make it hard to assess their general applicability, such as low community diversity [42] and specific fitness functions [43]. Some of these studies have used models of trait evolution [44, 45] and pathogen competition [46–49], and found that ecological interactions can generate effective frequency-dependent selection. Others have used consumer-resource models to explain evolutionary outcomes such as diversification [50]and functional robustness [51]. Yet others [14, 15] have focused on predicting mutation outcomes in complex communities. However, these studies have largely focused on deterministic dynamics and neglected stochastic dynamics, which are crucial for new mutant lineages which start at extremely low population sizes (often a single individual). Hence, despite these advances, a major outstanding challenge remains to develop a theoretical framework to predict the stochastic dynamics of mutants in highly-diverse communities that is general enough to apply across a wide variety of ecological models.

At first, incorporating all the complex details of community ecological interactions might seem intractable due to the large number of unknown parameters. However, recent approaches from statistical physics and random matrix theory have revealed that for a sufficiently complex ecological community, it is possible to gain useful theoretical insights about ecological properties of an ecosystem such as its diversity or stability [52–59]. More recent works have started to adapt these methods to ask about short-term evolutionary processes and ecological invasions [14, 15]. A key insight from these works has been the realization that at the community level, the effects of ecological interactions can often be summarized in terms of a few emergent parameters that relate to susceptibilities that measure the strength of ecological feedback [60]. These efforts suggest it should be possible to use tools from statistical physics to incorporate between ecological interactions into classic population genetics models.

Inspired by these works, here we derive an effective theory of population genetics in complex ecological communities near a steady state. We use dynamical meanfield theory (DMFT) to coarse-grain the effect of a highly-diverse, randomly assembled community on a mutant’s frequency dynamics. Our central result is that community-mediated feedbacks generically lead to emergent frequency-dependent selection across a variety of ecological models. Importantly, a community’s combined effect can be captured by only a single new parameter beyond the mutant’s fitness that quantifies the strength of community-mediated feedbacks on mutant dynamics. This parameter is tightly related to the community’s statistical properties such as its niche-packing. We find that community effects dramatically alter the fixation probabilities of mutants compared to Kimura’s classical predictions derived in the absence of frequency dependent selection. Further, in diverse communities, mutants can often coexist with their parents for extremely long periods, which drastically alters their fixation or extinction times. Our results highlight that the fate of a mutant can be quantitatively predicted by population genetics models even in complex ecological settings, and provide a path to estimate ecological feedbacks in a complex community from mutant frequency dynamics.

## RESULTS

### Ecological feedbacks manifest as emergent frequency-dependent selection

Population genetics aims to describe how mutations spread through a species population. To do so, it follows the dynamics of a mutant’s frequency *f*(*t*) in the population given by

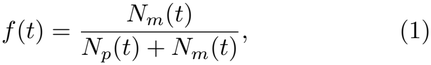

where *N*_*p*_(*t*) and *N*_*m*_(*t*) are the abundances of parent and mutant strains at time *t*, respectively (Fig. 1a and c). For convenience, throughout this work, we use the term “classical population genetics” synonymously with the single-locus diffusion limit of population genetics in hap-loid populations with constant selection [1, 2, 4]. Such models assume that mutant dynamics are shaped by two major evolutionary forces: (i) selection, which is characterized by a selection coefficient *s* that measures the fitness difference between the mutant and parent, and (ii) stochastic drift, which is parameterized in terms of an effective population size *N*_eff_ (Fig. 1d). Stochastic effects play an important role in mutational dynamics because mutants typically emerge at low frequencies *f*(0) ≪1, making their early dynamics inherently noisy, e.g., due to randomness in birth and death events [4, 61]. At long times, mutants either fix (*f* = 1) or go extinct (*f* = 0, Fig. 1f and h)

Nevertheless, by treating the selection coefficient *s* as a constant (Fig. 1d), these models implicitly assume that ecological interactions do not change the selective landscape over the course of mutant dynamics. For this reason, they often neglect the dynamic environmental feedbacks through which ecological communities can potentially alter mutant evolutionary trajectories (Fig. 1b) [17–19]. This assumption becomes especially hard to justify when communities are highly-diverse, from rain-forests to microbiomes.

**FIG. 1:**
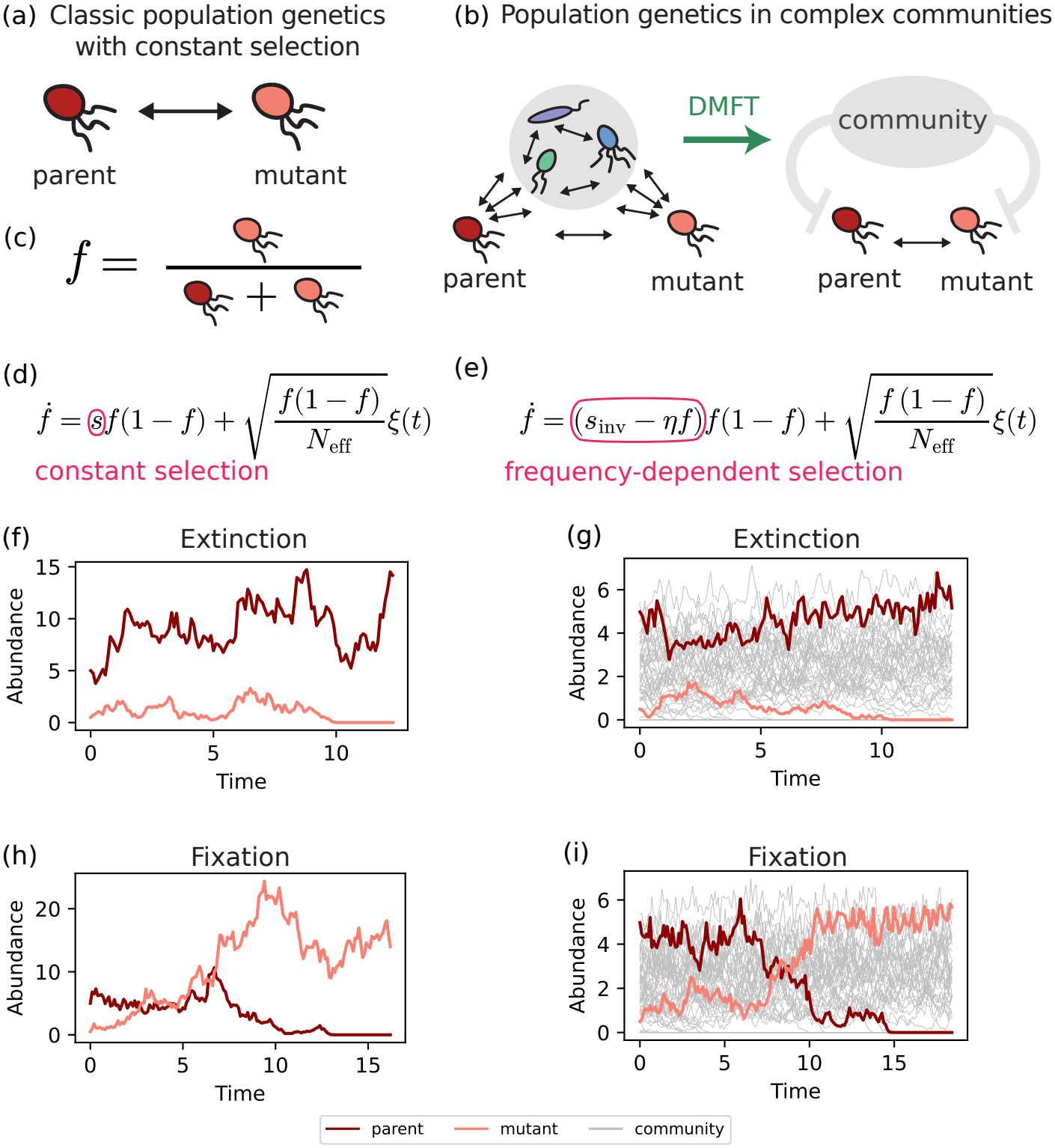
Population genetics and parent-mutant dynamics in the absence and presence of complex communities. (a) Classic population genetics, with constant selection, typically models dynamics only for the parent and the mutant, independent of community context. (b) In complex communities, ecological interactions among species collectively influence the parent-mutant dynamics. As shown using dynamical mean-field theory, the ecological community mediates an emergent frequency-depdendent selection between the parent and the mutant. (c) Parent-mutant dynamics can be expressed in terms of mutant frequency *f* . (d) In classical population genetics, the dynamics of *f* are driven by constant selection and stochastic drift. (e) In complex communities, additional frequency-dependent selection arises, with an emergent parameter *η* characterizing the strength of ecological feedbacks. (f-i) Examples of dynamics of species abundances in classical population genetics (left) versus in complex communities (other species’ dynamics in grey). After a long time, the mutant eventually reaches either extinction (f–g) or fixation (h–i).

In this work we develop a general framework to understand how ecology affects mutant dynamics. Consistent with Refs. [15, 50], we find that when mutations arise in a complex ecological community, the selection coefficient *s* can no longer be treated as a constant or simple time-dependent function [11]. Instead, for communities near a steady state, selection becomes frequency-dependent. Importantly, this frequency dependence emerges as a consequence of community-mediated ecological feedbacks on the parent and mutant (Fig. 1b), which can be captured with just a few parameters.

To illustrate these ideas, we begin by analyzing the evolutionary dynamics of a mutant in an ecosystem modeled using a generalized Lotka-Volterra model (GLV) with demographic noise. The GLV model describes a complex, highly-diverse community of *S*≫ 1 species with abundances *N*_*i*_ (*i* = 1, …, *S*) whose dynamics are governed by stochastic differential equations of the form

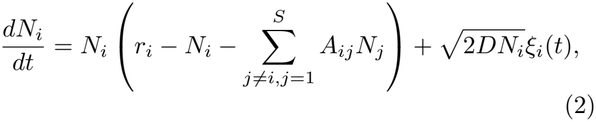

with the *ξ*_*i*_(*t*) independent normal random variables with ⟨ *ξ*_*i*_(*t*) ⟩ = 0 and ⟨ *ξ*_*i*_(*t*)*ξ*_*j*_(*t*^*′*^) ⟩= *δ*_*ij*_*δ*(*t* −*t*^*′*^). In this expression, *r*_*i*_ is the intrinsic growth rates of species *i*; *A*_*ij*_ captures ecological interactions between species and encodes how species *j* affects the abundance of species *i*; and 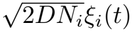 represents demographic noise. The parameter *D* sets the overall scale of demographic noise and serves as an analog of a temperature or diffusion coefficient. Following recent work, we model parameters *r*_*i*_ and *A*_*ij*_ as random variables [54, 57, 62] drawn from Gaussian distributions (Methods), although our results remain valid for a wide variety of other distributions, as long as they have finite mean and variance [63, 64] and negligible higher cumulants [65] (SI Sec. A).

To study population genetics in the GLV, we follow [15, 56, 59] and first initialize a community with *S* randomly chosen species at steady-state (Methods). We then pick one of these species at random to be the parent *p*. We denote the abundance of the parent species in the community by *N*_*p*_(*t*). At time *t* = 0, we introduce a mutant *m* to this community at a small abundance *N*_*m*_(0) ≪*N*_*p*_(0). This is equivalent to assuming that the mutant starts at low frequency *f*_0_ ≪ 1. To model the ecological similarity of parent and mutant, we assume that the mutant’s interaction parameters, *A*_*mj*_, with other species in the community are correlated with those of the parent, *A*_*pj*_. We denote the strength of this correlation as *ρ*, which is independent of any difference in intrinsic growth rates between parent and mutant.

Crucially, we assume that mutants and parents are highly correlated (*ρ* ≈1) in their interactions with the community. With a high *ρ*, the total parent and mutant population behaves effectively as a single species from the community perspective, which greatly simplifies the mutant dynamics by suppressing various community effects (SI Sec. A, F). Moreover, mutants and parents interact much more strongly with each other than with any other species in the community. To model this, we set parent-mutant interactions *A*_*mp*_, *A*_*pm*_ to be close to but weaker than the intraspecific competition (≈1, SI Sec. A), while interactions with all other species in the community are assumed to be much weaker (≈1*/S*, SI Sec. A). However, as we will show below, even though mutants interact weakly with any individual species in the community, these weak effects add up due to the large number of species (≈*S*) present in the community. Thus as a collective, community interactions can no longer be neglected and can strongly influence mutant dynamics. Throughout this work, we focus on the strong-selection-weak-mutation regime [66]: no subsequent mutations of the same species occur before a mutant fixes or goes extinct (Discussion). For this reason, we focus on a single mutation and ignore the scenarios of multiple mutants of the same species (known as clonal interference) or the role of mutation rates. We also assume that the community is at an ecological steady state when the mutant arises.

Our goal is to derive the effective dynamics of the mutant frequency *f*(*t*) in a complex steady community described by the GLV. To do so, we use dynamical meanfield theory (DMFT) [52, 53, 56, 67]. The key idea behind DMFT is that in large, diverse communities the net effect of the community can be coarse-grained into an effective feedback on parent-mutant dynamics which is encoded in a single emergent parameter *η* beyond fitness that summarizes the strength of ecological feedbacks (Fig. 1b). Using this technique, we find that mutant frequency can be described using the equation

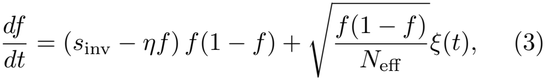

where *s*_inv_ measures the invasion fitness of the mutant in the community. Since *s*_inv_ contains both the intrinsic selection coefficient *s* (e.g., differences in intrinsic growth rates) and community effects (SI Sec. A), it plays a role analogous to the mutant’s selective coefficient in classical population genetics. *N*_eff_ = ⟨*N*_*p*_(0) + *N*_*m*_(0)⟩ */*2*D* measures the effective population size of the combined parent-mutant population, which is approximately constant on average, especially for mutants where *ρ*≈ 1 (SI Sec. A). *ξ*(*t*) is a random normal variable with ⟨*ξ*(*t*) ⟩ = 0 and ⟨*ξ*(*t*)*ξ*(*t*^*′*^)⟩ = *δ*(*t* −*t*^*′*^).

This equation results from two distinct eco-evolutionary processes: deterministic selection – the term proportional to *f*(1 −*f*) – and stochastic drift– the term proportional to *ξ*(*t*). Remarkably, once the (invasion) fitness is known, just introducing a single new parameter *η* suffices to capture the collective effect of the entire community on parent-mutant dynamics. Eq. (3) shows that selection is now frequency-dependent since the selection coefficient *s*_inv_ −*ηf* changes with mutant frequency. When *η* = 0, e.g., in the absence of a community, we recover classical population genetics with a constant selection coefficient. However, when *η≠*0, ecological feedbacks fundamentally alter evolutionary dynamics.

We show that community-mediated feedbacks can manifest as frequency-dependent selection across a wide variety of ecological models. In the SI, we derive expressions for the effective parameters—*s*_inv_, *N*_eff_ and *η*—for MacArthur Consumer-Resource Models (CRMs) [68, 69] and generalized CRMs with nonlinear per-capita growth rates, which includes models with cross-feeding [70] (SI Sec. B and C). Remarkably, despite the diverse mathematical structures of these models, they all reduce to the same form of frequency-dependent selection near ecological steady states. While it has been well known that ecological interactions with a few other species can lead to frequency-dependent selection (Introduction), our results show that the same type of frequency-dependent selection in Eq. (3) also emerges in communities with many species near a steady state. As a result, mutation outcomes can be predicted using simple models despite the ecological complexity of communities. Our results also generalize previous work [15] and predict the full stochastic trajectories of mutants, in addition to being applicable across a wide variety of ecological models.

### Generalization of Kimura’s formula for fixation probabilities

To test whether Eq. (3) indeed captures ecological effects, we now explore the consequences of community interactions on the fate of a mutant. To do so, we draw on existing results in population genetics on both constant and frequency-dependent selection [1, 23– 28, 71, 72]).

One of the celebrated results of classical population genetics is Kimura’s formula (with constant selection) for fixation probability *p*_fix_ [1], which describes how likely a mutant is to replace its parent and fix in a population (Fig. 2c). Kimura’s formula is given by

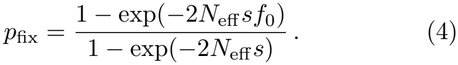

**FIG. 2:**
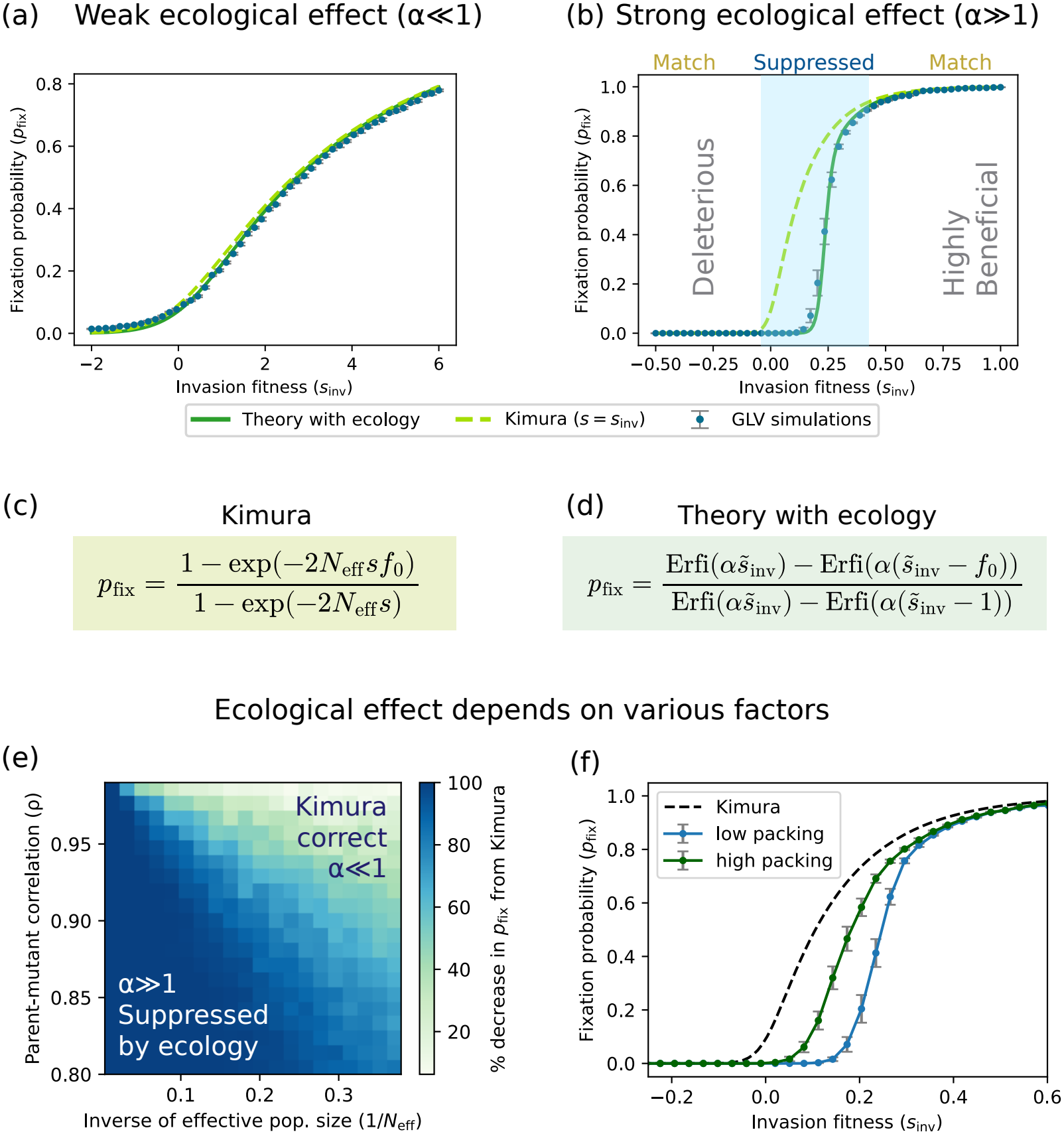
Fixation probabilities suppressed by strong ecological effects. We compute fixation probabilities *p*_fix_ at various invasion fitness *s*_inv_ by simulating mutations within generalized Lotka-Volterra models. Error bars denote standard errors from multiple instances of demographic noise and mutants. (a) When ecological effects are weak relative to stochastic drift (*α≪* 1), our theory matches Kimura’s formula with *s* = *s*_inv_ for fixation probabilities and predicts them accurately in simulations. (b) When ecological effects are strong (*α ≫*1), Kimura’s formula applies only for deleterious or highly beneficial mutants. For moderately beneficial mutants with *s*_inv_ ≲ *η/*2, fixation probabilities are strongly suppressed relative to Kimura’s prediction. Our theory with ecology can capture such suppression and matches simulations. We give analytic expressions for fixation probability in (c) Kimura’s theory and (d) our theory with ecology. Ecological suppression of mutants depends on several factors: (e) For mutations with *s*_inv_ = 0, we quantify the suppression by the percentage decrease in fixation probability from Kimura’s prediction (1 *−p*_fix_*/f*_0_). Suppression increases with larger effective population size *N*_eff_ and lower parent-mutant correlation *ρ*. (f) Suppression is also stronger when the community is less packed (has more “open niches”), i.e., the number of surviving species is low relative to the maximum number of species allowed. The extent of packing is controlled by different *σ*^2^, the variance of the interactions (SI Sec. J).

This formula shows that as the fitness difference between mutant and parent *s* increases, mutants become more likely to fix. Specifically, neutral mutants with *s* = 0 fix with a probability equal to their initial frequency *p*_fix_ = *f*_0_. When selection is weak, i.e., *N*_eff_ |*s*|≪ 1, mutants are “effectively neutral” and fix with the same probability *f*_0_. When selection is strong, i.e., *N*_eff_ |*s*| ≫ 1, the formula reduces to 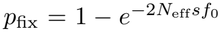 for positive *s* and 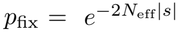 for negative *s*. Deleterious mutants with *s <* 0 thus have a negligible fixation probability, while for beneficial mutants with *s >* 0 the fixation probability increases linearly *p*_fix_ ∝ *s*. Ultimately, once a mutation is strongly beneficial (*s* ≫ 1*/N*_eff_*f*_0_) the mutant will almost surely fix since *p*_fix_ ≈1.

Starting with Eq. (3), one can solve the corresponding backward equation to obtain an analytic formula for the fixation probability *p*_fix_ in complex communities [1, 71] (see also SI Sec. D):

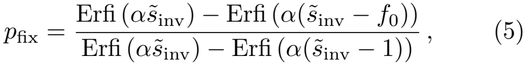

where

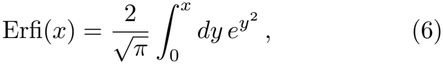

is the imaginary error function,

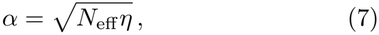

is the ratio of the strength of ecological feedbacks and stochastic drift and

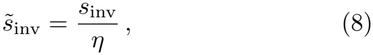

is the mutant invasion fitness normalized by the strength of the ecological feedback. The quantity 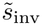 is closely related to the “dressed invasion fitness” introduced in [14] for predicting the outcomes of ecological invasions. In SI Sec. D, we elaborate on the similarities and differences between the two quantities. Eq. (5) serves as the generalization of Eq. (4) in the context of complex ecological communities.

To gain intuition for this formula, it is useful to look at various limits of this expression. When ecological effects are weak, i.e., *α*≪ 1, Eq. (5) reduces back to Kimura’s formula in Eq. (4) (SI Sec. D) with *s* = *s*_inv_. To numerically confirm this, we simulated the dynamics of complex communities governed by the GLV model into each of which we introduced a mutant of a randomly chosen parent strain. We made ecological effects weak simply by setting a large value of *D*, increasing the strength of stochastic drift. We repeated simulations for mutants with a given invasion fitness *s*_inv_ and measured the fraction of simulations in which mutants fixed (Methods). As shown in Fig. 2a, the simulated fixation probabilities match Eq. (5) remarkably well across a range of *s*_inv_. Further, in this limit, Eq. (5) virtually overlaps Kimura’s formula. In the next section, we will look at the opposite case of strong ecological effects, i.e., *α*≫ 1, and show that we get a qualitatively different picture of the fate of a mutant compared to Kimura’s formula.

### Ecological interactions strongly suppress mutant fixation

In the presence of strong ecological effects, i.e., when *α*≫ 1, our numerical simulations for the fixation probability show a stark deviation from Kimura’s predictions (Fig. 2b), with an almost switch-like behavior with increasing *s*_inv_. This is in contrast with Kimura’s formula which shows a gradual, linear increase in fixation probability over a range of *s*. The deviation from Kimura is most pronounced for moderately beneficial mutants where ecological effects strongly suppress the fixation probability (Fig. 2b, blue region). The origin of this suppression is easiest to understand for mutants with *s*_inv_ = 0 where Eq. (5) reduces to (SI Sec. D)

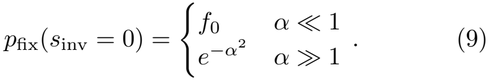

This formula agrees with Kimura’s results when ecological effects are weak (*α* ≪1) and shows that ecological effects exponentially suppress mutations with *s*_inv_ = 0 when ecological effects are large (*α* ≫ 1).

Using Eq. (5), we can also compute the full range of invasion fitness *s*_inv_ for which mutants are suppressed when ecological effects are strong (SI Sec. D). We find that suppression occurs when

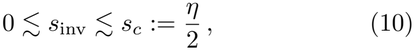

where *s*_*c*_ = *η/*2 is the critical invasion fitness at which fixation probability switches from roughly 0 to almost 1.As we show in SI Sec. D, in this case the dominant contribution to ecological suppression in the range 0 ≲ *s*_inv_ ≲ *s*_*c*_ comes from the last term in the denominator of Eq. (5).

The ecological suppression of mutants is an emergent community-mediated phenomenon whose strength depends on 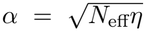. To understand how different ecological properties of the underlying community control the strength of suppression, we varied *α* by scanning over ranges of effective population sizes *N*_eff_ and parent-mutant correlation *ρ*, and plotted the corresponding deviation of the simulated fixation probability from Kimura’s predictions for a mutation with *s*_inv_ = 0 (Fig. 2e). Increasing *N*_eff_ further suppressed mutant fixation by increasing *α* (Fig. 2e). This can be understood by noting that increasing *N*_eff_ weakens stochastic drift, and hence increases the importance of ecological feedbacks. Suppression also increased with decreasing *ρ* (Fig. 2e). The reason for this is that the ecological feedback parameter *η* is proportional to 1 −*ρ* (SI Sec. A–C). Intuitively, as parent and mutant become less similar (i.e., *ρ* decreases), mutants interact less strongly with their parents, allowing community feedbacks to play a stronger role in the dynamics.

Most interestingly, suppression was stronger in less “niche-packed” communities (Fig. 2f), where we have defined packing as the ratio of surviving species *S*^*\**^ com-pared to largest number of species that can stably coexist in the community (often referred to as the species packing bound). In GLV, the maximum comes from May’s stability bound *S*^*\**^ ≲ 1*/σ*^2^ [55], where *σ*^2^ is the variance of the interactions *A*_*ij*_ (SI Sec. A); in CRM, it comes from competitive exclusion that *S*^*\**^ ≲ *M* ^*\**^, the number of non-depleted resources (SI Sec. B). This suppression suggests that ecological feedbacks become weaker as communities get closer to a fully packed regime where all niches are occupied, consistent with [15] (SI Sec. E). To understand this effect, we draw on an analogy between ecological communities and mechanical jamming [73]. Mechanical systems become harder to deform as they are packed with more particles, ultimately becoming rigid when they reach the jamming limit. Similarly, ecological communities become more “rigid” and less “deformable” as vacant niches get occupied due to a change in their susceptibility to perturbations. (Note that in consumer-resource models, the relevant susceptibility is in resource (niche) space, rather than species abundance space.) In support of this analogy, we have verified that it is really packing ratio that contributes to *η* by affecting community sus-ceptibility, rather than the biomass or species richness (SI Sec. A, B, E).

Finally, we show in the SI that the results in Fig. 2 generalize beyond the GLV model to many variants of consumer resource models. This leads us to conclude that ecological interactions likely generically suppress fixation of moderately beneficial mutants, strongly deviating from the predictions of classical population genetics.

### Parent-mutant coexistence alters fixation and extinction times

In addition to fixation probability, the second important quantity in population genetics is the time it takes for a mutant to reach fixation or extinction (which we refer to as absorption time). This quantity characterizes the timescale of adaptation in evolutionary dynamics. It is also informative to study the establishment time or equivalently the time to extinction. These two time scales are intimately related since extinction probability becomes negligible as soon as the mutant becomes established in the population, long before fixation. The mean extinction time can also be theoretically estimated (SI Sec. H). These times are relatively short in classic population genetics. Namely, they reach a maximum of order *N*_eff_ *f*_0_ |log *f*_0_| [74] for neutral mutations (*s* = 0), and decay with the magnitude of the fitness difference |*s*|.

On the other hand, we find that when ecological effects are strong (*α* ≫ 1), the mean times to absorption (Fig. 3a) and extinction (Fig. 3b) become exponentially large for moderately beneficial mutants (i.e., *s*_inv_ ≲ *η*). Both of these times show a sharp peak at the critical fitness difference *s*_inv_ ≃ *η/*2 (Eq. (10)). This increase in absorption time is so dramatic that it can even be directly observed in simulation trajectories of mutant and parent dynamics (see Fig. 3c–e). We further find that the maximum of both the absorption and extinction times grows exponentially with the effective population size *N*_eff_ . This is in contrast with Kimura’s theory where the maximum time grows linearly with *N*_eff_ . The presence of these exponentially long time scales raises the possibility that parents and mutants can transiently coexist for long periods in complex communities [14, 15], high-lighting a crucial difference between classical population genetics with constant selection and our eco-evolutionary approach.

**FIG. 3:**
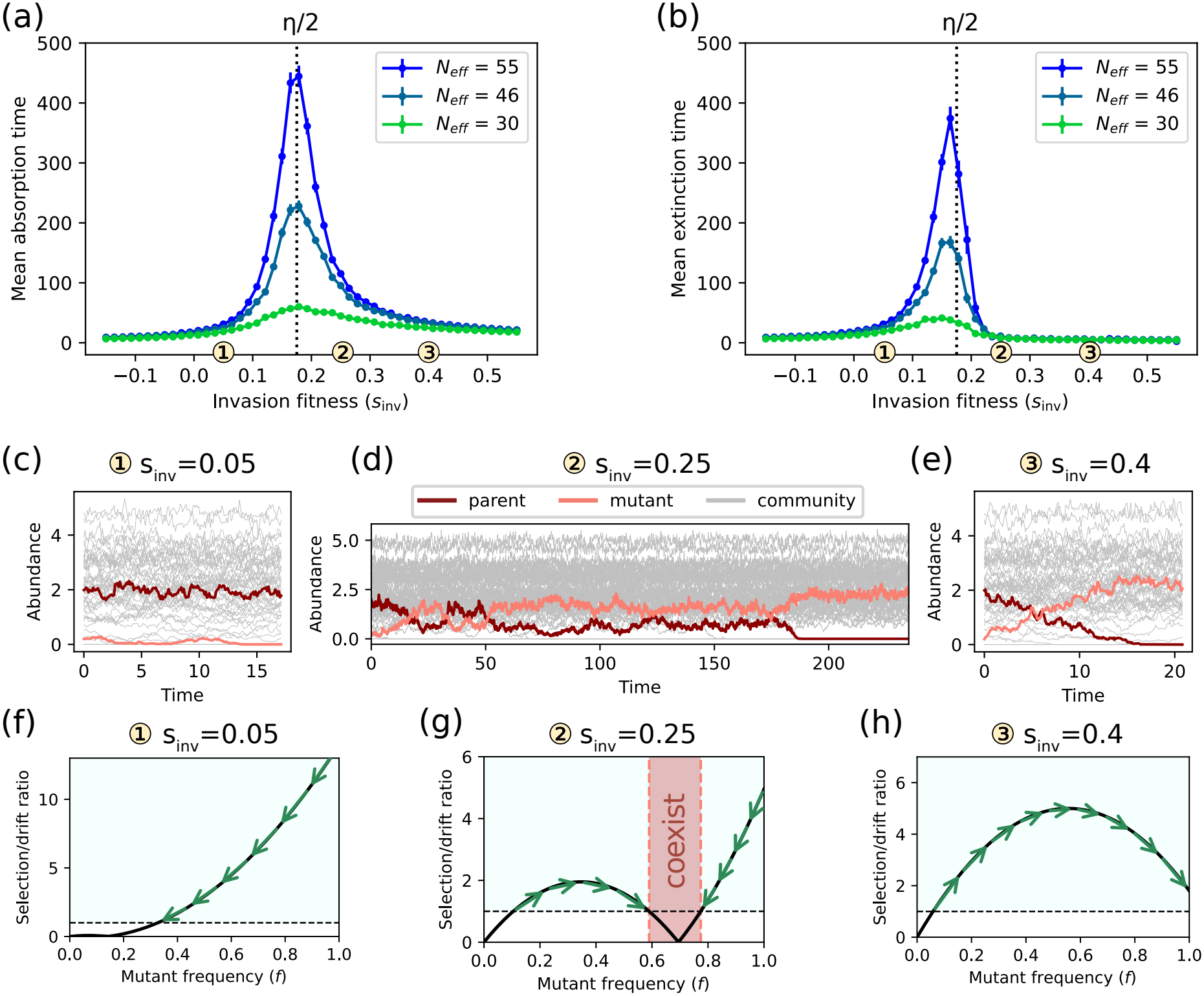
Parent-mutant coexistence in complex communities. We compute mean absorption and extinction times at various invasion fitness *s*_inv_ by simulating mutations within generalized Lotka-Volterra models. Error bars denote standard errors from multiple instances of demographic noise. (a, b) The time scales of parent-mutant dynamics at various invasion fitness *s*_inv_ reveal the possibility of coexistence. Under strong ecological effects, the mean times to absorption (either extinction or fixation) (a) and to extinction (b) are exponentially longer for moderately beneficial mutants with 0 ≲ *s*_inv_ ≲ *η*, peaking at *s*_inv_ *≃ η/*2 (dashed lines) as predicted by our theory. The maximum mean times also grow exponentially with the effective population size *N*_eff_ . (c-h) Representative trajectories and phase portraits (in terms of the ratio between selection and drift) show distinct behaviors across different *s*_inv_. For low *s*_inv_ (f) or high *s*_inv_ (h), there is a single crossover (dashed lines) from drift to selection dominated regimes. Selection drives (green arrows) the mutant to extinction (c) or fixation (e). However, there are additional crossovers (g) when *s*_inv_ is moderate with *s*_inv_ ≲ *η*. Selection instead drives the mutant into a coexistence region centered at frequency *f*^*\**^ (dark red), resulting in exponentially long times for parent-mutant coexistence (d).

Such coexistence is a direct consequence of frequency-dependent selection (Eq. (3)) [66]. To see this, it is helpful to think about the infinite effective population size limit of Eq. (3). In this limit, we can ignore stochasticity and focus on the fixed points of the deterministic dynamics which are given by the solutions to the equation 0 = (*s*_inv_ − *ηf*) *f*(1 − *f*). When the fitness satis-fies *s*_inv_ *> η* or *s*_inv_ *<* 0, this equation has two fixed points at *f* ^*\**^ = 0 and *f* ^*\**^ = 1. On the other hand, when 0 ≤*s*_inv_ ≤ *η*, this equation has an additional fixed point at 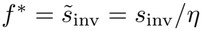, implying that the parent and mutant can potentially coexist in the absence of stochastic drift.

In the presence of drift, for small and large 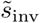 i.e., for deleterious and strongly beneficial mutants — selection almost always drives mutants to extinction (Fig. 3c) and fixation (Fig. 3e) respectively. On the other hand, for moderately beneficial mutants the presence of the additional fixed point in the deterministic limit gives rise to additional crossovers (Fig. 3g) which create a “coexistence region” between extinction and fixation, centered at frequency 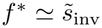. Selection drives the mutant into this region instead of extinction or fixation, hence the parent and mutant coexist for exponentially long times (Fig. 3d) before escaping the region (SI Sec. H). Such a region is absent in classical population genetics with constant selection.

Parent-mutant coexistence also qualitatively explains the suppression of fixation probabilities for moderately beneficial mutants with 0 *< s*_inv_ *< s*_*c*_ (Fig. 2b). Recall that in classical population genetics, beneficial mutants reach fixation through establishment, i.e., crossing the drift threshold from drift to selection dominated regimes (Fig. 4a, see also Fig. 3f and h). The process of establishment is central to Kimura’s theory, since after this point, mutants grow almost deterministically towards fixation. However, this qualitative picture is modified by the coexistence region under strong ecological effects, as selection now drives mutants into this region after establishment (Fig. 3g). When the coexistence region is closer to ex-tinction (Fig. 4b), i.e., *f* ^*\**^ *<* 1*/*2 or *s*_inv_ *< s*_*c*_, mutants that escape this region due to stochastic drift are more likely to go extinct than fix. Since such extinction occurs even after crossing the drift threshold, it suppresses fixation probabilities relative to Kimura’s prediction. On the other hand, when the coexistence region is closer to fixation (Fig. 4c), i.e., *f* ^*\**^ *>* 1*/*2 or *s*_inv_ *> s*_*c*_, it is more likely for mutants inside the region to drift to fixation instead, hence in this case coexistence does not suppress fixation.

**FIG. 4:**
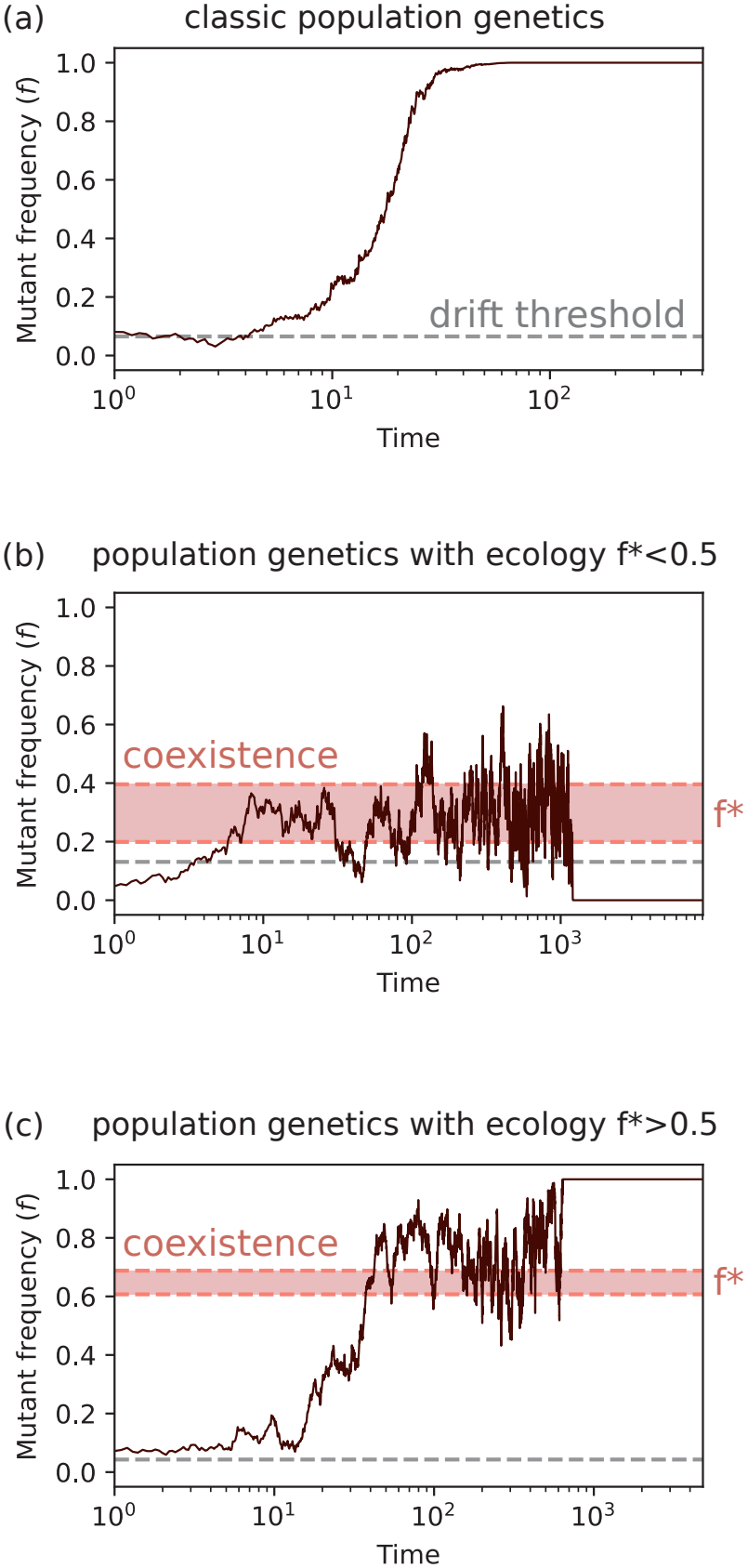
Suppression of fixation probabilities explained by parentmutant coexistence. (a) In classic population genetics, mutants are very likely to reach fixation after crossing the drift threshold (dashed gray line) at low frequencies, a process known as establishment. (b, c) With ecological effects, the coexistence region (dark red, Fig. 3(g)) modifies the fate of mutants. When the coexistence region is closer to extinction (b), i.e., *f*^*\**^ *<* 1*/*2, the region acts as an additional barrier to fixation, causing extinction even after crossing the drift thresh-old. This mechanism accounts for the suppression of fixation probabilities (Fig. 3b). In contrast, the coexistence region no longer prevents mutants from fixation when it becomes closer to fixation (c), i.e., *f*^*\**^ *>* 1*/*2.

### Mutant dynamics reveal ecological feedbacks in the human gut microbiome

A key prediction of our theory is that the strength of frequency-dependent selection *η* encodes the ecological feedbacks acting on a mutant. In practice, we can directly infer *η* from data by fitting Eq. (3) to a set of mutant frequency trajectories from the same community, similar to previous approaches [76].

To explore this application, we analyzed longitudinal metagenomic sequencing data from the gut microbiome of a single patient, sampled across 19 fecal samples over 5 months [75]. Deep sequencing allows us to infer frequency trajectories of single nucleotide variants (SNVs), which serve as proxies for mutant dynamics (SI Sec. K). These SNVs span 17 species from the same community (Fig. 5b), allowing us to estimate *η* by fitting Eq. (3) (Methods). During the 5-month period, the patient was diagnosed with disease at the end of the first month and received a two-week course of antibiotics at the end of the second month (Fig. 5). These perturbations have the potential to change community composition, resulting in an ecological feedback strength that shifts over time. To account for this possibility, we split the data into baseline, disease, and recovery periods (Fig. 5a). Further, we excluded the period shortly after the antibiotic treatment, during which the community underwent rapid compositional shifts, violating our assumption of ecological steady state. Note that our model does not require mutants to differ from parents by a single point mutation; they may differ by several SNVs, as is the case here. For simplicity, in our analysis we assumed that all mutants in the same community experience a common average ecological feedback strength *η*. In reality, different mutants may experience slightly different ecological feedbacks due to, e.g., differences in their phenotypic correlation *ρ* with their parents Taken together, this dataset is well suited for applying our model: it tracks the frequency dynamics of several mutants in a diverse community that appear to coexist for a long period of time (in fact, some of these were previously interpreted as supported by spatial or metabolic niches [75]).

**FIG. 5:**
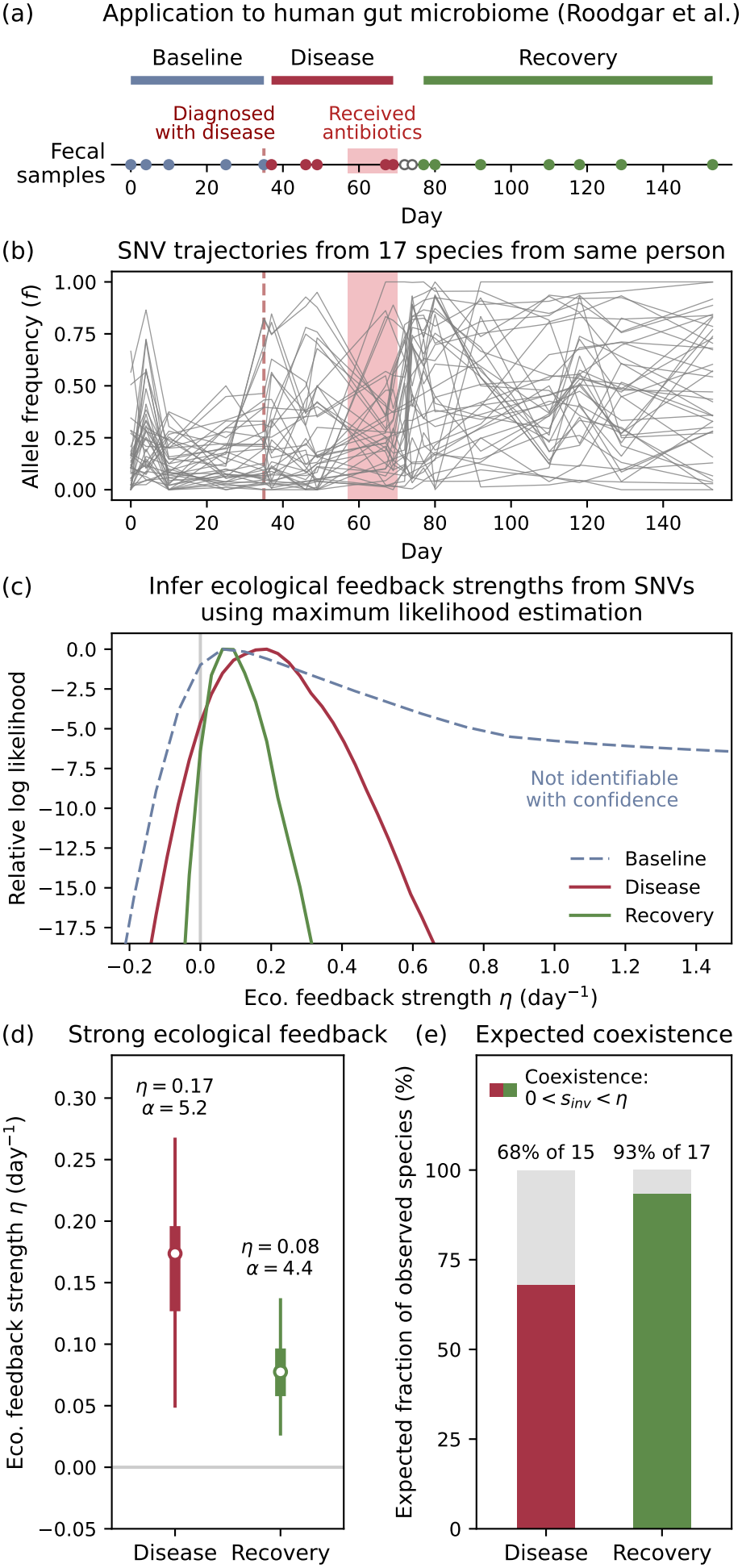
Mutant frequency dynamics recapitulate strong ecological feedbacks and parent-mutant coexistence in the human gut microbiome. (a) We analyze longitudinal sequencing data from the gut microbiome of a single patient across 19 fecal samples collected over 5 months [75]. The patient was diagnosed with disease at the end of the first month and received antibiotics for 2 weeks around day 60 (pink). We split the timeline into baseline, disease, and recovery periods, excluding the period shortly after antibiotic treatment when the community underwent substantial compositional shifts. (b) Frequency trajectories of sweeping single nucleotide variant (SNV) clusters [75] from 17 selected species within the same microbiome. (c) We infer the strength of ecological feedback *η* in different periods by fitting the trajectories to the model of frequency-dependent selection (Eq. (3), Methods). Maximum likelihood estimation yields confident estimates of *η* in the disease and recovery periods, but not in the baseline period. (d) The inferred *η* in disease and recovery phases are both positive. Thick and thin lines denote 50% and 95% confidence intervals obtained by bootstrapping across species. Combined with the inferred effective population sizes *N*_eff_ (SI Sec. K), these estimates indicate strong ecological effects in both periods, with 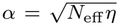 around 4 to 5 (Eq. (7), SI Sec. K). (e) We also infer the distribution of invasion fitness *s*_inv_ across species (SI Sec. K). In both periods, parent-mutant coexistence is expected for majority of the selected species, based on the fixed-point condition 0 *< s*_inv_ *< η* (Fig. 3).

We fit Eq. (3) to this dataset using a maximum likelihood estimation (MLE) method we developed (SI Sec. K). This method allowed us to infer the average strength of ecological feedbacks *η*, the average effective population size *N*_eff_, and the distribution of invasion fitness *s*_inv_ across different mutants (SI Sec. K). We then used the best-fit parameters to: (1) test whether the data are consistent with widespread negative frequency-dependent selection, i.e., *η >* 0 predicted by our theory; (2) estimate the strength of ecological effects 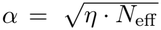 and (3) estimate the fraction of mutants that could coexist with their parents due to community ecological feedbacks. For simplicity, we also assumed there was only one mutant and one parent strain per species. This assumption is consistent with previous studies where two strains per species are commonly observed [7, 75, 77–79] Our MLE procedure led to confident estimates of *η* in the disease and recovery periods,but not in the baseline period. We confirmed that the best-fit models in these two periods were well-calibrated for one-step-ahead predictions (*p* = 0.60 and *p* = 0.15 respectively; Cramér-von Mises criterion, SI Sec. K).

Encouragingly, we found that the inferred *η* were positive during both the disease and recovery periods (Fig. 5d). This is consistent with our prediction that communities typically exert negative frequency-dependent selection on mutants (SI Sec. A–C). Combined with the estimates *N*_eff_ ≃160 and *N*_eff_ ≃250 (normalized by the number of generations within a day) for the two periods respectively (SI Sec. K), these results suggest that mutants in the gut microbiome are subject to strong ecological effects *α >* 1 in both periods, with *α* ≈ 4-5 (Eq. (7)). Although the inferred *η* is slightly lower in the recovery period, more data are required to statistically resolve the difference between the two periods.

Our theory also predicts that strong ecological effects promote parent-mutant coexistence when 0 *< s*_inv_ *< η* (Fig. 3). To examine this prediction, we used the inferred distribution of *s*_inv_ across species to estimate the expected prevalence of parent-mutant coexistence (SI Sec. K). We found that the inferred parameters are consistent with parent-mutant coexistence for most of the observed species in the community (Fig. 5e). Together, these results demonstrate how mutant frequency dynamics can be used to infer strong ecological effects and their implications across the community.

## DISCUSSION

In this study, we developed an analytic framework for integrating ecological interactions with a complex community into population genetics models. Our calculations show that interactions with a highly-diverse community can dramatically affect evolutionary outcomes. Despite their complexity, ecological feedbacks give rise to a remarkably simple form of frequency-dependent selection between ecologically similar parent and mutant strains when the community is near its ecological steady state. Surprisingly, the frequency dependence from the entire community can be captured by a single parameter *η* which can be easily calculated from the community’s statistics for a wide variety of ecological models. Thus our results reveal a principled method to apply existing population genetics models with frequency-dependent selection even in the presence of complex communities.

We find that when ecological feedbacks are strong, the fixation probabilities for moderately beneficial mutants are suppressed compared to Kimura’s predictions for constant selection (Fig. 2b) [24]. The suppression increases with effective population size (Fig. 2e) [24], weaker parent-mutant correlations (Fig. 2e), and decreases as ecosystems become more packed (Fig. 2f). We further show that the deviations from Kimura’s formula arise from prolonged parent-mutant coexistence arising from ecological interactions (Fig. 4b and c) [28].

This coexistence results in exponentially long absorption (Fig. 3a) and extinction times (Fig. 3b) for moderately beneficial mutants [24, 26, 28], as well as coexistence regions in the phase portraits of parent-mutant dynamics (Fig. 3g).

Our theory is robust to the details of the exact ecological model used to mathematically represent the community dynamics. We show that our framework applies to various ecological models including generalized Lotka-Volterra models and multiple variants of the consumer-resource model (SI Sec. A–C), which all support regimes with parent-mutant coexistence. Therefore, consistent with [15] for deterministic equilibria, our analysis suggests that the possibility of parent-mutant coexistence is a hallmark of population genetics in complex communities, and we expect the framework developed here to be widely applicable. As an exploration, we applied our framework to understand ecological feedbacks in data from the human gut microbiome [75] (Fig. 5).

Our theory makes testable predictions for high-resolution strain tracking in natural microbiomes [75, 79]. With high enough temporal resolution, strain dynamics should exhibit intermittent stochastic fluctuations in a range of frequencies *above* the drift barrier (Fig. 4b–c). These dynamics should be most pronounced for moderately beneficial mutants with *s*_inv_ ≲ *η/*2. Importantly, estimating both *f* ^*\**^ and *s*_inv_ from observed trajectories could enable direct inference of the ecological feedback strength *η* from strain dynamics alone. Together with the analysis in Fig. 5, our theory suggests a new empirical technique to estimate average ecological feedbacks on evolutionary dynamics [80].

Our findings also have broader implications for inference using population genetics models. Current methods largely neglect ecology and for this reason may yield biased estimates of effective population sizes, demographic histories, and selection coefficients. Designing predictors that incorporate *η* into coalescent models and demographic inference frameworks has the potential to improve the accuracy of statistical methods, especially in settings where ecology plays an important role in the evolutionary dynamics. Our results also have important implications for conservation genetics. They suggest that traditional metrics that ignore ecology may under- or over-estimate extinction risks.

In the future, it will be useful to extend our theory to a wider range of ecological and evolutionary settings. On the ecological side, while our theory has so far focused on communities near steady state, ecosystems can exhibit more complex dynamical behavior such as limit cycles, multistability [81, 82], and chaos [53, 56, 83–85]. Extending our theory to these settings will enable us to understand how the fate of a mutant depends on the underlying dynamical behavior of a community.

On the evolutionary side, for many species, especially microbes, mutation rates are sufficiently large so that multiple mutants of the same species can emerge before previous ones fix or go extinct [86]. This phenomenon challenges our assumption of the strong-selection-weak-mutation regime, since we have let a mutant to coexist transiently with its parent without new mutants. Mathematically, our assumption holds when the stochastic extinction time of a coexisting mutant is faster than the time for new mutations (see SI Sec. H for precise mathematical condition). This condition is typically violated in many microbial communities due to their large population sizes and mutation rates. In this case, multiple mutant lineages compete with each other simultaneously, a phenomenon known as clonal interference [7, 86, 87]. In the future, it would be interesting to extend DMFT approaches like the one we developed here to understand the effect of clonal interference on eco-evolutionary dynamics in complex communities.

On the modeling front, our model relies on the assumption that mutant strains have high phenotypic correlation *ρ* ≈1 with their parents. While this assumption might be reasonable for mutations with small effect sizes, the lack of genotype-phenotype maps in ecological contexts makes it challenging to estimate *ρ* from community data. Future work on better understanding how genotype-phenotype maps [88] affect ecological dynamics could help test the applicability of our approximation [89].

Finally, our theory assumes random and dense ecological interactions, and mutations that weakly alter many interactions, allowing feedbacks to be summarized using community-level statistics. However, ecological networks may become more nonrandom through ecological assembly [90, 91] or longer-term eco-evolution [14, 15, 50, 59]. Mutational effects may also be structured and context-dependent [89, 92, 93], affecting only a few partners or changing multiple interactions coherently. In the future, it will be useful to extend our framework to these scenarios.

## METHODS

The details of all the analytical and numerical methods used can be found in the Supplementary Information. Briefly, we started with defining the equations for the generalized Lotka-Volterra model and various consumer-resource models, designating two of the species to be a parent and mutant. We analyzed these equations using dynamical mean-field theory and other approximations to derive Eq. (3) and the value of *η*. We then solved the resulting stochastic differential equation to obtain fixation probabilities (Eq. (5)), selection-drift ratio, and mean extinction time (Fig. 3). We performed numerical simulations by sampling random ecosystems at steady states, then explicitly solved the equations in the ecological models using the Euler method with demographic noise [58, 94, 95]. We repeated the same simulation with multiple instances of demographic noise to obtain the statistical quantities such as fixation probabilities and mean absorption/extinction times. Finally, we fitted the SNV trajectories (Fig. 5) to Eq. (3), accompanied with a model of measurement noise, using maximum likelihood estimation. We provide details on the data analysis including the fitted model, the algorithms for maximum likelihood estimation, the full set of inferred values with bootstrapping, and the procedures for testing the goodness of fit on one-step-ahead predictions. This analysis was assisted by Claude Opus 5.5 for consultation of statistical approaches and production of the analysis code.

## Acknowledgements

We would like to thank Michael M. Desai and Benjamin H. Good for helpful discussions. This work was funded by NIH NIGMS R35GM119461 to PM and Chan-Zuckerberg Initiative Investigator grant to PM. SYL acknowledges support from the National Science Foundation through the Physics Frontier Center for Living Systems (PHY-2317138). ZF acknowledges support by the NSF-Simons National Institute for Theory and Mathematics in Biology (NITMB) Fellowship supported via grants from the NSF (DMS-2235451) and Simons Foundation (MPS-NITMB-00005320). AG acknowledges support from the Ashok and Gita Vaish Junior Researcher Award, the DST-SERB Ramanujan Fellowship, as well the DAE, Govt. of India, under project no. RTI4001. This work was completed in part with resources provided by the University of Chicago’s Research Computing Center. Claude Opus 5.5 and ChatGPT-6 Astra were used to assist with minor editing of the manuscript, writing the simulation code, and checking the theoretical calculations.

## Appendix A: Lotka-Volterra model

We begin with the details of our eco-evolution models. Consider an ecological community of *S* competing species, with the abundance of each species *N*_*i*_ (where 1 ≤ *i* ≤ *S*) described by the Lotka-Volterra dynamics:

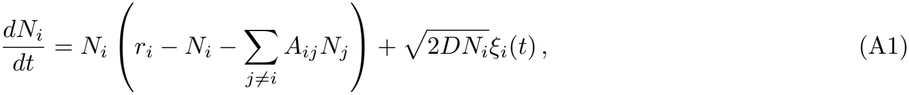

where *r*_*i*_ is the intrinsic growth rate of species *i*, and *A*_*ij*_ is the competition coefficient between species *i* and *j*. We have also introduced a white demographic noise *ξ*_*i*_(*t*) for each species with diffusion coefficient *D*, satisfying

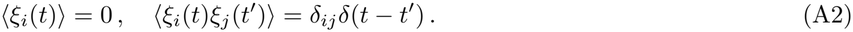

To model the diversity of this community, we work in the limit *S*≫ 1 and assume that *r*_*i*_ and *A*_*ij*_ are randomly drawn from Gaussian distributions, such that

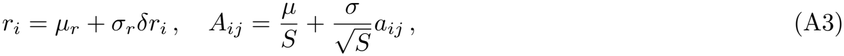

where *µ*_*r*_, *σ*_*r*_, *µ, σ* are the distribution parameters, and *δr*_*i*_, *a*_*ij*_ are zero-mean Gaussian random variables with

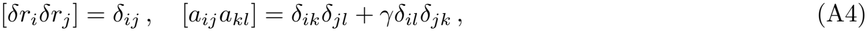

where the square brackets denote averages over random parameters instead of the demographic noise. The reciprocity − 1 ≤*γ* ≤1 is the correlation between the off-diagonal entries *A*_*ij*_ and *A*_*ji*_. The results below also apply to other distributions with mean and variance of order 1*/S*, and higher order in 1*/S* for higher cumulants.

Suppose at time *t* = 0, a mutation of a parent strain *p* occurs and a new mutant strain *m* with small abundance *N*_*m*_(0) ≪*N*_*p*_(0) invades the community. Biologically, the mutant should be highly similar to its parent; we assume the parent-mutant correlation of ecological interactions *ρ* to be close to 1. We sample the competition coefficients for the mutant as

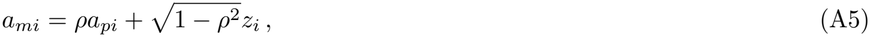

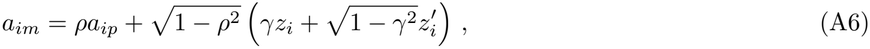

where 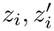 are independent zero mean, unit variance Gaussian random variables, and indices *i, j* now denotes the rest of the community without the parent or the mutant. These choices of coefficients ensure the right correlations Corr (*A*_*pi*_, *A*_*mi*_) = Corr (*A*_*ip*_, *A*_*im*_) = *ρ* and Corr (*A*_*mi*_, *A*_*im*_) = *γ*. We also have Corr (*A*_*pi*_, *A*_*im*_) = Corr (*A*_*mi*_, *A*_*ip*_) = *γρ*. On the other hand, the competition between the parent and the mutant should be much stronger than that with other species, controlled by their niche overlap. In the main text, we assume

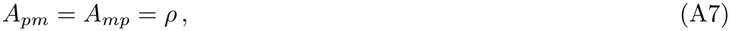

but here we leave them to be arbitrary for generality as long as 1 −*A*_*pm*_ and 1 −*A*_*mp*_ are at order 1− *ρ*. As defined in Eq. (A3), the other interaction coefficients *A*_*pi*_, *A*_*mi*_, *A*_*ip*_, *A*_*im*_ (where *i* ≠ *p, m*) are of order 1*/S* and much smaller than *A*_*pm*_, *A*_*mp*_. Finally, we do not specify any certain values for the intrinsic growth rate *r*_*m*_, in order to obtain the functional dependence of the fixation probability and to cont rol the order of 1 − *ρ* in selection. If we sample *r*_*m*_ as how we sample *A*_*mi*_, the difference *r*_*m*_ − *r*_*p*_ would be at order 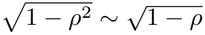. Throughout the analysis below, we make use of the small 1 −*ρ* expansion to simplify our results; see also Appendix F for more validation on results. Including the parent-mutant dynamics, the full set of Lotka-Volterra equations are

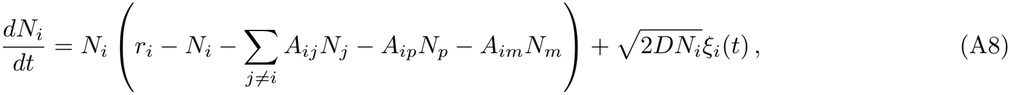

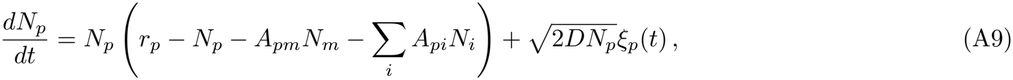

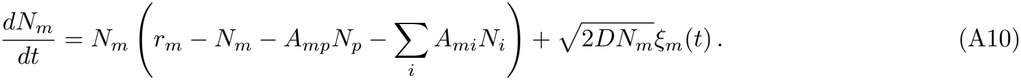

We focus on the “strong-selection-weak-mutation” limit: we assume an ecological steady state and let the mutant invade when the community is close to the steady state, then we let the community dynamics run till the parent or the mutant becomes extinct, even though the parent and mutant may transiently coexist for a long time. This approximation is one of the main assumptions of our analysis, which ignores the possibility of having multiple mutations of the same species in the community simultaneously due to nonzero mutation rates. By assuming an ecological steady state, we also require sufficiently small *D* such that demographic fluctuations around the deterministic steady state are small over the timescale we study, and that other species in the community do not go extinct during mutant dynamics. This is a self-consistent requirement: we show in Appendix H that under small 1 −*ρ*, the stochastic extinction time of a species is indeed much longer than that of the mutant. Furthermore, since selection (terms such as *r*_*m*_ − *r*_*p*_) is suppressed by 1 − *ρ, D* should also be suppressed by 1 − *ρ* accordingly in the strong-selection regime.

### 1. Dynamical mean-field theory (DMFT)

We now simplify the above set of Lotka-Volterra equations using DMFT. Instead of the abundances themselves, we will focus on the dynamics of the mutant frequency, as is typical in this literature:

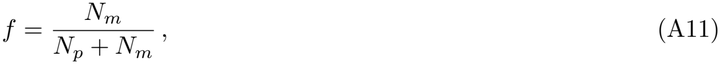

which changes dynamically and stochastically till reaching the point *f* = 0 or *f* = 1. We denote the initial frequency as *f*_0_ ≪ 1. Using the Lotka-Volterra equation, we have

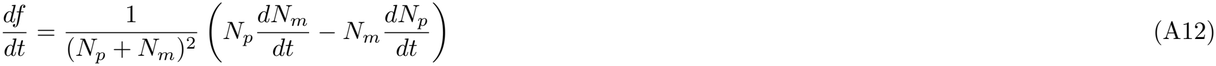

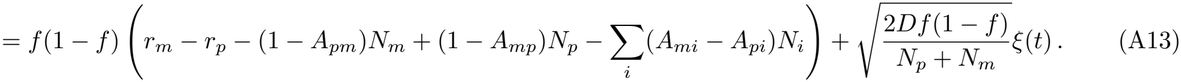

Here we have combined the noise terms of the parent and the mutant into a single white noise by adding their variances. The ecological feedback from the community on parent-mutant dynamics is determined by how the parent and the mutant impact other species in the community. We see that in Eq. (A8), the abundances *N*_*p*_, *N*_*m*_ serve only as a perturbation of order 1*/S*. We can then model their influence to *N*_*i*_ as a linear response. We write

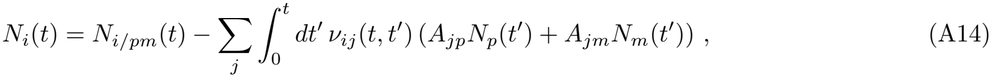

where *N*_*i/pm*_ is the abundance of species *i* if *both* the parent and the mutant are excluded from the community, and

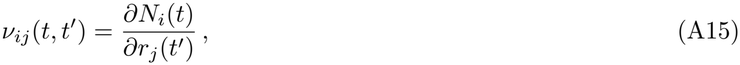

is the susceptibility kernel. Using the linear response, we can now write the community contribution in Eq. (A12) as

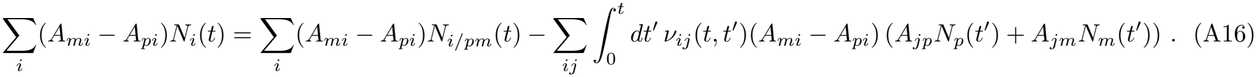

For the first term in Eq. (A16), we use the assumption that the community without the parent or the mutant fluctuates around its ecological steady state, hence we decompose

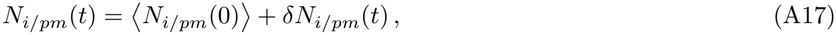

where we recall that the angle bracket denotes the average over demographic fluctuations, and *δN*_*i/pm*_(*t*) is an effective noise term. For the last term in Eq. (A16), it can be simplified at leading order with the self-averaging property of the community:

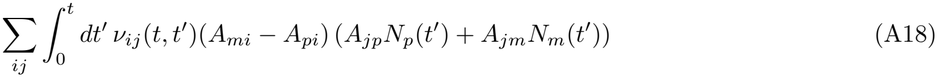

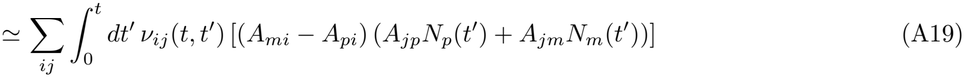

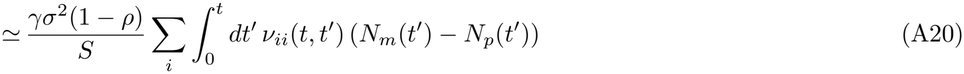

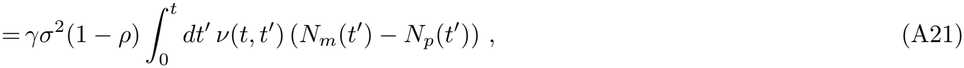

where we have introduced the susceptibility

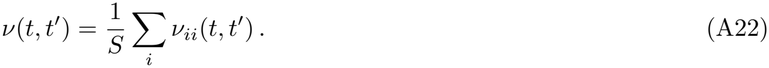

**FIG. S1:**
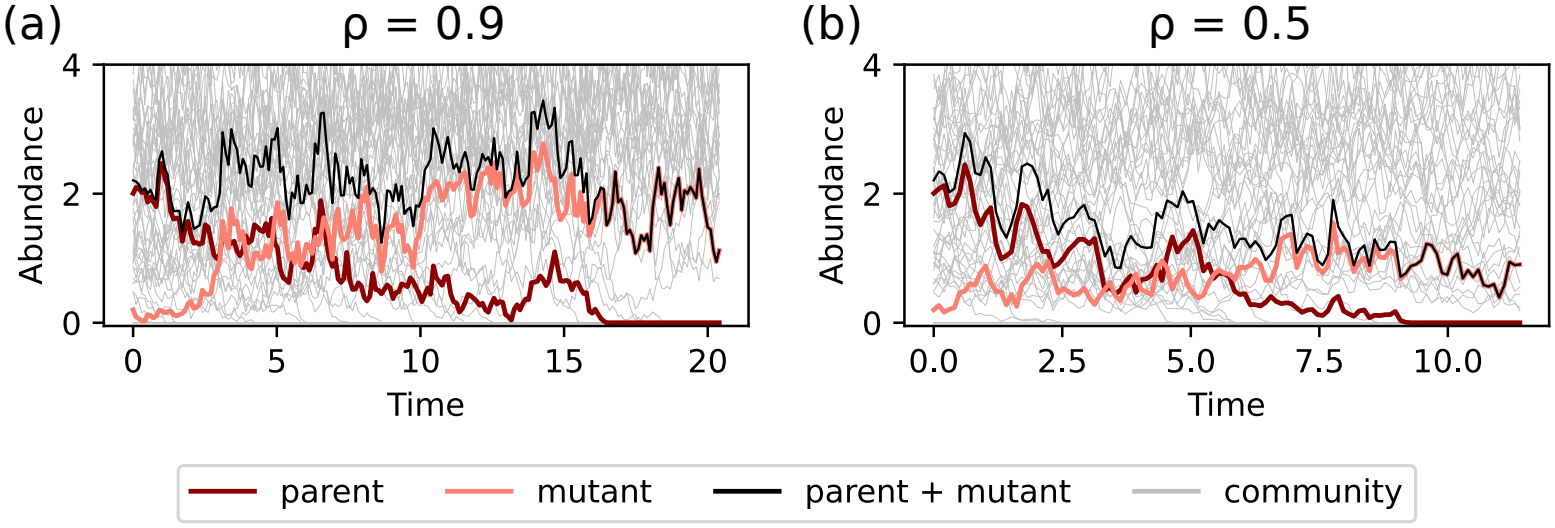
Dynamics of parent and mutant abundances during mutant fixation at various parent-mutant correlation. Fixation of a mutation generally requires deviation from the deterministic ecological steady state. At high *ρ* = 0.9 (a), the total parentmutant abundance remains constant on average throughout the full dynamics. At low *ρ* = 0.5 (b), the total parent-mutant abundance drops significantly towards the final mutant abundance.

Therefore, the mutant frequency satisfies

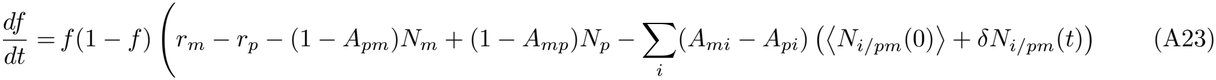

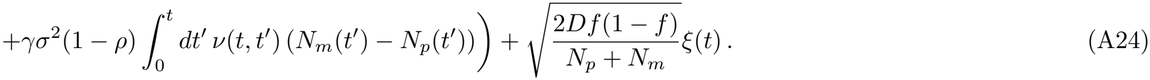

This is the DMFT equation for the mutant frequency with effective colored community noise and response, which in principle should be determined self-consistently together with the community dynamics. Note that in the conventional DMFT of the GLV, the term similar to the ⟨ *N*_*i/pm*_(0) ⟩ term in Eq. (A23) also contributes to the community noise, even when the full dynamics is deterministic. This is because the cavity species can be chosen to be any species in the community, hence there is noise from heterogeneity in interactions across different species. Here, the cavity species is instead a specific pair of parent and mutant with quenched interactions, hence the ⟨ *N*_*i/pm*_(0) ⟩ term is deterministic. Nevertheless, there is still community noise from demographic fluctuations in *δN*_*i/pm*_(*t*).

### 2. Small 1 *− ρ* expansion

We now show that the DMFT equation implies that the dynamics of *f* are slower than those of the community. This is because all the terms in Eq. (A23) are suppressed by small 1 − *ρ* to various extents. To obtain an equation involving *f* only, we then make further approximations using the small 1 − *ρ* expansion.

Qualitatively, when the parent and the mutant are highly similar in their ecological interactions with the community, the total parent-mutant population behaves effectively as a single species from the community perspective. Therefore, the community as a whole stays near the ecological steady state. This is true even when the parent and the mutant frequencies are rather different from their steady state frequencies during the dynamics. As a result, the total parentmutant abundance *N*_*p*_ +*N*_*m*_ always stays constant on average, with fluctuations coming from small demographic noise only. This is also consistent with Kimura’s theory, which assumes a constant effective population size. In contrast, when *ρ* is sufficiently less than 1, *N*_*p*_ + *N*_*m*_ may not be stationary on average when a mutant fixes, as shown in Fig. S1. In conclusion, under the small 1 − *ρ* expansion, we can replace

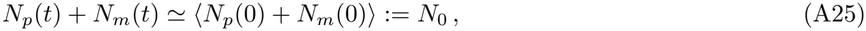

and the effective population size is *N*_eff_ = *N*_0_*/*2*D*. Note that we have not specified the physical unit of time for the dynamics; to match with real experiments, *N*_eff_ should be further normalized by the actual generation timescale.

Next, based on the ecological steady state, we calculate the community noise and the response in the DMFT equation and show that they can be greatly simplified under small 1 −*ρ*. Namely, the community noise is subdominant compared to the demographic noise, and the dynamic susceptibility can be replaced by a static one.

First, consider the deterministic version of the community without the parent and the mutant. The abundances of surviving species 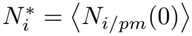 satisfies the fixed point condition

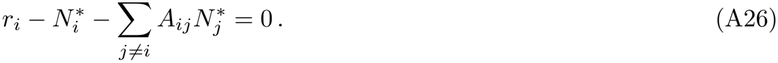

Next, we add two sources of perturbations: the demographic noise, and the parent-mutant dynamics as an effective *δr*_*i*_(*t*). Accordingly, we decompose 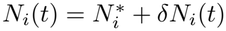 for the linear response. The GLV dynamics becomes

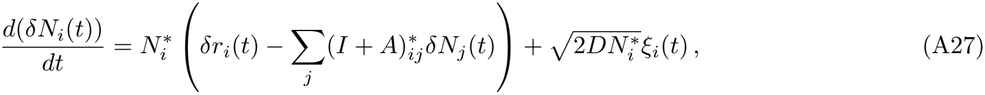

where the “star” on the matrices means taking the components of surviving species only. We can solve this ordinary differential equation using standard methods to obtain

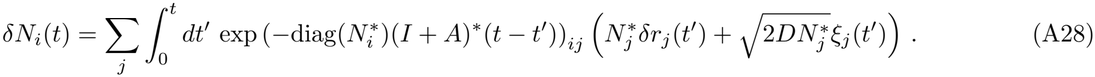

Using the definitions in Eqs. (A15) and (A17), we can then read off the susceptibility from the first term

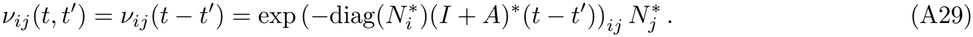

and the community noise from the second term

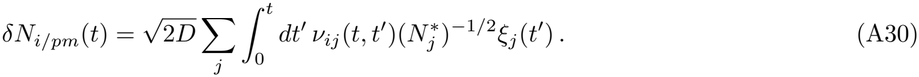

We now tackle each of the quantities separately. First, the susceptibility is time-translational invariant can be decomposed into a sum of exponential decays in time difference *τ* = *t* − *t*^*′*^. Notice that all the decay rates, given by the eigenvalues of diag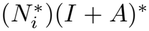, are not suppressed by 1 − *ρ*. Therefore, since the derivatives of *δr*_*i*_(*t*) are suppressed by 1 − *ρ* as in the frequency dynamics, we can approximate the response as

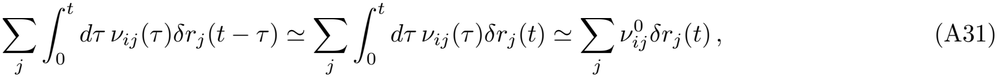

where

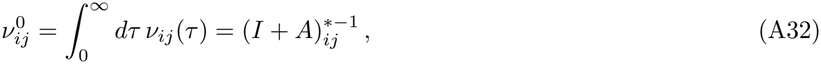

is the static susceptibility. Equivalently, we have approximated the dynamical susceptibility with a delta function 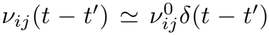 To numerically verify these approximations, we also compute the susceptibility function directly from the GLV simulation; see Appendix I for methods. The results in Fig. S2 indeed match with our theory.

**FIG. S2:**
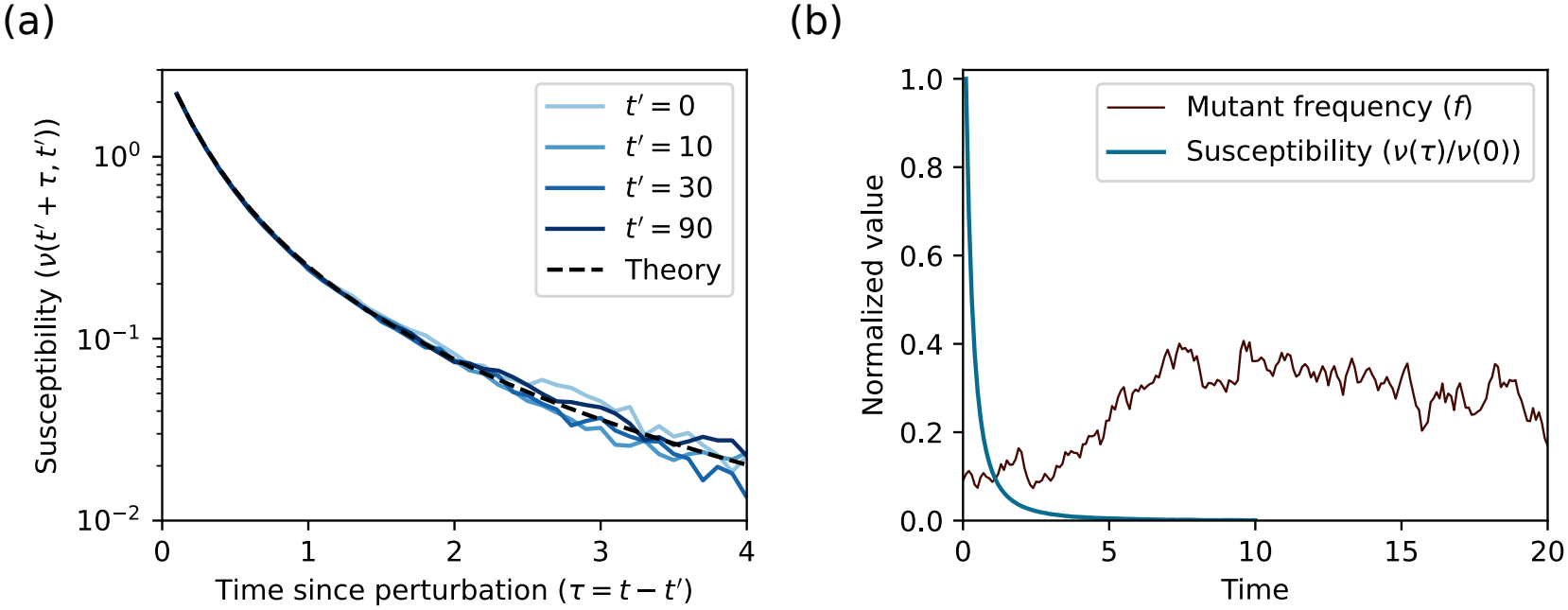
Dynamical susceptibility and comparison with the dynamics of mutant frequencies. (a) The susceptibility function is time-translational invariant and exponentially decaying, matching with the prediction of our theory (Eq. (A29)). (b) The susceptibility function decays much faster compared to the dynamics of mutant frequencies.

**FIG. S3:**
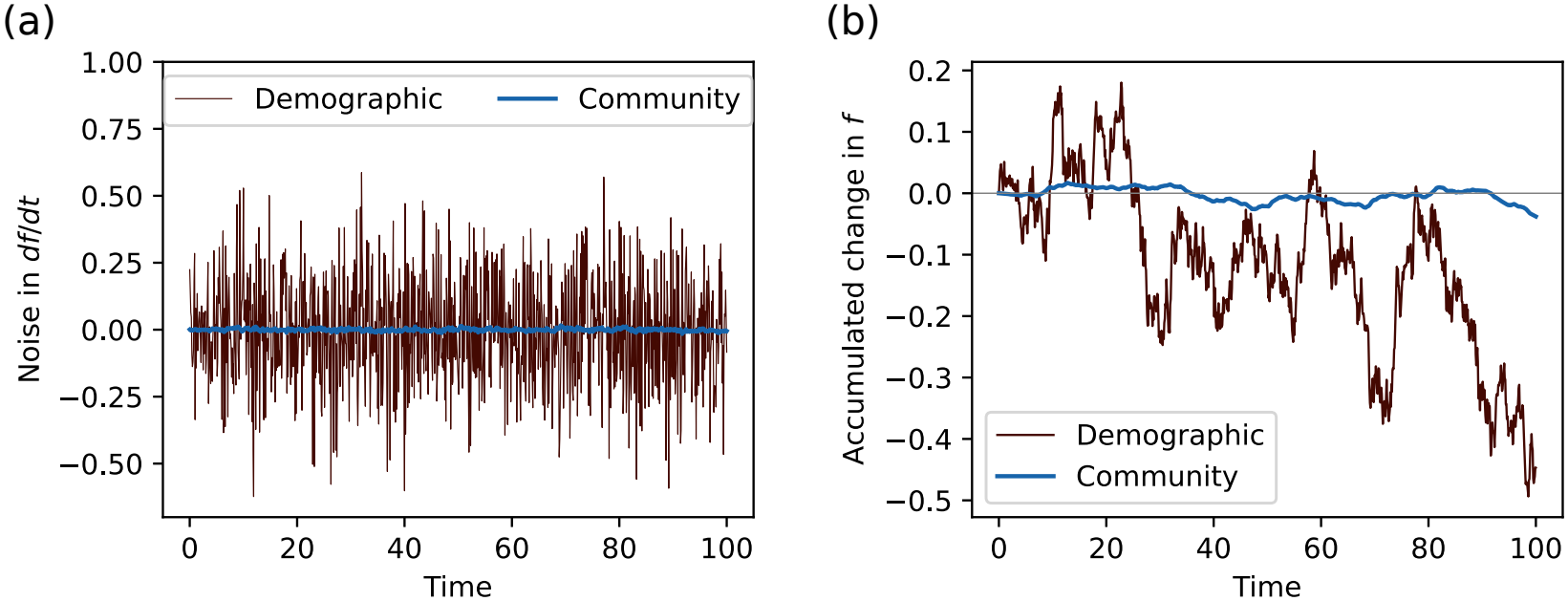
Community noise and comparison with demographic noise of the parent and the mutant. Given a mutation and its frequency trajectory, we re-simulate the community dynamics without the parent and the mutant. The resulting community noise is negligible compared to the demographic noise (or genetic drift) of the parent and the mutant, in terms of (a) the rates of change in *df /dt* and (b) the accumulated changes in *f* .

Next, for the community noise term in Eq. (A23), we can now calculate its covariance using the self-averaging property.

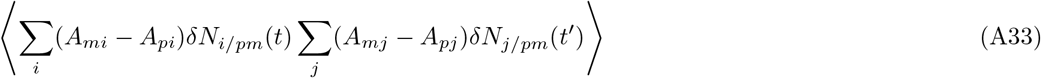

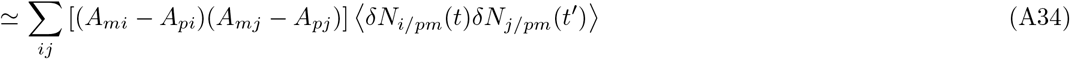

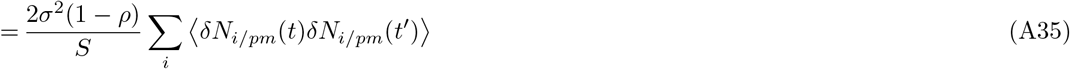

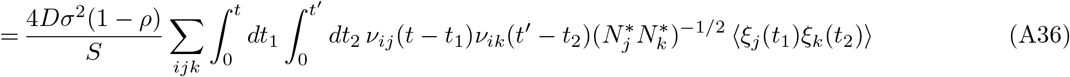

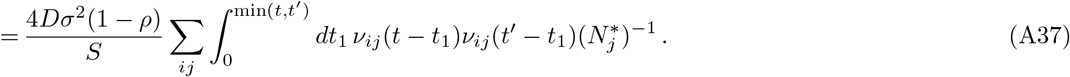

Note that the covariance decays in time in the same way as the susceptibility, hence the decay is faster than the frequency dynamics. We can then compare the community noise with the demographic noise in the following way. Suppose the frequency *f* evolves for a time Δ*t* that is longer than the timescale of covariance decay, but shorter than the timescale of frequency dynamics. In this regime, *f* is unchanged on average and we can focus on its variance. Since the demographic noise is given by a white noise *ξ*(*t*), its contribution to the variance is simply

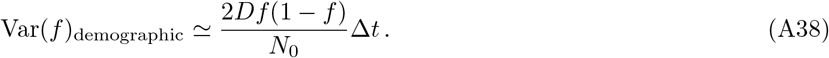

On the other hand, the variance from the community noise is

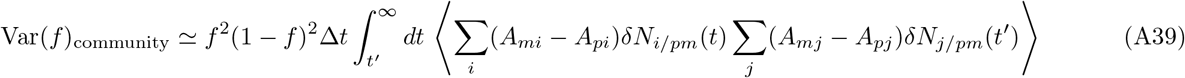

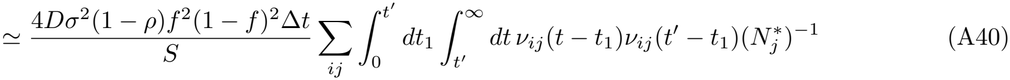

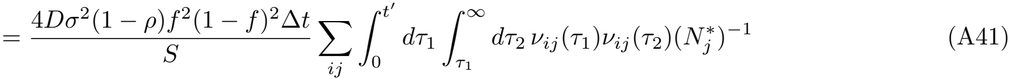

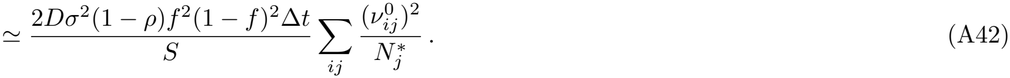

In other words,

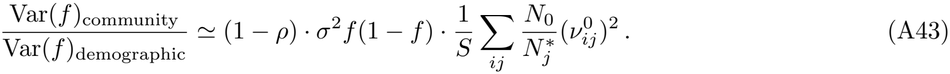

Since all the factors except 1 − *ρ* are *O*(1), we conclude that the demographic noise is always dominant over the community noise, hence we omit the community noise from now on. We also verify this approximation in the GLV simulation in Fig. S3.

### 3. Frequency-dependent selection

With all these approximations, we arrive at

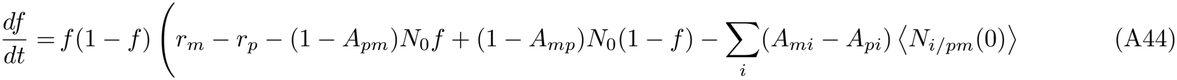

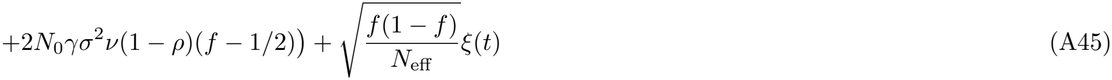

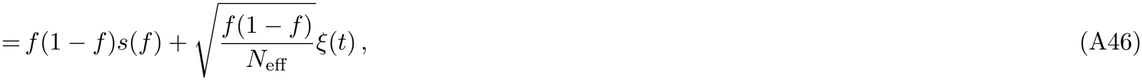

where *s*(*f*) represents frequency-dependent selection and the static susceptibility is

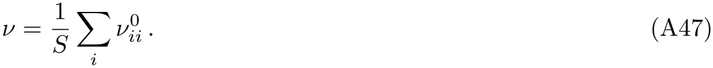

We see that *s*(*f*) is linear in *f*. To better interpret the above result, we note that *s*(*f*) is in fact the difference between the growth rates between the mutant and the parent at frequency *f*. The difference *s*(*f* = 0) is also the invasion fitness of the mutant *s*_inv_ and plays the same role as the selection coefficient *s* in classic population genetics. Therefore, it is useful to rewrite *s*(*f*) as a linear function in *f* involving *s*_inv_:

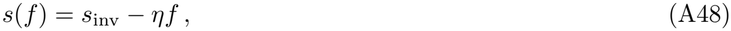

where

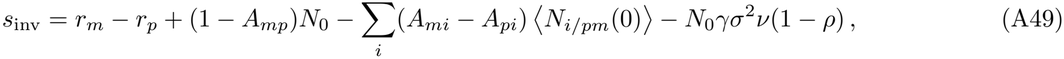

and

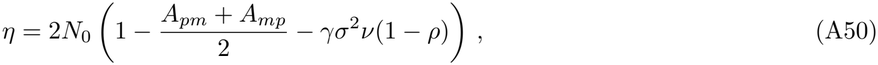

parametrizes the strength of the overall ecological effects. Note that the ecological feedback proportional to *v* also contributes to the invasion fitness since the parent was already in the community before *t* = 0. In conclusion, we get

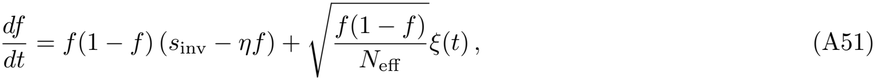

which is the same dynamics as in the main text. Note that all the three terms in the dynamics are suppressed by 1− *ρ* to different but independent extents. Therefore, one term may be dominant over the others depending on the parameter regime. Nevertheless, it is mathematically consistent in the sense that the effects we ignore in the previous section are all subleading compared to the dominant terms here.

To better understand the impact of the community, we can estimate the susceptibility using Eq. (A32), implying

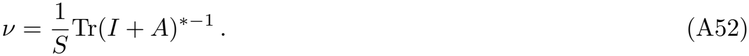

The trace is simply the resolvent of *A*^*\**^, which can be calculated using random matrix theory. The result is [96]

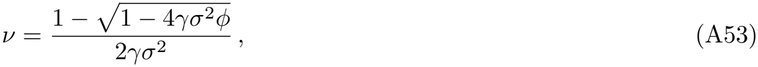

where *ϕ* is the fraction of surviving species. Since we have started with a near steady state of the community for the mutation, the above expression for *v* should also hold approximately after the mutation. Therefore, we also have

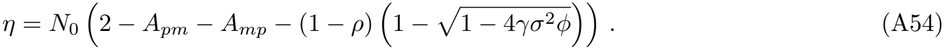

For the main text, we set *A*_*pm*_ = *A*_*mp*_ = *ρ* and the above nicely factorizes into

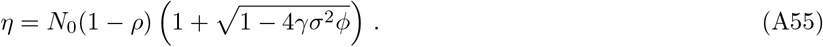

Note that this expression is valid only when *ϕ <* 1*/*4*γσ*^2^, which is an upper bound on the number of surviving species similar to May’s stability bound for ecological communities [55], since we have assumed that the community is near a stable state. This is the species packing bound referred in the main text.

## Appendix B: MacArthur consumer-resource model

Following the same procedure as in Appendix A, it is straightforward to extend the results to consumer-resource models. We will see that the mutant frequency follows the same dynamics as before, only with a different *η*.

We first study the MacArthur consumer-resource model with self-renewing resources. We consider an ecological community of *S* species and *M* resources. The species abundances *N*_*i*_ (where 1 ≤ *i* ≤ *S*) and resource abundances *R*_*α*_ (where 1 ≤ *α* ≤ *M*) follow

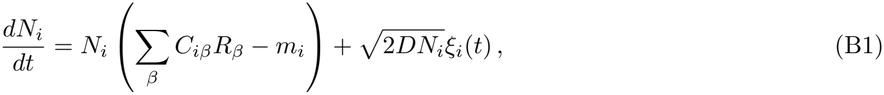

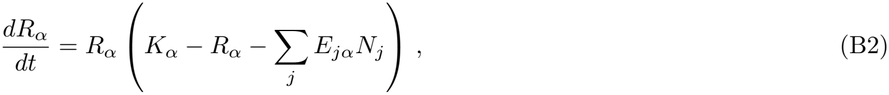

where *K*_*α*_ is the carrying capacity of resource *α, m*_*i*_ is the intrinsic mortality rate for species *i, C*_*iα*_ represents the consumption preferences of species *i* for resource *α*, and *E*_*iα*_ is the corresponding impact of species *i* on resource *α*. The demographic noise *ξ*_*i*_(*t*) is the same as in the previous section. For a diverse community, we work in the limit *S, M* ≫ 1 but finite *S/M*. We further assume that the parameters are randomly drawn from Gaussian distributions, such that

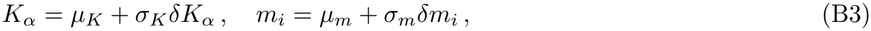

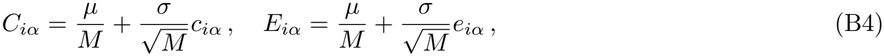

where *µ*_*K*_, *σ*_*K*_, *µ*_*m*_, *σ*_*m*_, *µ, σ* are the distribution parameters, and *δK*_*α*_, *δm*_*i*_, *c*_*iα*_, *e*_*iα*_ are zero-mean Gaussian random variables with

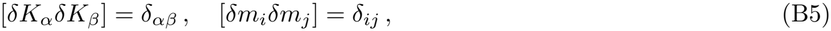

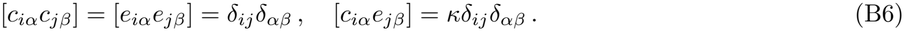

The reciprocity 0≤ *κ* ≤ 1 of species-resource interaction is the correlation between the matrices *C*_*iα*_ and *E*_*iα*_. Suppose at time *t* = 0, a mutation of a parent strain *p* occurs and a new mutant strain *m* with small abundance *N*_*m*_(0) ≪ *N*_*p*_(0) invades the community. Let the parent-mutant correlation be *ρ*. We sample the new interaction coefficients as

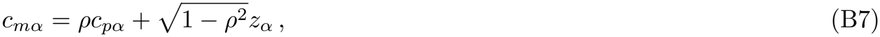

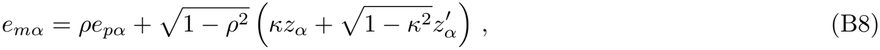

where 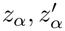 are independent zero mean, unit variance Gaussian random variables. These choices of coefficients ensure the right correlation Corr(*C*_*pα*_, *C*_*mα*_) = Corr(*E*_*pα*_, *E*_*mα*_) = *ρ* and Corr(*C*_*mα*_, *E*_*mα*_) = *κ*. We leave the mortality rate *m*_*m*_ to be arbitrary. Note that we do not introduce any new resources. After introducing the mutant, the consumer-resource dynamics becomes

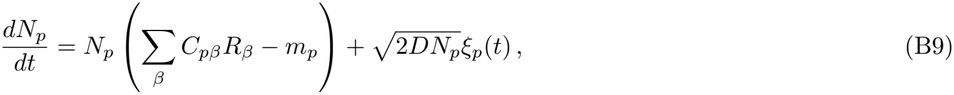

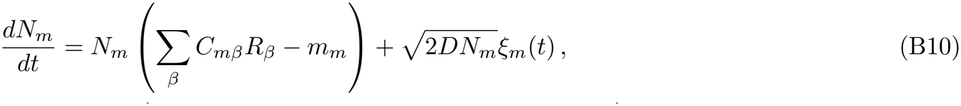

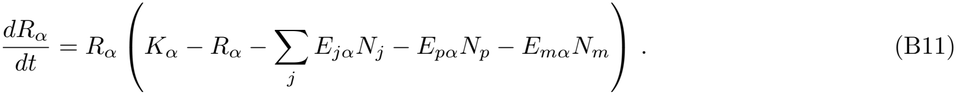

In terms of the mutant frequency *f* = *N*_*m*_*/*(*N*_*p*_ + *N*_*m*_), the above becomes

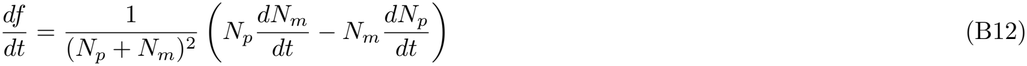

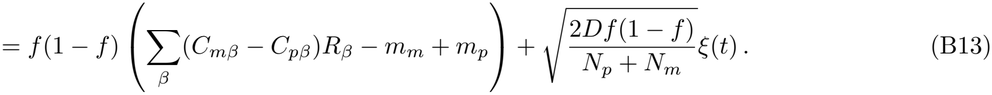

The dynamics of the other species *N*_*i*_ will not be relevant in the following calculation. Now we adopt the same set of approximations as before. We can model the changes of *R*_*α*_ as linear responses:

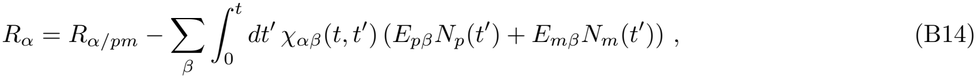

where

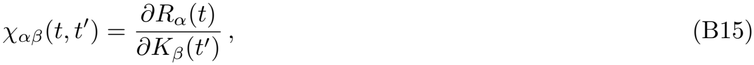

is the susceptibility kernel. Now we can write the resource contribution in Eq. (B12) as

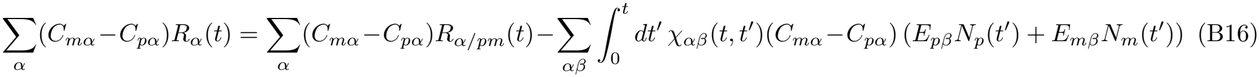

We can then apply self-averaging to the products between *C* and *E* and approximate the last term as

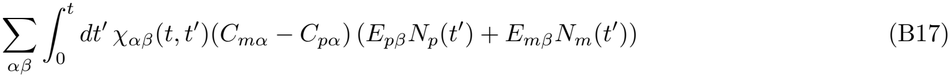

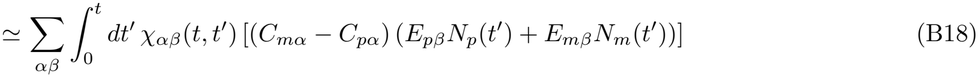

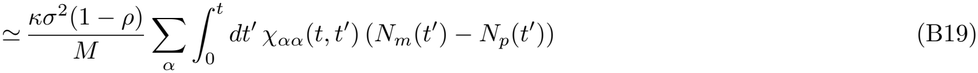

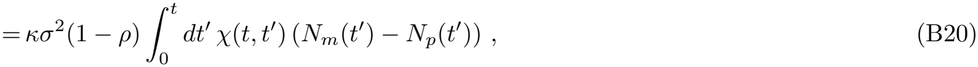

where we have introduced the susceptibility

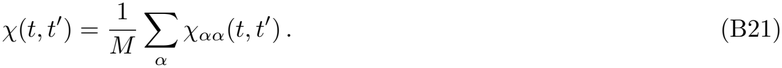

Therefore, the mutant frequency satisfies

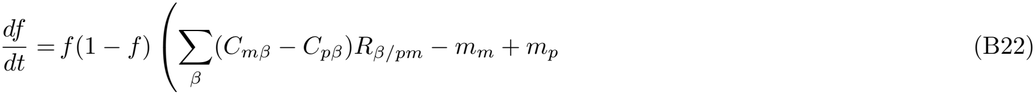

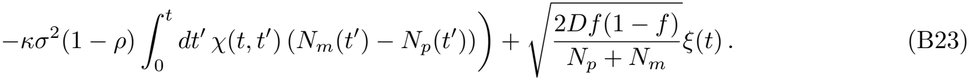

As before, we then further approximate

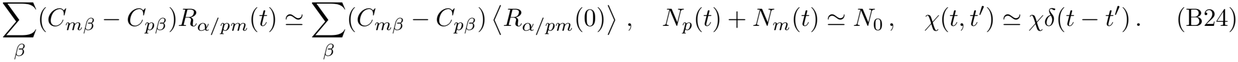

We then arrive at:

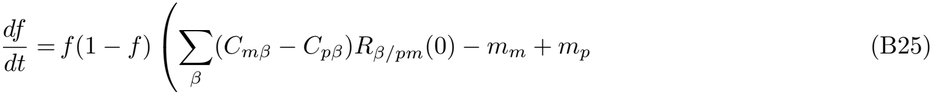

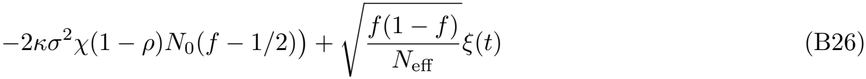

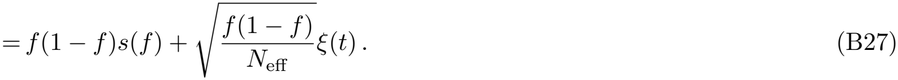

As before, we can separate the invasion fitness *s*_inv_ = *s*(*f* = 0) from *s*(*f*) and rewrite the dynamics in the form of Eq. (A51) as

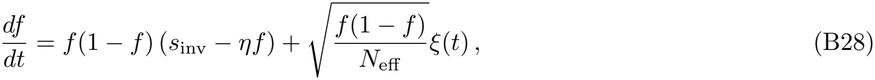

where the invasion fitness is

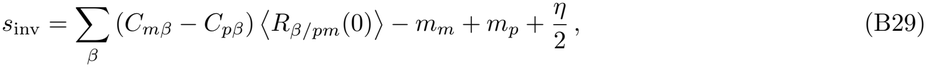

and the strength of ecological effect is

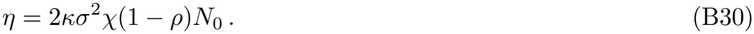

Now, we can estimate the susceptibility by the steady state conditions for surviving species and non-depleted resources (denoted by asterisks):

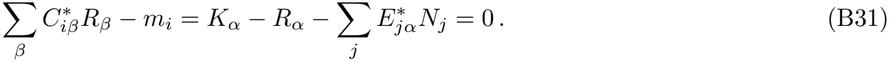

The variations satisfy

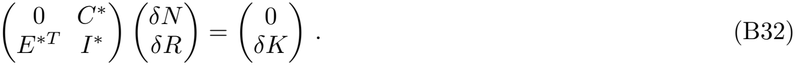

Using block matrix inversion, we have

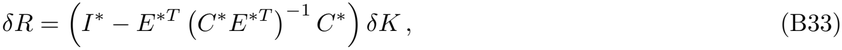

hence,

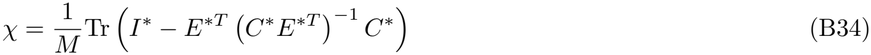

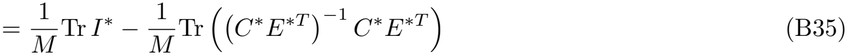

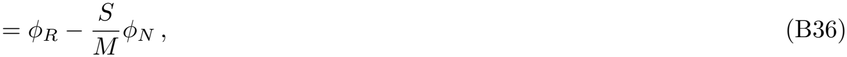

where *ϕ*_*R*_ is the fraction of non-depleted resources, and *ϕ*_*N*_ is the fraction of surviving species. In other words, *χ* measures how far the community is from the competitive exclusion limit. This *χ* is again the same as the corresponding susceptibility appeared in the static cavity method. We finally have

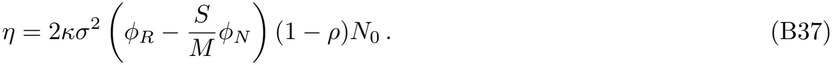

Note that when the community is fully-packed, i.e., the number of surviving species *Sϕ*_*N*_ is the same as the number of non-depleted resources *Mϕ*_*R*_, the above expression for *η* vanishes. However, the *η* term is only a linear approximation to the ecological effects. We expect that other nonlinear effects including mutation-induced extinctions [14, 15] and variances from random matrices can still contribute to the ecological suppression of fixation probabilities. These effects should still be suppressed by high *ρ*, and we show in Figs. S5 and S6 that the Kimura theory (i.e., with *η* = 0) indeed works well in the fully-packed regime when *ρ* is high. However, as *ρ* decreases, these effects can become stronger, and we would need nonlinear corrections to better predict the dynamics.

## Appendix C: Generalized consumer-resource models

We can further extend the above results to more general consumer-resource models with abiotic resources. The dynamics take the form [60]

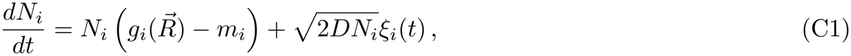

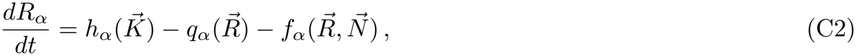

where 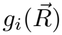 describes how the growth rate of species *i* depends on resource abundances, 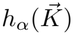 is the external supply rate of resource *α* to the community, 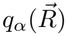represents the resource dynamics in the absence of species, and 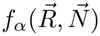 is the rate at which resource *α* is produced or consumed by the species in the community. An example of such a model is the consumer-resource model with externally supplied resources:

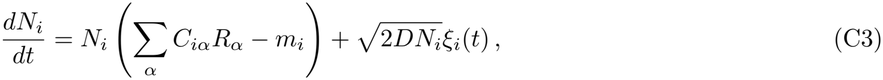

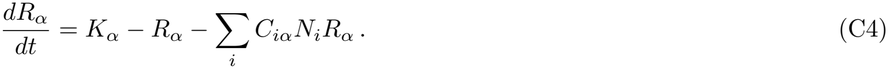

The consumer preference matrix *C*_*iα*_ matches the definition in below. The sampling schemes for *C*_*iα*_, *K*_*α*_, *m*_*i*_ are the same as in Sec. B.

Due to the generality, here we will derive the corresponding equation to Eq. (A51) in terms of the functions *g*_*i*_, *h*_*α*_, *q*_*α*_, *f*_*α*_ themselves, but not the random matrix statistics. It will be helpful to define the following matrices [60]:

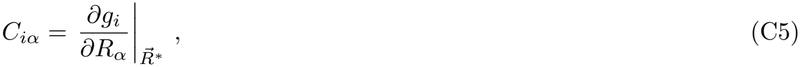

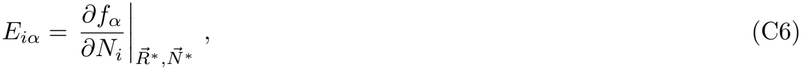

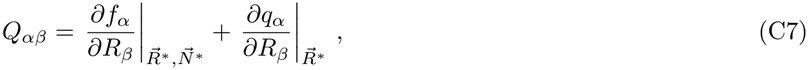

where 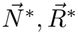 are the abundances at the steady state. *C*_*iα*_ and *E*_*iα*_ play similar roles as in the MacArthur consumer-resource model; see Eqs. (B1) and (B2). *Q*_*αβ*_ represents the interactions between resources.

Suppose at time *t* = 0, a mutation of a parent strain *p* occurs and a new mutant strain *m* with small abundance *N*_*m*_(0) ≪ *N*_*p*_(0) invades the community. The consumer-resource dynamics becomes

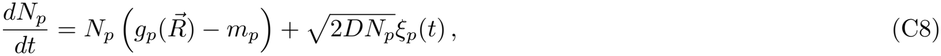

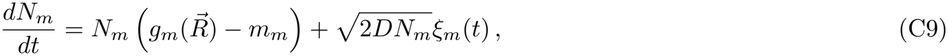

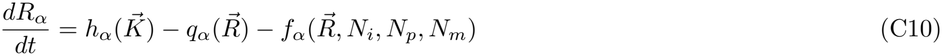

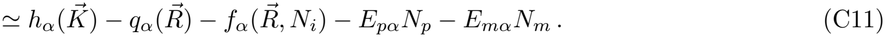

In terms of the mutant frequency *f* = *N*_*m*_*/*(*N*_*p*_ + *N*_*m*_), the above becomes

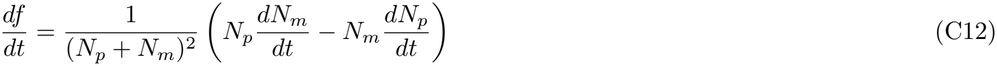

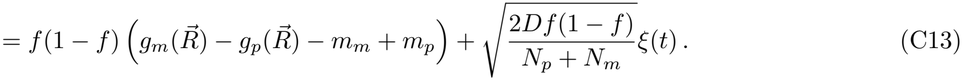

The dynamics of the other species *N*_*i*_ will not be relevant in the following calculation. Now we adopt the same set of approximations as before. We can model the changes of *R*_*α*_ as linear responses, which can be approximated by the variations of the steady state conditions:

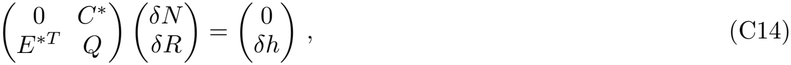

where

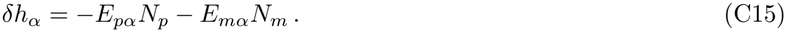

Using block matrix inversion, we have

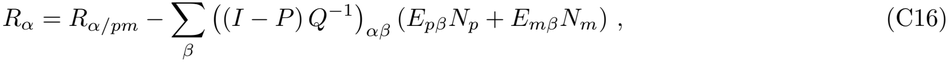

where

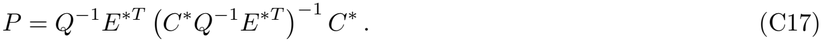

is a projector in resource space satisfying *P* ^2^ = *P* [60]. We see that the matrix (*I*−*P*)*Q*^*−*1^ generalizes the susceptibility *χ*_*αβ*_ in the MacArthur consumer-resource model. Now, the mutant frequency satisfies

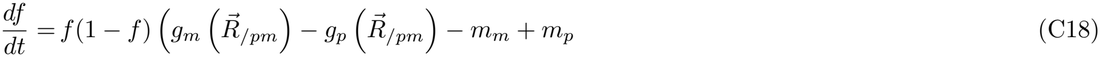

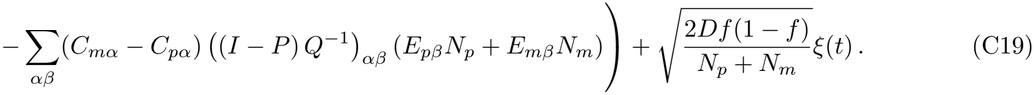

As before, we then approximate

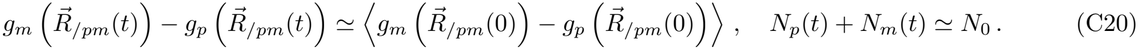

We then arrive at the same equation as Eq. (A51):

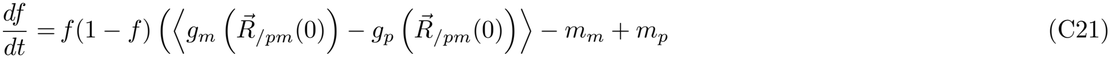

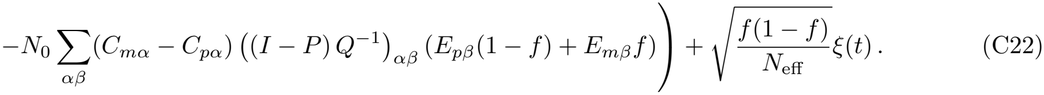

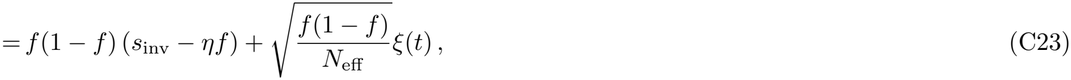

where the invasion fitness is

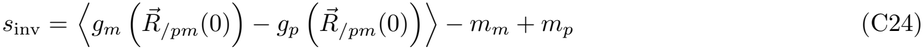

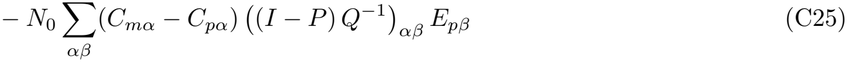

and the strength of ecological effect is

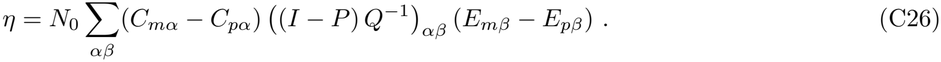

Here we do not specify the statistics of the matrices *C, E, Q*. Nevertheless, we expect that upon self-averaging, the expression of *η* involves only the trace of (*I*− *P*)*Q*^*−*1^ instead of the full matrix, corresponding to the scalar susceptibility *χ* in the MacArthur consumer-resource model. We then conclude that generalized consumer-resource models lead to the same dynamics of mutant frequency as in the MacArthur consumer-resource model.

## Appendix D: Derivation and analysis of fixation probability

From the previous sections, we have seen that all the ecological models give rise to the same dynamics of mutant frequency *f*, satisfying the stochastic differential equation

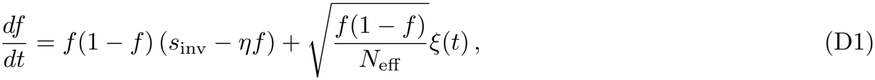

where *s*_inv_ is the invasion fitness, *η* is the strength of ecological effects depending on details of the ecological model, and *N*_eff_ is the initial effective population size of the parent and the mutant.

The fixation probability can be obtained by solving the stochastic differential equation. Let *p*(*f, t* |*x*) be the probability that the frequency is between *f* and *f* + *df* at time *t* given that *f*(0) = *x*. The corresponding Kolmogorov backward equation is

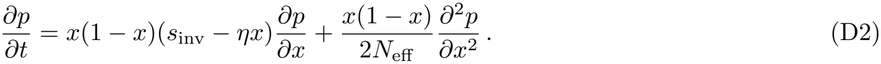

Since *f* stops at either 0 or 1 after sufficiently long time, *p* must approach the steady state *p*^*\**^(*f, x*) = *p*(*f*, ∞|*x*) given by

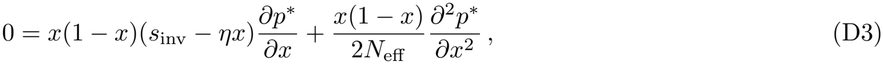

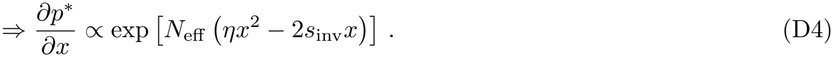

Using the stopping condition *p*(1, 0) = 0 and *p*(1, 1) = 1, we arrive at the fixation probability

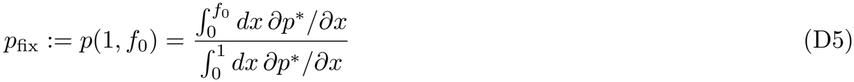

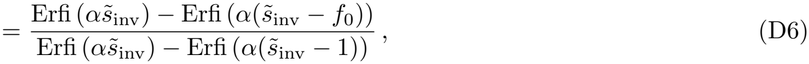

where

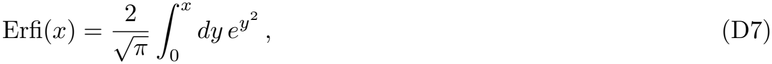

is the imaginary error function. We have denoted 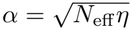 as the ratio of strength between ecological effects and demographic noise. Although *η* is suppressed by 1 − *ρ*, we have also assumed a large *N*_eff_ and it is possible that 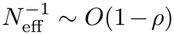 or lower, so that *α* can be large. Next, 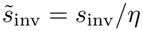 is closely related to the “dressed invasion fitness” *g*_dressed_ in [14]. 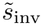predicts the frequency for the coexistence region when the parent and mutant can coexist. Meanwhile, *g*_dressed_ predicts the invader abundance when there is no invasion-induced extinction. Therefore, in the regime of parent-mutant coexistence, we have

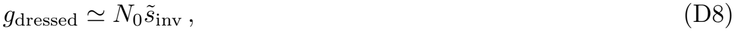

but there is no definite relationship between *g*_dressed_ and 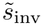outside this regime, especially when mutant invaders displace their parents.

We can better understand the formula for *p*_fix_ using the asymptotic forms

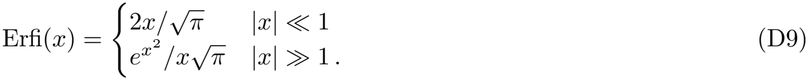

For example, when *s*_inv_ = 0,

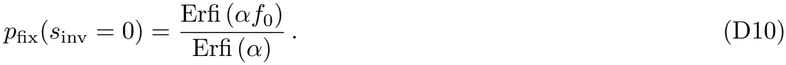

For *α* ≪ 1 (and *f*_0_ ≪ 1 as we have assumed), we can use the linear approximation for all the terms, thus

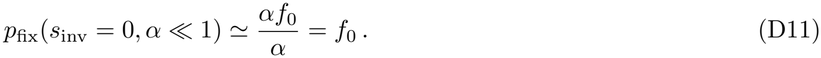

For *α* ≫1, the denominator is always exponentially large and dominating. Ignoring the subdominating factors, we have

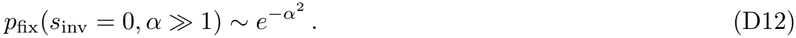

We see that *p*_fix_(*s*_inv_ = 0) matches Kimura’s prediction when *α*≪ 1, but exponentially suppressed by *α* otherwise.

The fixation probability for mutations with *s*_inv_ = 0 suggests that there is a crossover between the regimes of weak and strong ecological effects, characterized by small and large *η* respectively. First, for small *η* with *α* ≪ 1, we have 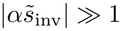 for any finite *s*_inv_ since 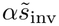scales as *η*^*−*1*/*2^. Therefore, we can use the exponential approximation for all the terms and *p*_fix_ becomes

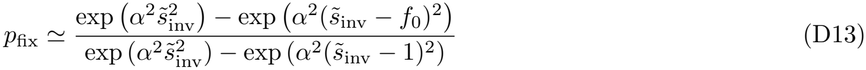

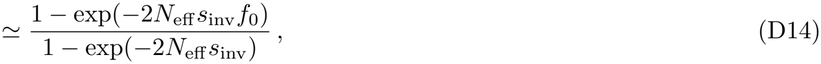

which is exactly Kimura’s formula for fixation probability with *s* = *s*_inv_. Combining the result for *s*_inv_ = 0, we see that Kimura’s formula works equally well in ecological context as long as *α* ≪ 1. On the other hand, for large *η* with *α* ≫ 1, the behavior of *p*_fix_ is divided into different regimes according to *s*_inv_. Now for 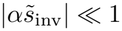, we again have

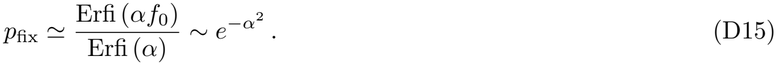

**FIG. S4:**
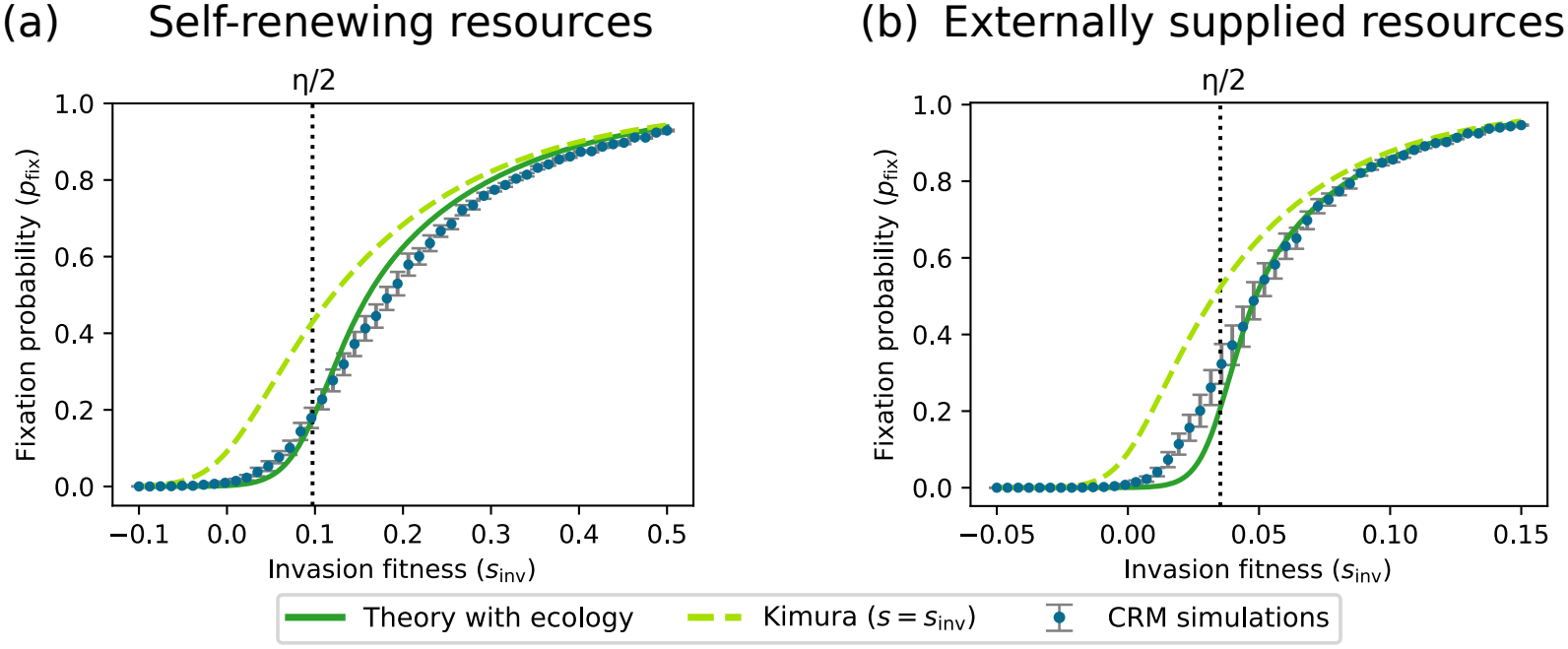
Theoretical and simulated fixation probabilities in consumer-resource models. We compute fixation probabilities *p*_fix_ at various invasion fitness *s*_inv_ by simulating mutations within consumer-resource models with (a) self-renewing resources (MacArthur consumer-resource model) and (b) externally supplied resources. In both cases, our theoretical predictions are more accurate than Kimura’s predictions, but do not fully fit with the simulation results. Error bars denote standard errors from multiple instances of demographic noise and mutants.

For 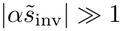, we have

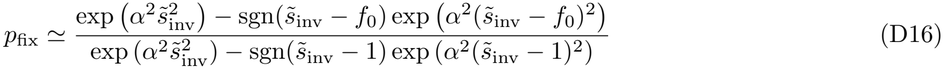

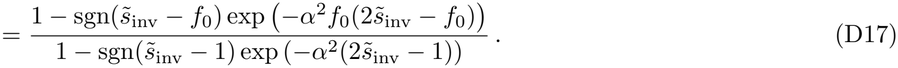

The behavior of the above expression is further divided into two regimes. When 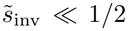, the denominator is dominating and *p*_fix_ is still exponentially suppressed:

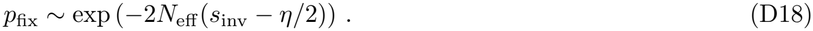

When 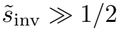, the denominator becomes very close to 1 and the numerator becomes dominating instead. We then have

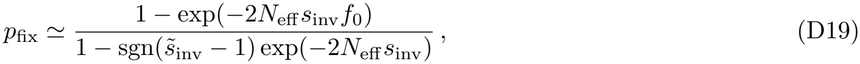

which matches Kimura’s formula approximately for 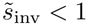 and exactly for 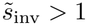.

In conclusion, while *p*_fix_ for negative *s*_inv_ is always exponentially suppressed regardless of ecology, the presence of a large *η* now also exponentially suppresses *p*_fix_ with positive *s*_inv_ up to

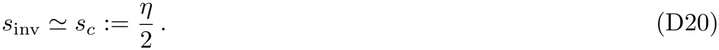

We can then interpret *s*_*c*_ as a new threshold in *s* for beneficial mutations that overcome the competitive ecological effects. In particular, the result for *s*_inv_ = 0 can now be understood as

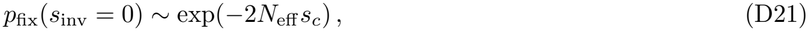

which is the same as Kimura’s prediction but for *s* = −*s*_*c*_.

## Appendix E: Fixation probabilities in consumer-resource models

In the main text, we have compared our predictions on *p*_fix_ with the simulation results from generalized Lotka-Volterra model. The comparison can similarly be done for various consumer-resource models as in Figure S4. We find that our predictions are in general more accurate than Kimura’s predictions. On the other hand, there are some deviations for 0 *< s*_inv_ *< η*, where the parent and the mutant can transiently coexist, hence the deterministic ecological dynamics affect the fixation probabilities more significantly. Compared to generalized Lotka-Volterra model, here the species interactions are mediated through resource dynamics. For self-renewing resources, there are additional complications since resources can also go extinct. We suspect that these additional complexities cause our DMFT approximations in Sec. B and C to be less accurate.

**FIG. S5:**
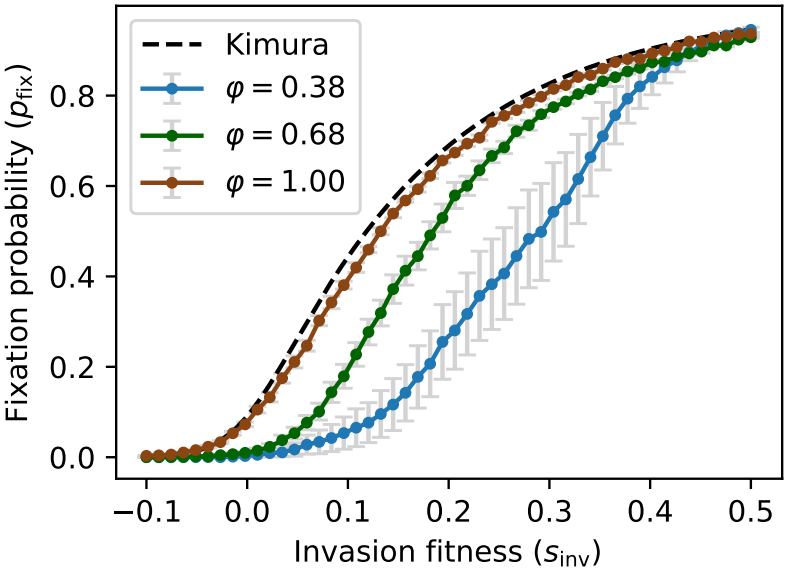
Ecological suppression of fixation probabilities controlled by species packing in consumer-resource models. We compute fixation probabilities *p*_fix_ at various invasion fitness *s*_inv_ by simulating mutations within MacArthur consumer-resource models with different packing fractions *φ* = *S*^*\**^*/M* ^*\**^. The ecological suppression of fixation probabilities is stronger when the packing fraction is lower, and there is nearly no suppression when the community is fully packed. Error bars denote standard errors from multiple instances of demographic noise and mutants.

We have also demonstrated in the main text that the ecological suppression of fixation probabilities is affected by species packing. While the packing is related to May’s stability bound for generalized Lotka-Volterra model, for consumer-resource models the packing is related to the competitive exclusion principle. The packing fraction is given by *S*^*\**^*/M* ^*\**^, where *S*^*\**^ is the number of surviving species and *M* ^*\**^ is the number of non-depleted resources. For externally supplied resources we have *M* ^*\**^ = *M*. The packing bound is *S*^*\**^*/M* ^*\**^ ≤ 1 for self-renewing resources and interestingly *S*^*\**^*/M* ^*\**^ ≤1*/*2 for externally supplied resources [73]. In Fig. S5, we indeed see that the fixation probabilities are more suppressed for a less packed community with lower packing fraction. In the other direction, we also see that the fixation probabilities nearly reduce to the results of Kimura when the community is fully packed. This observation apparently contradicts [15], which observed a sudden increase in certain coexistence probabilities near full packing in simulations (Fig. 3b in Ref. [15]). This might arise from minor differences in modeling assumptions which result in communities behaving differently when their niches are fully packed. These differences may affect the behavior of species and resource susceptibilities, with concomitant effects on invasion probabilities. An additional reason that the difference in behaviors between our models may arise is due to differences in how parameters are sampled, e.g., in Fig. S5, we have imposed specific values of *s*_inv_, in contrast with Ref. [15].

Note also that the numbers of surviving species in Fig. S5 are non-monotonic, namely 15, 26, 22 from low to high packing, showing that the number of surviving species is indeed not a good predictor for the ecological suppression, because of differences in the number of surviving resources *M* ^*\**^. The same non-monotonicity is also present in the amount of biomass.

In conclusion, we see that the results for different consumer-resource models are qualitatively the same as in generalized Lotka-Volterra model. Such agreement suggests that our theory is indeed applicable to various ecological models.

## Appendix F: Effects of parent-mutant correlation on fixation probabilities

Throughout the paper, we have assumed high parent-mutant correlation *ρ*≈ 1. As shown in Appendix A, this is one of the crucial assumptions that allows us to simplify the mutation dynamics into the form in Eq. (A51). Apart from the justifications made in Appendix A, here we also validate the small 1 −*ρ* expansion using the downstream quantities, namely the fixation probabilities. In Fig. S6, we see that there are indeed ranges of small 1 −*ρ* in which our approximated theory in Eq. (D5) agrees well with full simulations. That is, the fixation probabilities decrease with smaller *ρ* due to frequency-dependent selection. In particular, both theory and simulations reduce back to Kimura’s results as *ρ* approaches 1. The agreement remains true even for fully-packed communities (Fig. S6c, d) in which nonlinear effects become important (see the end of Appendix B). We also see that our theory starts to deviate from the simulations when *ρ* is no longer large, though the specific value of *ρ* at which the deviation rises depends on other properties of the community.

**FIG. S6:**
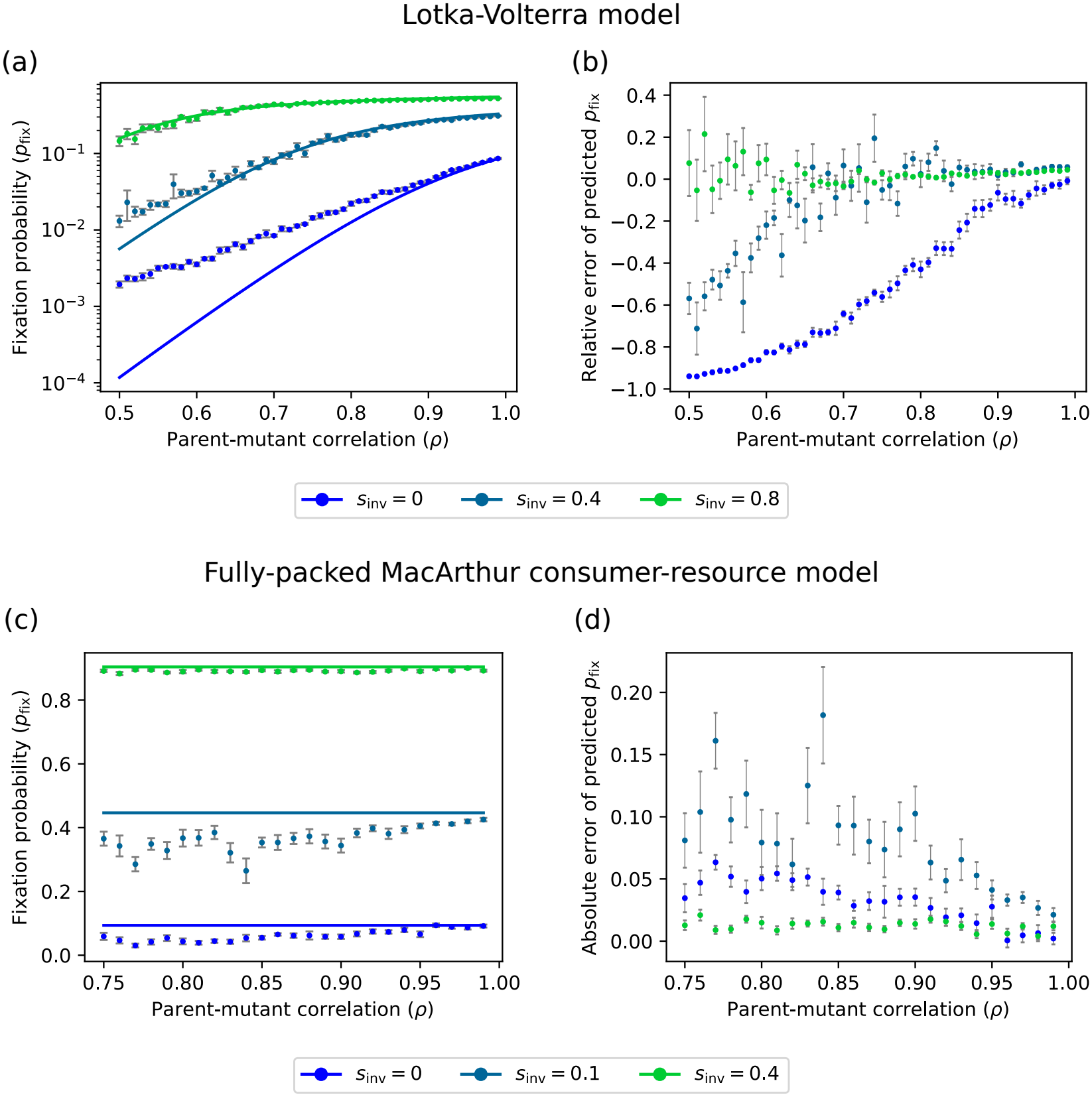
Theoretical and simulated fixation probabilities in generalized Lotka-Volterra model and MacArthur consumer-resource model at various parent-mutant correlation. We compute fixation probabilities *p*_fix_ at various invasion fitness *s*_inv_ by simulating mutations within generalized Lotka-Volterra model (a–b) and fully-packed MacArthur consumer-resource model (c–d). Our theory (solid curves) matches the simulations (dots) for high *ρ*, but deviates from the simulations for low *ρ* (a, c). The error is more significant for lower *s*_inv_ (b, d). Error bars denote standard errors from multiple instances of demographic noise and mutants.

## Appendix G: Comparison between selection and drift dynamics

In this appendix, we obtain a qualitative understanding of both the results in classic population genetics and our new results, by comparing the relative sizes of selection and drift terms in the differential equation for *f*. This analysis helps us understand how parent-mutant coexistence arises in the presence of ecological effects.

### 1. Without ecological effects

For pedagogical purposes, it is helpful to start by re-deriving some classic results from classical population genetics with constant selection (in the absence of many-species ecology). As explained in the main text, a mutant starts to invade at a very small frequency where genetic drift dominates. The mutant can either become extinct or grow to a larger frequency due to drift. This process continues until the surviving mutants reach a frequency *f*∼ 1*/N*_eff_ |*s*|, where selection becomes dominant instead. This critical frequency is the drift threshold as shown in Fig. 4a in the main text.

We now obtain a heuristic analytical estimate of drift threshold, following the approach in [97, 98]. Recall that the dynamics of *f* in classic population genetics is given by

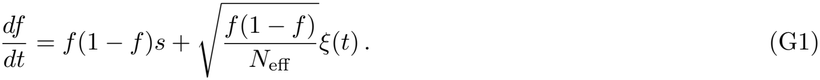

The exact solution to this differential equation is in general complicated as both the growth rate and diffusion rate depends on *f*. On the other hand, analogous to the Euler method for numerically solving differential equations, we can approximate the dynamics of *f* by splitting it into infinitesimal time intervals with width Δ*t*. Within a time interval around some time *t*, the changes in both the growth rate and the diffusion rate can be neglected. Therefore, the change in frequency Δ*f* within the time interval can be easily solved and is given by

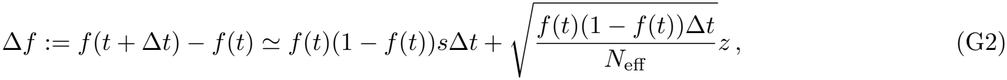

where *z* is a zero mean, unit variance Gaussian random variable. Below, we can simply use *f* to denote *f*(*t*).

Let us focus on small frequencies *f* ≪1 where drift dominates over selection. In this case, Δ*f* can be either positive or negative due to randomness, and its magnitude can be estimated by the variance

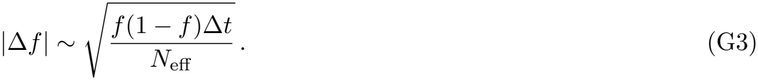

The above approximation of constant growth and diffusion rates remains valid until Δ*f* is comparable to *f*, that is

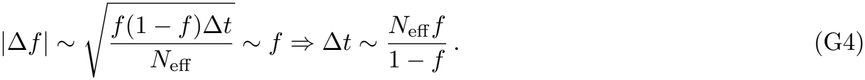

Within this time interval, the contribution to Δ*f* from selection is

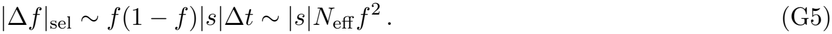

Note that the absolute value is needed since Δ*f* due to selection can also be either positive or negative depending on the sign of *s*. To ensure that the approximation is self-consistent, Δ*f* due to selection must be negligible, i.e., |Δ*f*|_sel_ ≪ |Δ*f*|. Therefore, we require

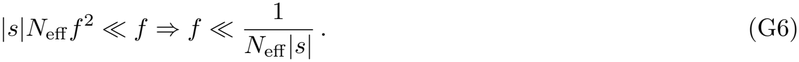

Indeed, we see that drift dominates the dynamics when *f* is below drift threshold.

### 2. With ecological effects

We can repeat the above analysis under the presence of ecological effects. As explained in the main text, when 0 ≲ *s*_inv_ ≲ *η*, there is an additional deterministic fixed point 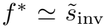. We then expect a new region around both sides of *f* ^*\**^ where drift dominates the dynamics, apart from the one near *f* = 0. This is because at every fixed point, by definition, the deterministic contribution to the dynamics vanishes.

Suppose the mutant reaches frequency *f* ≫ *f*_0_ at time *t*. Again, we focus on an infinitesimal time interval with width Δ*t*. From Eq. (A51), we can approximate Δ*f* within this time interval as

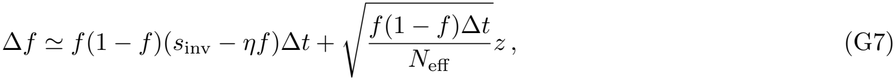

where *z* is a zero mean, unit variance Gaussian random variable. Note that the variance is the same as in the case without ecological effects. Therefore, if drift dominates the dynamics, the above expression of Δ*f* remains a good approximation until

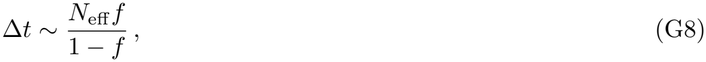

which is the same as Eq. (G4). To ensure that this approximation is self-consistent, the contribution to Δ*f* from selection must be negligible. Therefore, we require

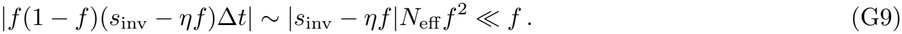

**FIG. S7:**
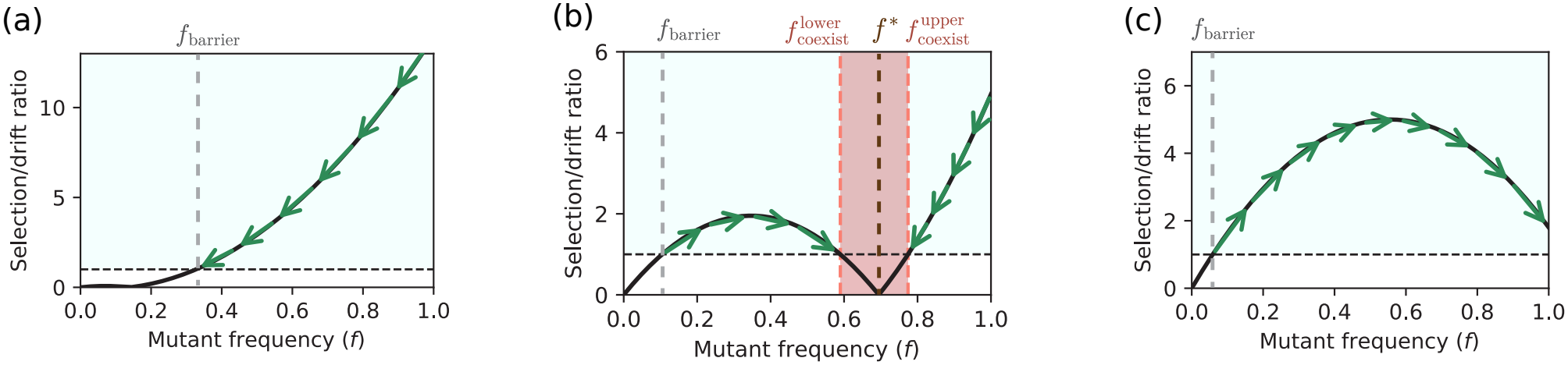
Phase portraits of frequency dynamics in terms of selection-drift ratio. The selection-drift ratios *R* (Eq. (G10)) are calculated in different regimes of invasion fitness *s*_inv_, namely 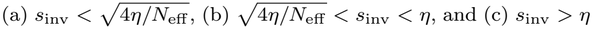. When *s*_inv_ is in the intermediate range, there are more crossovers between selection and drift dominated regimes, hence a coexistence region centered at *f* ^*\**^ emerges. The selection-drift ratios are identical to the ones in Fig. 3 in the main text.

Note again that the absolute value is needed since frequency can either increase or decrease, i.e., Δ*f* due to selection can be either positive or negative, particularly when *f* is around the new fixed point 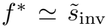, which is between *f* = 0 and *f* = 1. Based on the above condition, we can define the selection-drift ratio

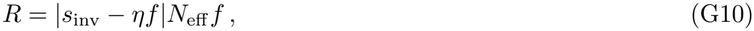

such that drift (selection) dominates the dynamics when *R <* 1 (*R >* 1), and *R* = 1 is the crossover boundary between selection and drift dominated regimes. This is the ratio as shown in Fig. 3f-h in the main text.

Since we already know that our results reduce back to those in population genetics when *α* ≪ 1 (*η* ≈ 0), we now focus on the case of large *η* and *α* ≫ 1. In this regime, due to the absolute value in *R*, the solutions to *R* = 1 have different types of behavior depending on the value of *s*. Recall that we require 0 ≤ *f* ≤ 1. When 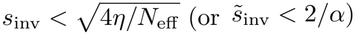, there is only one solution with *s*_inv_ − *ηf <* 0: (Fig. S7a)

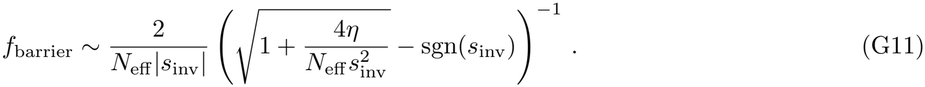

When *s*_inv_ *> η*, there is also only one solution but with *s*_inv_ − *ηf >* 0: (Fig. S7c)

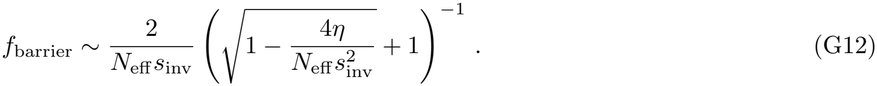

Therefore, the dynamics of *f* is qualitatively the same as the one in classic population genetics for 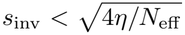 or *s*_inv_ *> η*, with *f*_barrier_ being the drift threshold.

In contrast, the dynamics of *f* is qualitatively distinct for 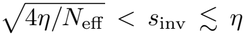. We see that there are three solutions to *R* = 1: (Fig. S7b)

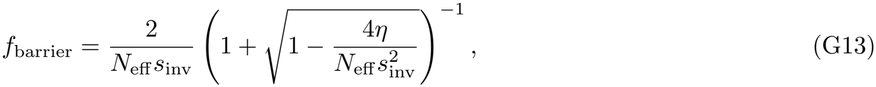

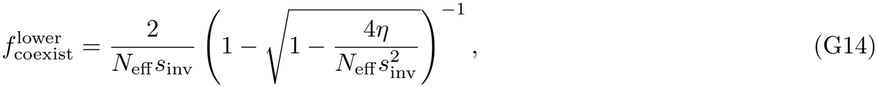

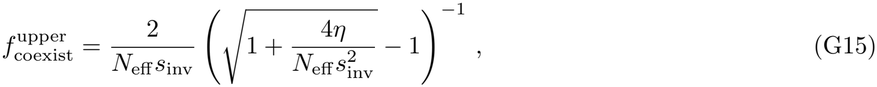

with 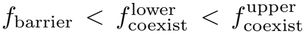 . Note that we have 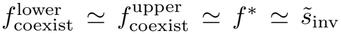, hence these two frequencies correspond to the bounds of the coexistence region centered at *f* ^*\**^.

As demonstrated in the main text, our results in fixation probability and mean absorption/extinction time can be qualitatively understood using the nonlinear behavior of *R* (Fig. S7b). Suppose a mutant starts with *f* = *f*_0_≪ 1 at the beginning. It must overcome the first barrier at *f* = *f*_barrier_ or it becomes extinct. After that, selection brings the mutant to the region 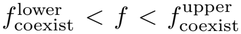, which is the coexistence region. Note that the deterministic fixed point 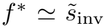 is within this region. Since 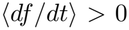 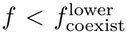 and ⟨*df /dt*⟩ *<* 0 when 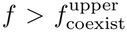, both boundaries of the coexistence region are reflecting and the mutant is trapped inside this region. It can escape this region and reach near the absorbing boundaries *f* = 0, 1 only when the demographic noise reaches the tail of its distribution with exponentially low probability.

Now we see that if *s*_inv_ *< s*_*c*_ = *η/*2, the coexistence region is closer to *f* = 0 and it is exponentially more likely for the demographic noise to drive the mutant to extinction. Therefore, *p*_fix_ is exponentially suppressed for *s*_inv_ *< s*_*c*_. In contrast, when *s*_inv_ *> s*_*c*_, the coexistence region is closer to *f* = 1 and it is exponentially more likely for the demographic noise to drive the mutant to fixation. The mutant becomes extinct mostly because it cannot overcome the first barrier at *f* = *f*_barrier_, hence *p*_fix_ agrees with Kimura’s formula for *s*_inv_ *> s*_*c*_.

The above argument also explains the behavior of mean absorption/extinction time. It grows and decays exponentially in the region 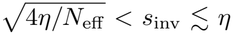 due to the required time to escape the coexistence region. More precisely, the time grows exponentially as *s*_inv_ increases and the coexistence region becomes farther from *f* = 0. The growth continues till *s*_inv_ = *s*_*c*_. For *s*_inv_ *> s*_*c*_, the coexistence region becomes closer to fixation, and the mean absorption time decays exponentially as fixation becomes more likely. For the mean extinction time, although the time for escaping from the coexistence region to *f* = 0 is even longer, it also becomes exponentially unlikely that the mutant becomes extinct after reaching the coexistence region. Namely, a mutant becomes extinct mainly due to the first barrier at *f* = *f*_barrier_. As a result, the mean extinction time reaches maximum at *s*_inv_ = *s*_*c*_ and decays exponentially for *s*_inv_ *> s*_*c*_.

## Appendix H: Mean extinction time

Using the above qualitative picture of coexistence region, we can derive analytical estimates for the mean extinction time of the mutant. It is also similar to the establishment time in population genetics, which is the time when selection dominates over drift and drives a mutant with positive *s* to fixation. As explained in the main text, the extinction or establishment time is less sensitive to model details and is easier for analytical estimation.

For the regimes without the coexistence region, i.e. 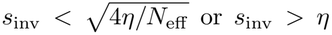, the extinction tim e can be approximated by the time required to reach the barrier at *f* = *f*_*R*=1_; see Eqs. (G11) and (G12). 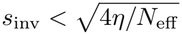,the barrier is reflecting and the mutant becomes extinct in the same order of magnitude of time after reaching the barrier. For *s*_inv_ *> η*, the barrier is absorbing and the mutant can no longer become extinct after passing through the barrier. Now, the time for reaching the barrier can be estimated using Eq. (G4); since Δ*t* increases with *f*, the required time to reach *f* = *f*_*R*=1_ is dominated by Δ*t*(*f*_*R*=1_). Therefore, for 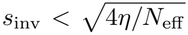, the mean extinction time *T* is

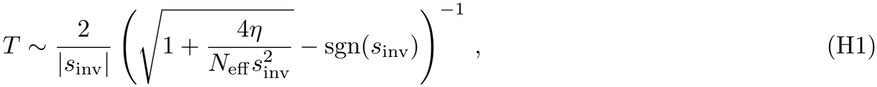

and for *s*_inv_ *> η*,

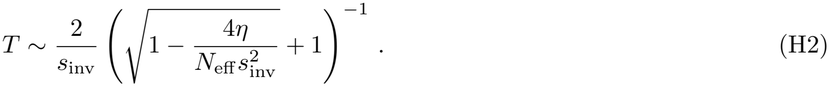

Both cases match the expectation from population genetics that *T* ∼ 1*/* |*s*_inv_| asymptotically.

On the other hand, the mean extinction time deviates from the expectation from population genetics when the coexistence region is present, i.e. when 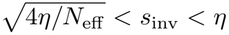. In this regime, the mean extinction time is dominated by the exponentially long time to escape the coexistence region driven by the stochastic dynamics. The scale of the escape time can be estimated by translating the stochastic dynamics into 1D diffusion in the presence of a potential *V* (*f*). Since *f* relaxes to near equilibrium around the deterministic fixed point 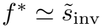 in the long term (but before being absorbed at *f* = 0, 1), the probability distribution *p*(*f*) can be obtained from the Fokker-Planck equation:

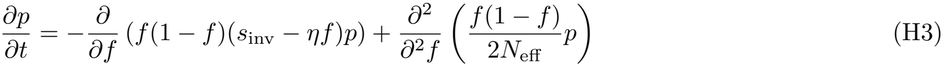

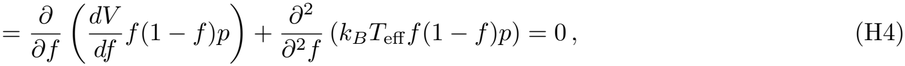

where we have used the language of statistical physics and defined a potential for *f* (with minimum at *f* = *f* ^*\**^)

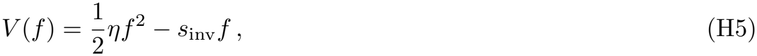

and an effective temperature

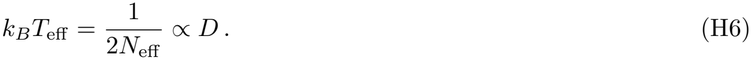

As long as *f* is not absorbed at 0 or 1, the solution involves a Boltzmann distribution [99]

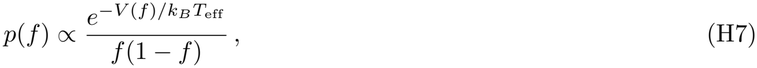

For large *η* with *α*≫ 1, we can focus on the exponential factor and approximate the escape time as 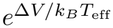 according to Kramer’s formula, where Δ*V* is the barrier height in the potential. In particular, the escape time to extinction is

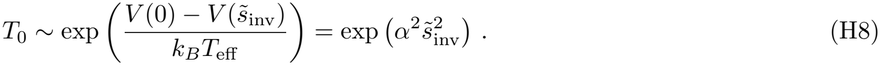

However, *T*_0_ is conditional on that the mutant reaches near equilibrium around *f* ^*\**^ and does not eventually escape to fixation. When *s*_inv_ *> s*_*c*_ = *η/*2, this is no longer the main route to extinction since the coexisting region is closer to fixation as in Fig. S7b. Instead, almost all extinctions occur below the drift threshold, i.e., *f < f*_barrier_ in Fig. S7, with much shorter extinction time *T*_*b*_∼ 1*/s*_inv_ according to population genetics. As explained in the main text, the probability for a mutant to pass through the drift threshold is precisely Kimura’s fixation probability

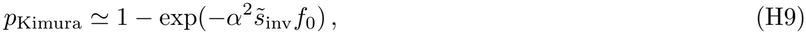

for large *α* and *s*_inv_ *> s*_*c*_. Note that the exponent is much smaller than the one in *T*_0_ since *f*_0_ is small. Meanwhile, conditional on passing through the drift threshold, the probability for a mutant to reach *f* = 0 instead of *f* = 1 is

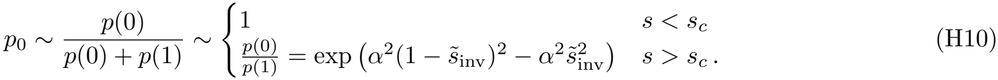

Therefore, the mean extinction time is

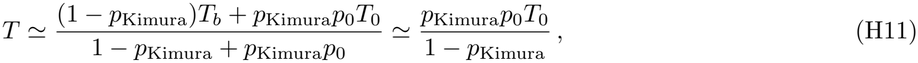

since *p*_0_*T*_0_ is still exponentially larger than *T*_*b*_, while 1− *p*_Kimura_ is less suppressed than *p*_0_. Taking the leading exponents, we arrive at

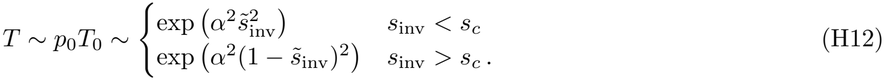

In particular, the mean extinction time reaches maximum when *s*_inv_ = *s*_*c*_ ≃ *η/*2, i.e.

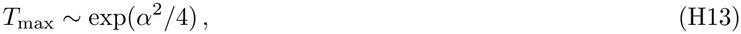

which is determined by *α* only. Importantly, *T* is still exponentially suppressed by the exponent of 1 −*ρ* compared to the stochastic extinction time of a whole species. Therefore, we can indeed observe the stochastic extinction or fixation of mutants well before the community collapses from its steady state.

Apart from the exponential behavior of *T*, it is useful to estimate its prefactor, which has the unit of time. In the ordinary Kramer’s theory, where both the potential well (let say at *f* = *f*_*a*_) and the barrier (at *f* = *f*_*b*_) contain saddle points, the prefactor is roughly

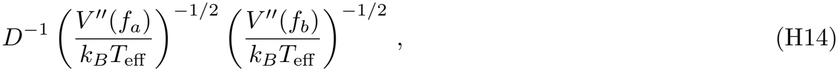

where the second derivative factors come from the normalization of the Boltzmann distribution. However, in our case the barrier contains an absorbing boundary at *f* = 0 instead of a saddle point. Its second derivative factor is then replaced by a first derivative factor [100], hence the prefactor of *T* is

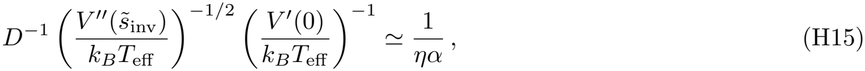

where we have used the fact that *s*_inv_ ∼ *η* for parent-mutant coexistence. Therefore,

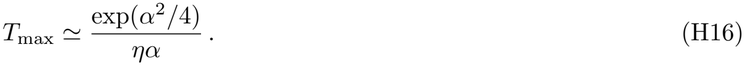

**FIG. S8:**
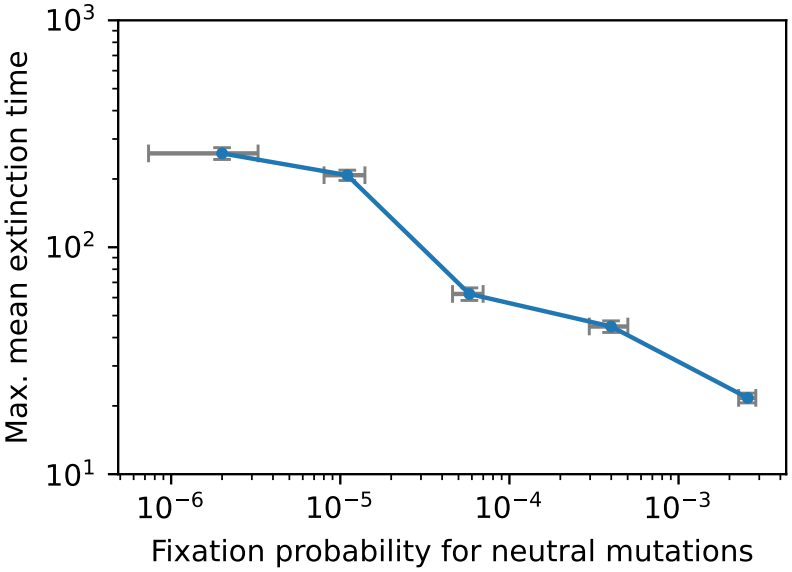
Correlation between fixation probabilities and mean extinction times. In log scales, there is negative correlation between the fixation probability for mutations with *s*_inv_ = 0 and the maximum mean extinction time. Error bars denote standard errors from multiple instances of demographic noise and mutants.

This expression can be directly compared to the typical time for a new mutation *T*_mut_ ≃1*/N*_ind_*U*, where *N*_ind_ is the number of individuals of a species and *U* is the mutation rate of an individual. The “strong-selection-weak-mutation” limit holds when *T*_max_ *< T*_mut_.

It is interesting to observe that there are two independent quantities that are related to 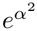, namely the fixation probability for mutations with *s*_inv_ = 0 and the maximum mean extinction time. In particular, we expect that the maximum time is exponentially longer as the fixation probability is exponentially suppressed, since both exponential behavior is caused by parent-mutant coexistence as explained in the main text. Fig. S8 shows that the expectation is indeed realized in simulations.

## Appendix I: Details of simulation methods

We use simulations to compute the fixation probabilities and other statistical quantities within the full ecological models. The code for the below simulation and the associated figures in this paper can be found on *https://github.com/Emergent-Behaviors-in-Biology/Pop-Gen-Ecology*.

### 1. Community dynamics

First, we sample the ecological properties of the community as described in Sec. A and B. We then evolve the community towards its unique steady state *without demographic noise* by solving the full differential equations using the LSODA method (with absolute and relative tolerances 10^*−*12^). The convergence is reached when all the derivatives *dN*_*i*_*/dt, dR*_*α*_*/dt* are less than 10^*−*10^. We can then use these results to compute the predicted values of *η*.

After that, since we only focus on a single step of mutation, we choose one of the surviving species as the parent.

Then, we look at multiple instances of the mutation with the same parent. Each time, we introduce a mutant associated with the chosen parent, sampled as described in Sec. A and B. We then solve the full differential equations *with demographic noise* using a generalized Euler method as explained below. Due to the multiplicative nature of the noise, the simulation can become numerically unstable if we sample the noise directly. Instead, we make use of the result in [94]. Within a time step *dt*, we can approximate that the total growth rates of the species remain unchanged, since the growth rates involve averages over all the species or resource abundances. In other words, the species dynamics within time *dt* becomes

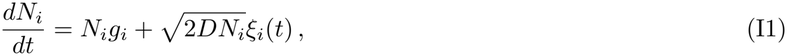

where *g*_*i*_ is the momentarily constant growth rate depending on the current species or resource abundances. It was noticed that the above equation can be solved exactly, and the solution is given by

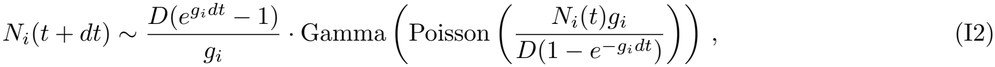

where Gamma(*α*) is the gamma distribution with shape parameter *α* and unit scale. We define Gamma(0) as the Dirac delta distribution at 0. For each time *t*, we sample *N*_*i*_(*t* + *dt*) according to the above distribution. If resources are involved in the model, their abundances are updated at the same time using the ordinary Euler method. After that, to ensure uninvadability, we impose a hard wall for the abundances at *λ* = 10^*−*7^, i.e.

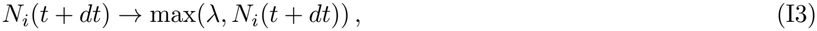

and similarly for resources if present. We iterate the above process with a time step *dt* = 0.1 till the parent or mutant abundance reaches *λ*, which is treated as extinction.

The parent and mutant trajectories in classic population genetics are simulated in the same way except that *g*_*i*_ are now truly constants and the fitness difference is *s* = *g*_*m*_ − *g*_*p*_.

### 2. Susceptibility

Recall the definition of susceptibility in the GLV in Eq. (A15). Following [101, 102], we use the likelihood-ratio estimation method to numerically compute the susceptibility function; here we provide the full derivation of the method for completeness. Due to the stochastic nature of our simulation, we need not perturb *r*_*i*_ and rerun the noisy simulation to calculate the derivatives. This is because the rerun would result in the same possible set of trajectories but weighted by a different probability measure based on *δr*. Therefore, we instead apply a change of measure to the original unperturbed trajectories:

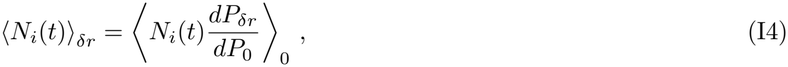

where the subscript 0 represents the unperturbed measure. By the Girsanov’s theorem,

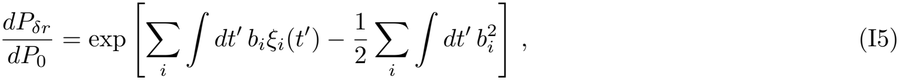

where

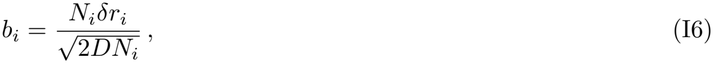

is the ratio between the deterministic perturbation *N*_*i*_*δr*_*i*_ and the stochastic standard deviation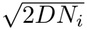 The suscep-tibility can then be obtained by taking the derivative on *δr*.

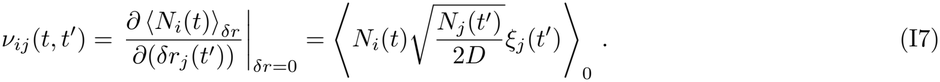

In the simulation, we reconstruct *ξ*_*j*_(*t*^*′*^) using Eq. (I1):

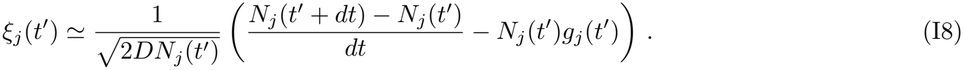

In conclusion, we can numerically compute the susceptibility using the formula

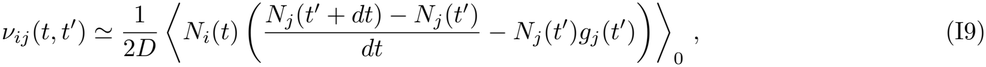

applied on the unperturbed trajectories.

### 3. Fixation probabilities and times

To obtain fixation probabilities, we sample 10 mutants with the same chosen parent. For each mutant, we adjust the value of *r*_*m*_ (for generalized Lotka-Volterra model) or *m*_*m*_ (for consumer-resource models) so that the invasion fitness is at a given value of *s*_inv_. We then run the above simulation 10^3^ times with different instances of demographic noise and count the number of fixations. We take the average of the above results to find the fixation probability *p*_fix_(*s*_inv_).

To obtain absoprtion and extinction times, we sample only one mutant for the chosen parent, then run the simulation 10^3^ times and record the time to fixation or extinction for each run.

## Appendix J: Simulation parameters used in the figures

Throughout all simulation figures, the initial mutant frequency is *f*_0_ = 0.09. We set *A*_*pm*_ = *A*_*mp*_ = *ρ* for all simulations of generalized Lotka-Volterra models.

In Fig. 1(f) and (h), we simulate the dynamics as in classic population genetics with the parent growth rate *g*_*p*_ = 0.1, the parent abundance *N*_*p*_ = 5.0, the fitness difference *s* = 0.01, and the diffusion coefficient *D* = 0.5. In Fig. 1(g) and (i), we simulate generalized Lotka-Volterra model with parameters

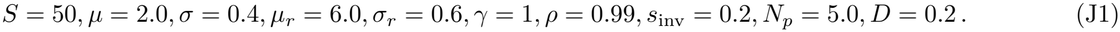

In all the subfigures in Fig. 2, we simulate generalized Lotka-Volterra model with parameters

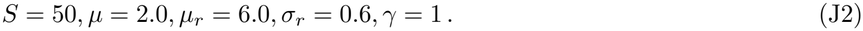

In Fig. 2(a) and (b), we further use the parameters

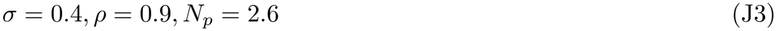

We change the control parameter *α* by using *D* = 1.0 in Fig. 2(a) and *D* = 0.04 in Fig. 2(b). In Fig. 2(e), we use

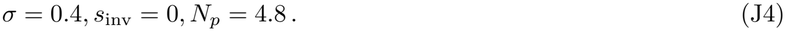

In Fig. 2(f), the simulation parameters for the case of low packing are the same as those for Fig. 2(b). To control the extent of packing, we keep all the parameters unchanged except using *σ* = 0.7 for the case with high packing. While *σ* also controls the ecological contribution in *s*_inv_, it is accordingly balanced by shifting *r*_*m*_, so that the two communities can be compared at given *s*_inv_.

In Fig. 3(a) and (b), we simulate generalized Lotka-Volterra model with parameters

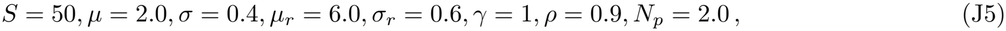

and *D* = 0.020, 0.024, 0.036 corresponding to the three values of *N*_eff_ . We focus on *D* = 0.024 in Fig. 3(c)-(h). The selection-drift ratios in Fig.3 are calculated using Eq. (G10) with the predicted values of *η*.

In Fig. 4(a), we simulate the dynamics as in classic population genetics with parameters

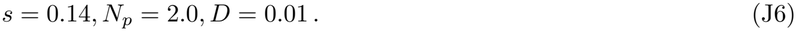

In Fig. 4(b) and (c), we simulate generalized Lotka-Volterra model with parameters

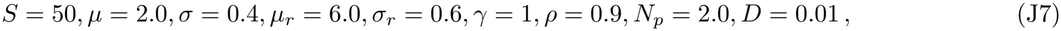

and *s*_inv_ = 0.12, 0.23 in Fig. 4(b) and (c) respectively.

In Fig. S1, we simulate generalized Lotka-Volterra model with parameters

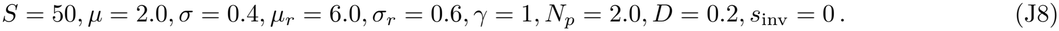

Figs. S2 and S3 use the same parameters with *ρ* = 0.9, except that *s*_inv_ = 0.1.

In Fig. S4(a), we simulate MacArthur consumer-resource model with parameters

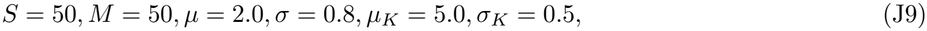

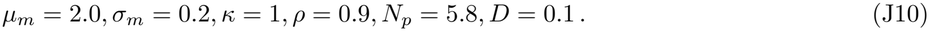

In Fig. S4(b), we simulate the consumer-resource model with externally supplied resources with parameters

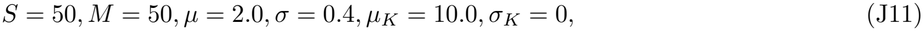

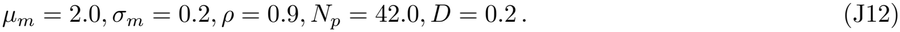

In Fig. S5, the simulation parameters for the case of *S*^*\**^*/M* ^*\**^ = 0.68 are the same as in Fig. S4. To control the extent of packing, we keep all the parameters unchanged except using *S* = 20 for the case of *S*^*\**^*/M* ^*\**^ = 0.38, and (*µ*_*m*_, *σ*_*m*_) = (0.2, 0.02) for the case of *S*^*\**^*/M* ^*\**^ = 1.

Fig. S6(a) and (b) use the same parameters as in Fig. S1 with *ρ* = 0.9, except that *s*_inv_ = 0, 0.4, 0.8. To obtain the exponentially suppressed fixation probabilities, we repeat the simulation for each mutant 10^4^ times to count the number of fixations. Fig. S6(c) and (d) use the same parameters as in the case of *S*^*\**^*/M* ^*\**^ = 1 in Fig. S5, with *s*_inv_ = 0, 0.1, 0.4.

In Fig. S7, the selection-drift ratios are identical to the ones in Fig. 3.

In Fig. S8, the simulation parameters are the same as in Fig. 3(a) and (b), except that we now use *D* = 0.02, 0.024, 0.03, 0.036, 0.05. To obtain the exponentially suppressed fixation probabilities, we repeat the simulation for each mutant 10^5^ times to count the number of fixations.

## Appendix K: Details of experimental data analysis

We analyze the longitudinal sequencing data in [75] from the gut microbiome of a single patient. Here, we provide the details including the fitted model, the procedures of maximum likelihood estimation, the inference results with bootstrapping, and the procedures for testing the goodness of fit. The dataset, the code for data analysis, and the detailed fit results can be found on *https://github.com/Emergent-Behaviors-in-Biology/Pop-Gen-Ecology*.

### 1. Data and model

First, we download the dataset from Supplemental Data S1–S3 in [75]. The dataset contains time series data for sweeping SNV clusters (with frequencies ranging from *f <* 0.2 to *f >* 0.7) from 17 abundant species in the same microbiome. For each cluster, the read count of the mutant allele *A*_*t*_ and the total read count *D*_*t*_ are reported for each of the observed time points among the 19 sampled time points. Note that a species may have multiple SNV clusters, but they may not correspond to different mutant strains. We make a simplifying but biologically reasonable assumption [7, 75, 77, 79] that different clusters within the same species are proxy for the dynamics of a single mutant.

For each of the periods in Fig. 5, we fit the model of frequency-dependent selection in Eq. (A51) to the mutant trajectories. However, Eq. (A51) only describes the dynamics of frequencies *f*(*t*), and we need an additional model of measurement noise to convert *f*(*t*) to the read counts *A*_*t*_. First, the read sampling noise in *A*_*t*_ given a true *f* is described by a binomial distribution *A*_*t*_ ∼ *B*(*D*_*t*_, *f*).

Moreover, there may be various errors from the data source itself during DNA extraction from samples, PCR amplification, and sequencing. In previous work [103, 104], these errors were typically modeled as random Poisson noise. It was because they largely focused on trajectories with very low frequencies, hence very small populations in which counting errors dominate. However, we specifically focus on sweeping trajectories with intermediate frequencies and large populations, in which multiplicative errors (such as in amplification factors) dominate instead. Therefore, we assume log-normal errors for these factors and let the sampled abundances be

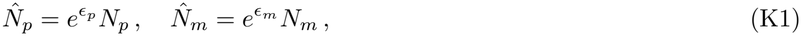

where *ϵ*_*p*_, *ϵ*_*m*_ are random Gaussian. Note that by applying the logit transform

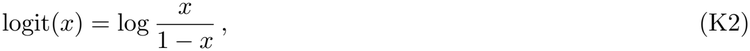

the sampled frequency 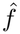 becomes

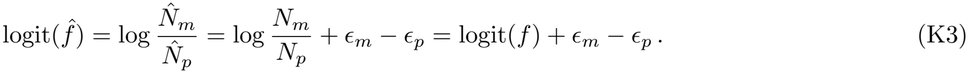

Therefore, it is equivalent to assume a random Gaussian noise on logit(*f*(*t*)) with strength *σ*. To summarize, we assume that *A*_*t*_ follows the distribution

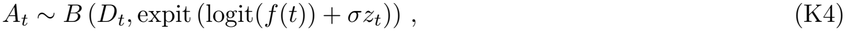

where *f*(*t*) follows Eq. (A51) and *z*_*t*_ are standard Gaussian random variables. The function

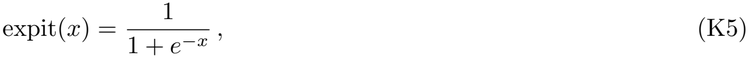

is the inverse of the logit transform.

As described in the main text, we assume shared values of *η, N*_eff_, *σ* across species since they belong to the same community, but different invasion fitnesses *s*_inv_ for each species. This is because while we mainly focus on pooled estimates of the average parameters, we expect *s*_inv_ to be more heterogeneous and possibly change sign across species. A distribution of *s*_inv_ is also necessary for predicting the extent of coexistence in the community.

To model the distribution of *s*_inv_, we make the simplifying assumption that the invasion fitness of each species is drawn from a Gaussian distribution with some mean *µ* and variance *τ* ^2^, which we must infer from the data. That is,

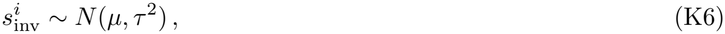

where *i* denotes the species. However, as shown later, our results are largely insensitive to the choice of a Gaussian distribution.

### 2. Likelihood function

We fit the above model to the trajectory data using maximum likelihood estimation (MLE). This procedure asks which combinations of the model parameters make the observed data most probable. The crucial part of this procedure is to compute the likelihood function of the above model, which is a function of the model parameters. Given the model parameters, the function outputs the joint probability for the observed data. The fit is then obtained by maximizing the likelihood function.

We shall compute the likelihood function using filtering algorithms. For an observed trajectory *u* with reads 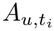at times *t*_*i*_, the log-likelihood function conditioned on *s*_inv_, denoted by *l*_*u*_(*θ, s*_inv_) (where {*θ* = *η, N*_eff_, *σ}* is the set of model parameters) can be decomposed into the log-likelihoods for each time point under one-step-ahead predictions

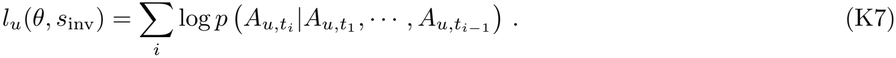

Each term in the above can be computed as follows. To propagate from 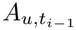 to 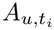, we first need to convert them into the latent frequency variables 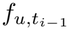and 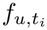 . For this purpose, in the algorithm we maintain a posterior distribution of 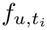

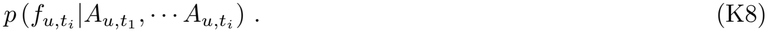

Using this posterior, we propagate from 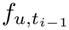 to 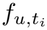 using the dynamics by

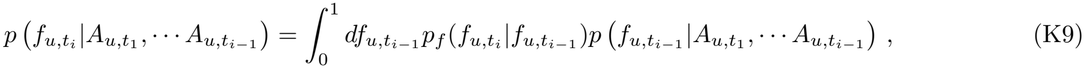

where *p*_*f*_ is the probability distribution of *f* from the dynamics in Eq. (A51) with absorbing boundaries at *f* = 0, 1, obtained from integrating forward the Fokker-Planck equation in Eq. (H3) (see below for detail). Finally, we sample 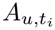 using the measurement noise model by

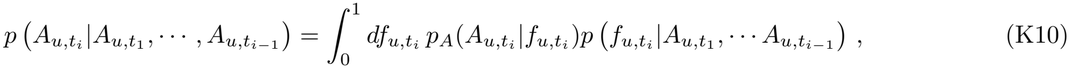

where

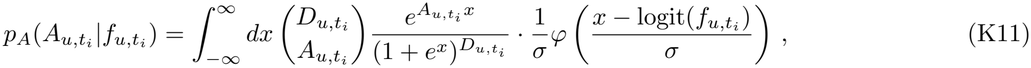

is the probability distribution of *A* from the sampling in Eq. (K4),

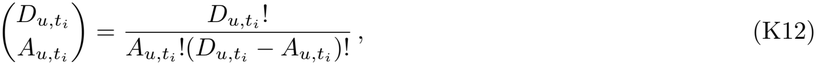

is the binomial coefficient, and

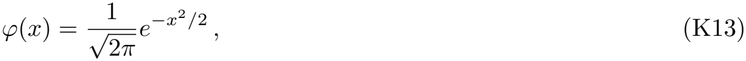

is the standard normal PDF. We first numerically find the unique peak of the integrand, then compute the integral around the peak till the integrand decays by a factor of *e*^*−*40^. Note that the above is undefined at the two absorbing boundaries; we additionally define

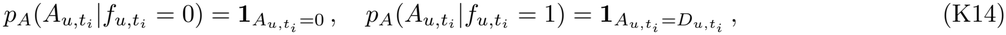

where

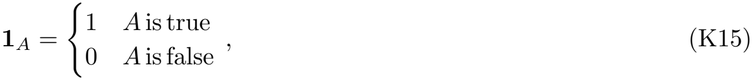

is the indicator function. To apply this algorithm, we first need to choose an initialization for the posterior 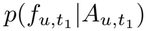. For simplicity, we assume a uniform prior on *f* before any observation, hence

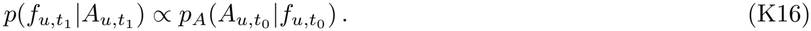

As the algorithm goes, we recursively update the posterior by

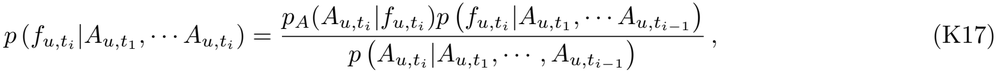

which can be computed according to the above using the posterior at the previous time point. To perform all the above integrations in frequencies, we split the space of *f* into *N* = 400 cells. Taking account of the absorbing boundaries, the fit would be more accurate with higher resolution of points near *f* = 0, 1. For this, we choose the points to be uniform in the variance-stabilizing coordinate *y* = 2 arcsin 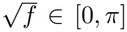 i.e., the cells are the intervals *f* ∈ [sin^2^((*i* − 1)*π/*2*N*), sin^2^(*iπ/*2*N*)] for 1 ≤ *i* ≤ *N*.

After computing *l*_*u*_(*θ, s*_inv_) for each cluster trajectory *u*, we define the log-likelihood for a species as

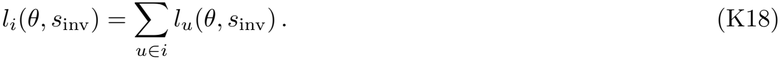

Note that there is no additional weight for each cluster, since their log-likelihoods are already scaled with the read counts through the measurement noise model. Now, we can integrate out *s*_inv_ to find the total log-likelihood

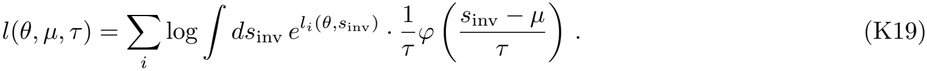

The Gaussian integral is computed in the same manner as in *p*_*A*_.

In the filtering algorithm, we obtain *p*_*f*_ by numerically solving the Fokker-Planck equation in Eq. (H3) on the grid of *f* with size *N*. Let *p*_*i*_(*t*) be the probability that *f* is within cell *i* at time *t*. Our goal is to convert Eq. (H3) into the form of master equation

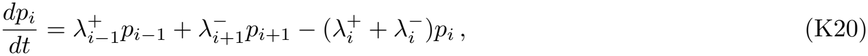

where 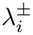 are transition rates to be determined. For the interior cells with 1 *< i < N*, the transition rates can be found by discretizing the derivatives in Eq. (H3). Let *f*_*i*_ = sin^2^((*i* − 1*/*2)*π/*2*N*) be the midpoint of cell *i* and define the cell distances 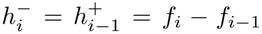 and 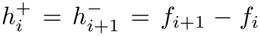. For second derivatives, we use the central approximation

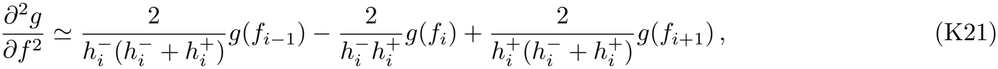

for some function *g*. For first derivatives, to ensure that the resulting transition rates are positive, we use the upwind approximation (instead of, e.g., the central approximation) [105]

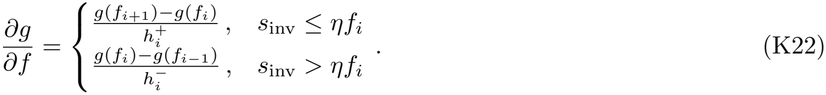

Substituting these derivatives into Eq. (H3) and comparing the coefficients, we then obtain the transition rates

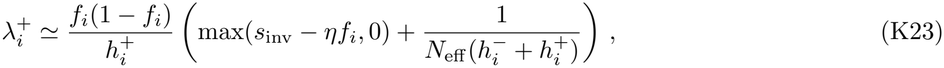

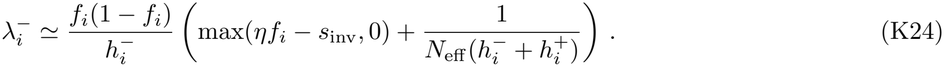

However, note again that the above derivation is only valid for the interior cells. For the two end cells, we impose absorbing boundary conditions and set

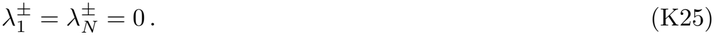

The above defines the full transition matrix on the grid of *f*. By exponentiating the transition matrix, we finally obtain *p*_*f*_ used in the likelihood function.

### 3. Maximum likelihood estimation

We now maximize the log-likelihood *l*(*θ, µ, τ*) for each period. We first perform grid search on the parameters *η, N*_eff_, *σ, µ, τ*, then refine the neighbor of grid points with maximum likelihood using optimization algorithms. We use the following grid:

- *η*∈ [−0.25, 0.75] in steps of 0.03125 for the disease and recovery periods; *η* ∈ [−0.25, 0.75] in steps of 0.0625, then [0.75, 3] in steps of 0.125 for the baseline period;
- 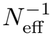∈ [5.6 *×* 10^*−*6^, 0.3] in 26 logarithmic steps;
- *σ* ∈ *{*0, 0.025, 0.05, 0.1, 0.15, 0.2, 0.25, 0.3, 0.4, 0.5, 0.6, 0.7, 0.8, 0.9, 1.05, 1.25, 1.5*}*;
- *µ* ∈ [−0.4, 1.2] in 80 uniform steps;
- *τ* ∈ [0.003, 1] in 40 logarithmic steps.

For integrating out *s*_inv_, we also compute *l*_*u*_(*θ, s*_inv_) with *s*_inv_∈ [−0.6, 1.6] in steps of 0.01. We use natural cubic splines to interpolate between grid points of *s*_inv_.

We refine the maximum likelihood from grid search as follows. First, for each given *η, N*_eff_, *σ*, we find the optimal *µ, τ* using Nelder-Mead method on the space (*µ*, log *τ*). This is computationally efficient since *l*_*u*_(*θ, s*_inv_) remains unchanged at every iteration of the method. Next, since *η* is the main focus of this analysis, we compute its marginal likelihood (see Fig. 5c in main text)

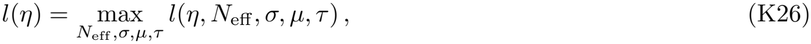

using the grid points of *N*_eff_, *σ* and optimized *µ, τ* . We then approximate the actual peak of *l*(*η*) by fitting a smoothing spline around the maximum points. The remaining parameters are then fitted at the optimal *η* with the above grid search, followed by Nelder-Mead method on the space (log *N*_eff_, *σ*).

### 4. Inference results and confidence

As shown in Fig. 5c in main text, *l*(*η*) has clear positive peaks significantly away from *η* = 0 for the disease and recovery periods. On the other hand, the peak in *l*(*η*) for the baseline period is not significantly different from *η* = 0. Moreover, *l*(*η*) for the baseline period reaches plateau instead of decaying for large *η*. Therefore, we cannot confidently identify the optimal *η* for the baseline period. Below we focus on the disease and recovery periods.

**TABLE S1:** Fitted model parameters for the human gut microbiome data. We fit the model of frequency-dependent selection to the human gut microbiome data [75] and infer the optimal parameters *η, N*_eff_, *σ, µ, τ* . Confidence intervals are obtained using bootstrapping on species. The inferred values indicate strong ecological feedback with parent-mutant coexistence in the community.

|  | Disease | Recovery |
|---|---|---|
| η | 0.174 | 0.078 |
| bootstrap 50% CI | [0.127, 0.196] | [0.058, 0.097] |
| bootstrap 95% CI | [0.048, 0.268] | [0.026, 0.137] |
| N<sub>eff</sub><sup>−1</sup> | 6.39 × 10<sup>−3</sup> | 3.93 × 10<sup>−3</sup> |
| bootstrap 95% CI | [2.73, 14.1] × 10<sup>−3</sup> | [1.64, 6.60] × 10<sup>−3</sup> |
| α | 5.21 | 4.44 |
| bootstrap 95% CI | [2.6, 7.9] | [2.4, 8.2] |
| σ | 0.473 | 0.760 |
| bootstrap 95% CI | [0, 0.69] | [0.51, 0.97] |
| μ | 0.097 | 0.040 |
| τ | 0.104 | 0.022 |
| P(0 < s<sub>inv</sub> < η) | 0.59 | 0.92 |
| expected coexistence | 10.2/15 | 15.9/17 |

**TABLE S2:** Variations of frequency-dependent selection under refitting on jackknife and on synthetic data. We check the robustness of the inferred *η* with two tests. First, we refit *η* on data with one species removed, i.e., the jackknife resampling. Second, we refit *η* on *B* = 200 set of synthetic data. The variations are well within the confidence intervals of *η*.

|  | Disease | Recovery |
|---|---|---|
| *η* | 0.174 | 0.078 |
| jackknife range | [0.148, 0.196] | [0.064, 0.095] |
| synthetic mean | 0.200 | 0.097 |
| synthetic 95% CI | [0.113, 0.318] | [0.058, 0.136] |

The optimal parameters are presented in Table S1. The confidence intervals are obtained from bootstrapping on species. That is, we produce *B* = 4000 communities of resampled species with replacement, then refit the optimal parameters. Compared with the original MLE procedure, the optimal *N*_eff_, *σ* are now also inferred using smoothing splines, so that the computational cost is controlled. This bootstrap approach is valid for the parameters *η, N*_eff_, *σ*, but not for the distribution parameters *µ, τ*, which are expected to depend on the species composition.

With the inferred distribution of *s*_inv_, we can estimate the expected fraction of species that satisfies 0 *< s*_inv_ *< η*, hence supports parent-mutant coexistence. A simple estimate for the fraction of coexistence is the probability *P*(0 *< s*_inv_ *< η*) under the inferred Gaussian distribution, see Table S1. However, we are more interested in the coexistence of the selected species, and the probabilities can be refined for each species using the trajectory data. Namely, we can compute the posterior distribution of *s*_inv_

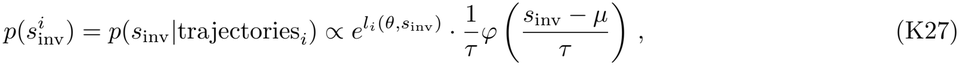

where *l*_*i*_(*θ, s*_inv_) has been computed in the MLE. Then, the expected number of species with coexistence is

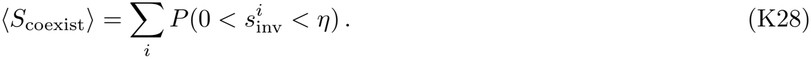

This number is presented in Table S1 and Fig. 5e in the main text.

We also perform a number of independent checks for the robustness of our results, as summarized in Tables S2 and S3. First, we apply jackknife resampling, i.e., refit *η* on data with one of the species removed. We go through all possible one-species removals and see little variation in *η* (second row in Table S2), indicating that the positive *η* is not driven by just one of the species. Second, we refit *η* by generating *B* = 200 set of synthetic data using the optimal parameters (see the detail of parametric bootstrap below). The refitted mean slightly deviates from the optimal *η*, but is still well within the confidence intervals (third and fourth rows in Table S2). The bias does not bring the intervals across *η* = 0 either. Therefore, our inference method has reliable level of accuracy. Finally, in Table S3, we show that our results are robust under different choices of the distribution of *s*_inv_.

**TABLE S3:** Variations of fit under different assumed distributions of *s*_inv_ across species. To check whether our results are sensitive to the Gaussian assumption on the distribution of invasion fitness *s*_inv_, we perform the same fit using several other two-parameter distributions. In both periods, the model parameters and the statistics of *s*_inv_ vary little across different choices of the distribution.

| Disease |  |  |  |  |  |  |
| *s*<sub>inv</sub> distribution | η | 10<sup>3</sup>*N*<sub>eff</sub><sup>−1</sup> | σ | *s*<sub>inv</sub> mean | *s*<sub>inv</sub> s.d. | coexistence |
|---|---|---|---|---|---|---|
| Gaussian | 0.174 | 6.39 | 0.47 | 0.097 | 0.104 | 10.2/15 |
| Logistic | 0.171 | 6.19 | 0.48 | 0.081 | 0.102 | 10.4/15 |
| Laplace | 0.169 | 5.96 | 0.49 | 0.066 | 0.104 | 10.5/15 |
| Uniform | 0.181 | 6.66 | 0.46 | 0.144 | 0.111 | 9.9/15 |
| Gumbel, right-skewed | 0.170 | 5.98 | 0.49 | 0.093 | 0.095 | 10.4/15 |
| Gumbel, left-skewed | 0.173 | 6.74 | 0.46 | 0.085 | 0.143 | 10.0/15 |
| Recovery |  |  |  |  |  |  |
| *s*<sub>inv</sub> distribution | η | 10<sup>3</sup>*N*<sub>eff</sub><sup>−1</sup> | σ | *s*<sub>inv</sub> mean | *s*<sub>inv</sub> s.d. | coexistence |
| Gaussian | 0.078 | 3.93 | 0.76 | 0.040 | 0.022 | 15.9/17 |
| Logistic | 0.074 | 3.79 | 0.76 | 0.039 | 0.023 | 15.5/17 |
| Laplace | 0.070 | 3.65 | 0.76 | 0.040 | 0.024 | 15.1/17 |
| Uniform | 0.083 | 4.13 | 0.76 | 0.040 | 0.021 | 17.0/17 |
| Gumbel, right-skewed | 0.085 | 4.27 | 0.76 | 0.044 | 0.025 | 16.4/17 |
| Gumbel, left-skewed | 0.069 | 3.60 | 0.76 | 0.036 | 0.022 | 15.5/17 |

### 5. Goodness of fit

Apart from the above confidence intervals, we can test the goodness of fit of our model by asking how well the distribution of 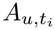 from our fitted model matches with the data distribution. Specifically, we focus on the goodness of one-step-ahead predictions for each time point. In the data, there may be more correlations across multiple days, such as memory effects beyond the current frequencies. Our model does not take these effects into account, and we do not measure these effects in the statistical test.

To assess the goodness of fit, there have been well-established statistical tests that compare data with continuous and common distributions, such as a uniform distribution. However, our data are discrete and follow distributions of arbitrary shape. To better utilize those statistical tests, we thus choose to use the randomized probability integral transform (PIT) to convert the data into new coordinates 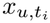, which we expect to follow a standard uniform distribution if our model is the true model underlying the data. We can therefore test our model by comparing the observed data in *x*-space with a standard uniform distribution using standard statistical tests.

The randomized PIT is performed as follows. Consider the cumulative distribution

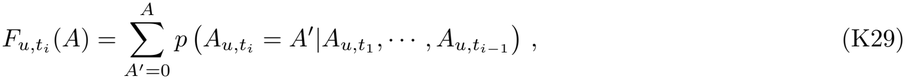

where the probabilities were already computed in the likelihood function. Since 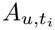 are integer data, the possible values of 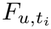are also discrete. To obtain a continuous coordinate, we further smooth these spikes in 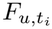by interpolation with uniform random noise. Define the variables

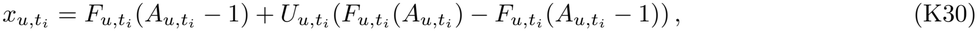

where 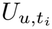 are independent standard uniform random variables. Intuitively, since observations with higher probabilities now occupy larger intervals in 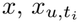 should follow a uniform distribution.

We collect a set of 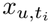 using 20 realizations of 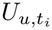 . To focus on one-step-ahead predictions, we split the set of 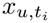 by the time points *t*_*i*_ and compute the test statistics for each *t*_*i*_. We compare 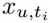 to standard uniform distribution using the Cramér-von Mises (CvM) statistics *ω*^2^ and the Anderson-Darling (AD) statistics *A*^2^.

We further calibrate the *p*-values for the tests using parametric bootstrap. Namely, we generate *B* = 200 sets of synthetic data using the fitted model, then refit the parameters and compute the test statistics, so that we obtain the null distributions for the tests. For the synthetic data, we use the fitted optimal *η, N*_eff_, *σ*, and the posterior mean for each 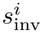 (see Eq. (K27)), so that the tests focus on the selected species. For consistency, the synthetic trajectories are drawn using the same distributions as in the likelihood computations. To keep the computational cost under control, we keep the same *N*_eff_, *σ* from the original model and only refit *η, µ, τ* for each set of synthetic data.

**TABLE S4:** Goodness of fit on one-step-ahead predictions. We perform Cramér-von Mises (CvM) and Anderson-Darling (AD) tests for our model of frequency-dependent selection at each predicted time point in the disease and recovery periods. The *p*-values shown are calibrated using parametric bootstrap. The first row shows the Bonferroni-corrected overall *p*-value for each period.

| Time | CvM | AD | Time | CvM | AD |
|---|---|---|---|---|---|
| Disease | 0.597 | 0.318 | Recovery | 0.149 | 0.149 |
| 46 | 0.149 | 0.080 | 80 | 0.741 | 0.701 |
| 49 | 0.363 | 0.512 | 92 | 0.443 | 0.274 |
| 67 | 0.726 | 0.617 | 110 | 0.219 | 0.184 |
| 69 | 0.428 | 0.368 | 118 | 0.025 | 0.025 |
|  |  |  | 129 | 0.378 | 0.308 |
|  |  |  | 153 | 0.443 | 0.398 |

The resulting *p*-values for each time point in the disease and recovery periods (except the first time points at which there are no predictions) are listed in Table S4. We also obtain an overall *p*-value per period using the Bonferroni correction:

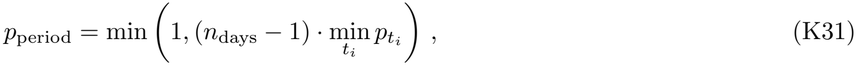

where *n*_days_ = 5, 7 are the number of days in each period. We see that there is no *p*-value rejecting our model except on day 118. The overall *p*-value cannot rule out that this rejection possibly occurs only by chance. Therefore, we conclude that our model is calibrated for one-step-ahead predictions, showing that frequency-dependent selection can indeed describe the data.

## References

[1] M. Kimura, On the probability of fixation of mutant genes in a population, Genetics 47, 713 (1962).

[2] M. Kimura, Theoretical foundation of population genetics at the molecular level, Theoretical population biology 2, 174 (1971).

[3] M. Kimura, Stochastic processes and distribution of gene frequencies under natural selection (1955), Population Genetics, Molecular Evolution, and the Neutral Theory: Selected Papers. The University of Chicago Press, Chicago 144 (1994).

[4] W. J. Ewens and W. Ewens, Mathematical population genetics: theoretical introduction, Vol. 27 (Springer, 2004).

[5] B. H. Good and O. Hallatschek, Effective models and the search for quantitative principles in microbial evolution, Current opinion in microbiology 45, 203 (2018).

[6] M. Möhle and S. Sagitov, A classification of coalescent processes for haploid exchangeable population models, Annals of Probability, 1547 (2001).

[7] B. H. Good, M. J. McDonald, J. E. Barrick, R. E. Lenski, and M. M. Desai, The dynamics of molecular evolution over 60,000 generations, Nature 551, 45 (2017).

[8] L. M. Wahl, P. J. Gerrish, and I. Saika-Voivod, Evaluating the impact of population bottlenecks in experimental evolution, Genetics 162, 961 (2002).

[9] J. F. Crow, An introduction to population genetics theory (Scientific Publishers, 2017).

[10] D. E. Dykhuizen and D. L. Hartl, Selection in chemostats, Microbiological reviews 47, 150 (1983).

[11] I. Cvijović, B. H. Good, E. R. Jerison, and M. M. Desai, Fate of a mutation in a fluctuating environment, Proceedings of the National Academy of Sciences 112, E5021 (2015).

[12] M. C. Whitlock, Fixation probability and time in sub-divided populations, Genetics 164, 767 (2003).

[13] O. Hallatschek and D. R. Nelson, Gene surfing in expanding populations, Theoretical population biology 73, 158 (2008).

[14] Z. Feng, E. Blumenthal, P. Mehta, and A. Goyal, A theory of ecological invasions and its implications for eco-evolutionary dynamics, ArXiv, arXiv (2025).

[15] J. McEnany and B. H. Good, Predicting the first steps of evolution in randomly assembled communities, Nature Communications 15, 8495 (2024).

[16] B. H. Good and L. B. Rosenfeld, Eco-evolutionary feed-backs in the human gut microbiome, Nature Communications 14, 7146 (2023).

[17] G. F. Fussmann, M. Loreau, and P. A. Abrams, Eco-evolutionary dynamics of communities and ecosystems, Functional ecology, 465 (2007).

[18] A. P. Hendry, Eco-evolutionary dynamics (Princeton university press, 2017).

[19] L. Govaert, E. A. Fronhofer, S. Lion, C. Eizaguirre, D. Bonte, M. Egas, A. P. Hendry, A. De Brito Martins, C. J. Melián, J. A. Raeymaekers, et al., Eco-evolutionary feedbacks—theoretical models and perspectives, Functional Ecology 33, 13 (2019).

[20] R. D. Holt, On the integration of community ecology and evolutionary biology: historical perspectives and current prospects (Elsevier, 2005).

[21] G. G. Mittelbach and D. W. Schemske, Ecological and evolutionary perspectives on community assembly, Trends in ecology & evolution 30, 241 (2015).

[22] L. Govaert, F. Altermatt, L. De Meester, M. A. Lei-bold, M. A. McPeek, J. H. Pantel, and M. C. Urban, Integrating fundamental processes to understand eco-evolutionary community dynamics and patterns, Functional Ecology 35, 2138 (2021).

[23] A. Robertson, Selection for heterozygotes in small populations, Genetics 47, 1291 (1962).

[24] T. Antal and I. Scheuring, Fixation of strategies for an evolutionary game in finite populations, Bulletin of mathematical biology 68, 1923 (2006).

[25] P. M. Altrock, C. S. Gokhale, and A. Traulsen, Stochastic slowdown in evolutionary processes, Physical Review E—Statistical, Nonlinear, and Soft Matter Physics 82, 011925 (2010).

[26] M. Assaf and M. Mobilia, Large fluctuations and fixation in evolutionary games, Journal of Statistical Mechanics: Theory and Experiment 2010, P09009 (2010).

[27] T. Connallon and A. G. Clark, A general population genetic framework for antagonistic selection that accounts for demography and recurrent mutation, Genetics 190, 1477 (2012).

[28] Z. J. Assaf, D. A. Petrov, and J. R. Blundell, Obstruction of adaptation in diploids by recessive, strongly deleterious alleles, Proceedings of the National Academy of Sciences 112, E2658 (2015).

[29] Z. D. Blount, C. Z. Borland, and R. E. Lenski, Historical contingency and the evolution of a key innovation in an experimental population of escherichia coli, Proceedings of the National Academy of Sciences 105, 7899 (2008).

[30] M. D. Herron and M. Doebeli, Parallel evolutionary dynamics of adaptive diversification in escherichia coli, PLoS biology 11, e1001490 (2013).

[31] E. M. Frenkel, M. J. McDonald, J. D. Van Dyken, K. Kosheleva, G. I. Lang, and M. M. Desai, Crowded growth leads to the spontaneous evolution of semistable coexistence in laboratory yeast populations, Proceedings of the National Academy of Sciences 112, 11306 (2015).

[32] P. A. Marquet, G. Espinoza, S. R. Abades, A. Ganz, and R. Rebolledo, On the proportional abundance of species: Integrating population genetics and community ecology, Scientific reports 7, 16815 (2017).

[33] I. Overcast, B. C. Emerson, and M. J. Hickerson, An integrated model of population genetics and community ecology, Journal of biogeography 46, 816 (2019).

[34] K. R. Foster, J. Schluter, K. Z. Coyte, and S. Rakoff-Nahoum, The evolution of the host microbiome as an ecosystem on a leash, Nature 548, 43 (2017).

[35] A. Ferreiro, N. Crook, A. J. Gasparrini, and G. Dantas, Multiscale evolutionary dynamics of host-associated microbiomes, Cell 172, 1216 (2018).

[36] D. Lawrence, F. Fiegna, V. Behrends, J. G. Bundy, A. B. Phillimore, T. Bell, and T. G. Barraclough, Species interactions alter evolutionary responses to a novel environment, PLoS biology 10, e1001330 (2012).

[37] V. Calcagno, P. Jarne, M. Loreau, N. Mouquet, and P. David, Diversity spurs diversification in ecological communities, Nature communications 8, 15810 (2017).

[38] S. Venkataram, H.-Y. Kuo, E. F. Hom, and S. Kryazhimskiy, Mutualism-enhancing mutations dominate early adaptation in a two-species microbial community, Nature Ecology & Evolution 7, 143 (2023).

[39] D. Schluter and M. W. Pennell, Speciation gradients and the distribution of biodiversity, Nature 546, 48 (2017).

[40] J. P. Hall, E. Harrison, and M. A. Brockhurst, Competitive species interactions constrain abiotic adaptation in a bacterial soil community, Evolution Letters 2, 580 (2018).

[41] T. Scheuerl, M. Hopkins, R. W. Nowell, D. W. Rivett, T. G. Barraclough, and T. Bell, Bacterial adaptation is constrained in complex communities, Nature communications 11, 754 (2020).

[42] M. Yamamichi, T. Gibbs, and J. M. Levine, Integrating eco-evolutionary dynamics and modern coexistence theory, Ecology Letters 25, 2091 (2022).

[43] M. Sireci and M. A. Muñoz, Statistical mechanics of phenotypic eco-evolution: From adaptive dynamics to complex diversification, Physical Review Research 6, 023070 (2024).

[44] M. Scheffer and E. H. Van Nes, Self-organized similarity, the evolutionary emergence of groups of similar species, Proceedings of the National Academy of Sciences 103, 6230 (2006).

[45] M. Doebeli and I. Ispolatov, Complexity and diversity, Science 328, 494 (2010).

[46] D. Zinder, T. Bedford, S. Gupta, and M. Pascual, The roles of competition and mutation in shaping antigenic and genetic diversity in influenza, PLoS pathogens 9, e1003104 (2013).

[47] T. Azarian, P. P. Martinez, B. J. Arnold, X. Qiu, L. R. Grant, J. Corander, C. Fraser, N. J. Croucher, L. L. Hammitt, R. Reid, et al., Frequency-dependent selection can forecast evolution in streptococcus pneumoniae, PLoS biology 18, e3000878 (2020).

[48] Q. He, S. Pilosof, K. E. Tiedje, S. Ruybal-Pesántez, Y. Artzy-Randrup, E. B. Baskerville, K. P. Day, and M. Pascual, Networks of genetic similarity reveal non-neutral processes shape strain structure in plasmodium falciparum, Nature communications 9, 1817 (2018).

[49] J. Liu and Q. He, An eco-evolutionary framework to predict long-term strain diversity and turnover in infectious diseases, bioRxiv, 2025 (2025).

[50] B. H. Good, S. Martis, and O. Hallatschek, Adaptation limits ecological diversification and promotes ecological tinkering during the competition for substitutable re-sources, Proceedings of the National Academy of Sciences 115, E10407 (2018).

[51] L. Fant, I. Macocco, and J. Grilli, Eco-evolutionary dynamics lead to functionally robust and redundant communities, bioRxiv, 2021 (2021).

[52] F. Roy, G. Biroli, G. Bunin, and C. Cammarota, Numerical implementation of dynamical mean field theory for disordered systems: Application to the lotka–volterra model of ecosystems, Journal of Physics A: Mathematical and Theoretical 52, 484001 (2019).

[53] M. T. Pearce, A. Agarwala, and D. S. Fisher, Stabilization of extensive fine-scale diversity by ecologically driven spatiotemporal chaos, Proceedings of the National Academy of Sciences 117, 14572 (2020).

[54] G. Bunin, Ecological communities with lotka-volterra dynamics, Physical Review E 95, 042414 (2017).

[55] R. M. May, Will a large complex system be stable?, Nature 238, 413 (1972).

[56] A. Mahadevan, M. T. Pearce, and D. S. Fisher, Spatiotemporal ecological chaos enables gradual evolutionary diversification without niches or tradeoffs, Elife 12, e82734 (2023).

[57] M. Advani, G. Bunin, and P. Mehta, Statistical physics of community ecology: a cavity solution to macarthur’s consumer resource model, Journal of Statistical Mechanics: Theory and Experiment 2018, 033406 (2018).

[58] A. Altieri, F. Roy, C. Cammarota, and G. Biroli, Properties of equilibria and glassy phases of the random lotka-volterra model with demographic noise, Physical Review Letters 126, 258301 (2021).

[59] A. Mahadevan and D. S. Fisher, Continual evolution in nonreciprocal ecological models, PRX Life 3, 033008 (2025).

[60] A. Goyal, J. W. Rocks, and P. Mehta, Universal niche geometry governs the response of ecosystems to environmental perturbations, PRX Life 3, 013010 (2025).

[61] M. Kimura and T. Ohta, Theoretical aspects of population genetics, 4 (Princeton University Press, 1971).

[62] C. K. Fisher and P. Mehta, The transition between the niche and neutral regimes in ecology, Proceedings of the National Academy of Sciences 111, 13111 (2014).

[63] M. Barbier and J.-F. Arnoldi, The cavity method for community ecology, bioRxiv, 147728 (2017).

[64] W. Cui, R. Marsland III, and P. Mehta, Les houches lectures on community ecology: From niche theory to statistical mechanics, ArXiv, arXiv (2024).

[65] S. Azaele and A. Maritan, Generalized dynamical mean field theory for non-gaussian interactions, Physical Review Letters 133, 127401 (2024).

[66] J. H. Gillespie, Some properties of finite populations experiencing strong selection and weak mutation, The American Naturalist 121, 691 (1983).

[67] H. Sompolinsky, Statistical mechanics of neural networks, Physics Today 41, 70 (1988).

[68] R. Levins, Theory of fitness in a heterogeneous environment. i. the fitness set and adaptive function, The American Naturalist 96, 361 (1962).

[69] R. Macarthur and R. Levins, The limiting similarity, convergence, and divergence of coexisting species, The American Naturalist 101, 377 (1967).

[70] R. Marsland III, W. Cui, J. Goldford, A. Sanchez, K. Korolev, and P. Mehta, Available energy fluxes drive a transition in the diversity, stability, and functional structure of microbial communities, PLoS computational biology 15, e1006793 (2019).

[71] M. Nei and A. Roychoudhury, Probability of fixation and mean fixation time of an overdominant mutation, Genetics 74, 371 (1973).

[72] F. Ruzicka, M. K. Zwoinska, D. Goedert, H. Kokko, X.-Y. Li Richter, I. R. Moodie, S. Nilén, C. Olito, E. I. Svensson, P. Czuppon, et al., A century of theories of balancing selection, Biological Reviews 101, 804 (2026).

[73] W. Cui, R. Marsland III, and P. Mehta, Effect of resource dynamics on species packing in diverse ecosystems, Physical review letters 125, 048101 (2020).

[74] M. Kimura and T. Ohta, The average number of generations until fixation of a mutant gene in a finite population, Genetics 61, 763 (1969).

[75] M. Roodgar, B. H. Good, N. R. Garud, S. Martis, M. Avula, W. Zhou, S. M. Lancaster, H. Lee, A. Babveyh, S. Nesamoney, et al., Longitudinal linked-read sequencing reveals ecological and evolutionary responses of a human gut microbiome during antibiotic treatment, Genome research 31, 1433 (2021).

[76] R. Wolff, W. Shoemaker, and N. Garud, Ecological stability emerges at the level of strains in the human gut microbiome, MBio 14, e02502 (2023).

[77] N. R. Garud, B. H. Good, O. Hallatschek, and K. S. Pollard, Evolutionary dynamics of bacteria in the gut microbiome within and across hosts, PLoS biology 17, e3000102 (2019).

[78] S. Zhao, T. D. Lieberman, M. Poyet, K. M. Kauffman, S. M. Gibbons, M. Groussin, R. J. Xavier, and E. J. Alm, Adaptive evolution within gut microbiomes of healthy people, Cell host & microbe 25, 656 (2019).

[79] A. Goyal, L. S. Bittleston, G. E. Leventhal, L. Lu, and O. X. Cordero, Interactions between strains govern the eco-evolutionary dynamics of microbial communities, Elife 11, e74987 (2022).

[80] J. A. Ascensao, K. D. Abedi, A. N. Prasad, and O. Hallatschek, Frequency-dependent fitness effects are ubiquitous, bioRxiv (2025).

[81] A. Goyal, V. Dubinkina, and S. Maslov, Multiple stable states in microbial communities explained by the stable marriage problem, The ISME journal 12, 2823 (2018).

[82] W. Lopes, D. R. Amor, and J. Gore, Cooperative growth in microbial communities is a driver of multistability, Nature Communications 15, 4709 (2024).

[83] E. Blumenthal, J. W. Rocks, and P. Mehta, Phase transition to chaos in complex ecosystems with nonreciprocal species-resource interactions, Physical review letters 132, 127401 (2024).

[84] T. Arnoulx de Pirey and G. Bunin, Many-species eco-logical fluctuations as a jump process from the brink of extinction, Physical Review X 14, 011037 (2024).

[85] J. Hu, D. R. Amor, M. Barbier, G. Bunin, and J. Gore, Emergent phases of ecological diversity and dynamics mapped in microcosms, Science 378, 85 (2022).

[86] M. M. Desai and D. S. Fisher, Beneficial mutation– selection balance and the effect of linkage on positive selection, Genetics 176, 1759 (2007).

[87] T. Yoshida, S. P. Ellner, L. E. Jones, B. J. M. Bohannan, R. E. Lenski, and N. G. Hairston Jr, Cryptic population dynamics: rapid evolution masks trophic interactions, PLoS biology 5, e235 (2007).

[88] S. Liu and P. Mehta, Ecology of metagenomes: incorporating genotype-to-phenotype maps into ecological models, bioRxiv (2026).

[89] G. Kinsler, K. Geiler-Samerotte, and D. A. Petrov, Fitness variation across subtle environmental perturbations reveals local modularity and global pleiotropy of adaptation, Elife 9, e61271 (2020).

[90] D. S. Maynard, C. A. Serván, and S. Allesina, Network spandrels reflect ecological assembly, Ecology letters 21, 324 (2018).

[91] S. Pearl Mizrahi, H. Lee, A. Goyal, E. Owen, and J. Gore, Structured interactions explain the absence of keystone species in synthetic microcosms, The ISME Journal 19, wraf211 (2025).

[92] S. F. M. Hart, J. M. B. Pineda, C.-C. Chen, R. Green, and W. Shou, Disentangling strictly self-serving mutations from win-win mutations in a mutualistic microbial community, Elife 8, e44812 (2019).

[93] J. A. Ascensao, K. M. Wetmore, B. H. Good, A. P. Arkin, and O. Hallatschek, Quantifying the local adaptive landscape of a nascent bacterial community, Nature Communications 14, 248 (2023).

[94] I. Dornic, H. Chaté, and M. A. Munoz, Integration of langevin equations with multiplicative noise and the vi-ability of field theories for absorbing phase transitions, Physical review letters 94, 100601 (2005).

[95] G. Garcia Lorenzana, A. Altieri, and G. Biroli, Interactions and migration rescuing ecological diversity, PRX Life 2, 013014 (2024).

[96] W. Cui, J. W. Rocks, and P. Mehta, An elementary mean-field approach to the spectral densities of random matrix ensembles, Physica A: Statistical Mechanics and its Applications 637, 129608 (2024).

[97] D. S. Fisher, Course 11 evolutionary dynamics, Les Houches 85, 395 (2007).

[98] B. H. Good, Molecular Evolution in Rapidly Evolving Populations, Ph.D. thesis, Harvard University, Graduate School of Arts and Sciences (2016).

[99] P. Hänggi, P. Talkner, and M. Borkovec, Reaction-rate theory: fifty years after kramers, Reviews of modern physics 62, 251 (1990).

[100] C. M. Bender and S. A. Orszag, Advanced mathematical methods for scientists and engineers: Asymptotic methods and perturbation theory, Vol. 1 (Springer, 1999).

[101] P. W. Glynn, Likelihood ratio gradient estimation for stochastic systems, Communications of the ACM 33, 75 (1990).

[102] P. B. Warren and R. J. Allen, Malliavin weight sampling for computing sensitivity coefficients in brownian dynamics simulations, Physical review letters 109, 250601 (2012).

[103] S. F. Levy, J. R. Blundell, S. Venkataram, D. A. Petrov, D. S. Fisher, and G. Sherlock, Quantitative evolutionary dynamics using high-resolution lineage tracking, Nature 519, 181 (2015).

[104] A. N. Nguyen Ba, I. Cvijović, J. I. Rojas Echenique, K. R. Lawrence, A. Rego-Costa, X. Liu, S. F. Levy, and M. M. Desai, High-resolution lineage tracking reveals travelling wave of adaptation in laboratory yeast, Nature 575, 494 (2019).

[105] H. J. Kushner, Numerical methods for stochastic control problems in continuous time, SIAM Journal on Control and Optimization 28, 999 (1990).

